# Unexpected population structure and plant-pathogen interaction as revealed by very large-scale molecular and biological analyses of *Leptosphaeria maculans* avirulence genes in France

**DOI:** 10.64898/2026.11.27.769512

**Authors:** Angélique Gautier, Audren Jiquel, Marie-Hélène Balesdent, Valérie Laval

## Abstract

**BACKGROUND:** Rational management of rapeseed *(Brassica napus)* varieties resistant to the fungus *Leptosphaeria maculans* requires regular estimates of virulent isolate proportions in field populations. This paper describes a large-scale survey of French *L. maculans* populations conducted in 2022-23 at 11 sites. Isolate Phenotyping focused on avirulence genes *AvrLm6, AvrLm11*, and *AvrLmS-Lep2*, corresponding to effective resistance genes, and *AvrLm4* as a virulence control. In parallel, the molecular tool MPSeqM was applied to 44 pools of symptomatic leaves (1402 leaf spots) to estimate proportions of virulent isolates for 10 *AvrLm/Rlm* interactions.

**RESULTS:** Phenotyping of 716 isolates on a plant differential set revealed that *L. maculans* populations from 10 sites were similar across sites and to those from the 2019-20 sampling, with 97% virulent isolates on *Rlm4*, 0% on *Rlm6*, 0.5% on *RlmS-Lep2*, and 3.7% on *Rlm11*. The last site, located in Bourgogne and still never analysed, differed from the others, with 53.2%, 64.6%, and 21.5% of virulent isolates on *Rlm4, Rlm6*, and *Rlm11*, respectively. MPSeqM also identified this site as distinct from the others for both allelic variation and virulence proportions, including genes not assessed by phenotyping, such as *AvrLm5*, with 35.8% of virulent isolates compared to 0% elsewhere. At this site, phenotyping and molecular analyses diverged for *AvrLm11*, suggesting an atypical expression of the *AvrLm11/Rlm11* interaction.

**CONCLUSION:** This study confirmed the potential of the MPSeqM tool for large-scale surveys and for rapid identification of atypical populations or interactions. It led us to propose a schedule and sampling scheme for further surveys.

## 1 INTRODUCTION

The fungal pathogen *Leptosphaeria maculans* (syn. *Plenodomus lingam)* is one of the pRlmary threats to oilseed rape (rapeseed) cultivation in most parts of the world^1^. The disease is characterized by the development of a basal stem, leading to plant lodging before harvest and seed loss. The epidemic begins early in the growing season when the fungus releases ascospores, infecting young leaves and causing small symptoms (leaf spots) before it systemically and symptomlessly colonizes petioles and stems until stem canker develops^2,3^. When climatic conditions are favourable, the fungus surviving on residues may continue releasing ascospores for many weeks or months^3^. Consequently, fungicide treatments offer only limited efficiency against this pathogen. The main control strategy currently deployed in most parts of the world is breeding for resistance or tolerant oilseed rape varieties combined with a reasoned deployment of these varieties.

Resistance of oilseed rape to *L. maculans* combines two types of resistance: polygenic, partial resistance and monogenic, specific resistance^4^. Partial resistance limits the progression of the fungus from leaf spots to the stem base and/or limits the stem canker severity. Specific resistance stops the fungal development at the leaf spot stage following the gene-for-gene model; plants carrying a specific resistance gene *(Rlm* for Resistance *to L. maculans)* will induce their defence mechanisms and block the progression of any *L. maculans* isolate possessing an avirulent allele of the corresponding avirulence *(AvrLm)* gene^1,5^. Specific resistance is therefore highly efficient for disease control, but it targets only a part of the pathogenic population. In addition, it exerts strong selection pressure on pathogenic populations, causing resistance breakdown due to the selection and spread of isolates lacking the avirulence gene or possessing a virulent allele of the gene^6^. This is particularly true for resistance toward *L. maculans* as this fungus displays a very high capacity to generate new alleles, including gene deletion^7^, and all biological traits favouring rapid evolution (i.e., large population size coupled with both asexual and sexual reproduction)^8^. The gene-for-gene relationship between *B. napus* and *L. maculans* has been well studied, with 10 avirulence genes cloned to date^1^. Based on this knowledge, many countries have developed pathogen survey campaigns to choose effective *Rlm* genes and improve their durability. For instance, large-scale population surveys were performed in Germany, Canada, and Australia, leading to advice for the deployment of oilseed rape varieties with specific and effective resistance genes^9,10,11,12^. In France, *L. maculans* populations have been monitored at regular time intervals since the beginning of the century^13,14,15^.They have highlighted putative effective *Rlm* genes, i.e., corresponding to *AvrLm* genes with avirulent alleles present in 100% of the population (for instance, *Rlm7* in 2000). They also detected and characterized successive *Rlm* breakdowns, such as the rapid overwhelming of *Rlm1* or the much slower and still partial *Rlm7* breakdown^13,14^. Such surveys were also helpful to identify atypical gene-for-gene interactions. For instance, large-scale surveys evidenced a negative correlation between avirulence on *Rlm3* and *Rlm7;* while 100% of the population was avirulent on *Rlm7* and virulent on *Rlm3* in the early 2000’s, all virulent isolates toward *Rlm7* isolated in the course of *Rlm7* breakdown appeared to be avirulent on *Rlm3*^16^. This observation was the starting point of molecular studies that deciphered this complex interaction and demonstrated that the *AvrLm7* gene, or at least some of its avirulence alleles, could mask the *AvrLm3-Rlm3* recognition^17^.

*L. maculans* population surveys initially relied on *in vitro* isolation and cultivation of field isolates, collected in different regions, followed by phenotyping of each isolate by inoculation on a plant differential set comprising defined *Rlm* genes^13^. Although accurate, this method is time-consuming and needs specific growth chambers with controlled conditions, allowing a correct expression of resistance or susceptibility symptoms. In France, the last *L. maculans* large-scale survey was conducted on a 2019-20 sampling consisting of 844 isolates from 10 locations^15^. This sampling was used to develop in parallel a molecular method aiming at reducing the time and costs of such surveys^18^. Briefly, this method (“MPSeqM” tool) bypassed the isolation and inoculation stages. Instead, 32 randomly-taken leaf spots were pooled as one sample, subsequently analysed after DNA extraction, multiplex PCR amplification of eight genes, and HTS (High Throughput Sequencing) of the PCR products using MiSeq technology. After mapping the reads on available databases of *AvrLm* allele sequences^7^, the percentages of each allele, and consequently the proportion of virulent or avirulent alleles, were calculated for each pooled sample, and then for each sampled site or at the country scale. This proof-of-concept work demonstrated the feasibility and accuracy of the method, with a high correlation between phenotyping results by the standard method and sequencing results using the MPSeqM tool. It offers the possibility to characterize, in a single sequencing run, up to 150 pools of 32 leaf spots, corresponding to nearly 50 sampling sites with a total of more than 4500 leaf spots^18^.

In this paper, we monitored French *L. maculans* populations over a new growing season using both methods and including sites that had been rarely or never analysed before. The main aim was to determine the dominant alleles of avirulence genes of interest, compared to the last survey, and to investigate a potential regional effect on these allele frequencies.

This study confirmed the MPSeqM potential while clarifying its limitations. Above all, it highlighted one specific expeRlmental site that differed from the others in several characteristics. Finally, the combination of biological and molecular approaches suggests a new complex interaction involving *AvrLm11*.

## 2 MATERIAL AND METHODS

### 2.1 Reference Isolates

Seventy-one characterized *L. maculans* isolates were chosen to prepare DNA controls in MPSeqM expeRlments, including the reference isolate JN3 (v23.1.3)^7^ (Supporting information Table SI). Five isolates (JN2, X82.14, INV13.269 and X83.51) were used as controls in pathotyping expeRlments^5,18^.

### 2.2 Field samples

Two cropping seasons, 2019-20 and 2022-23, were compared here. The 2019 sampling was previously described and comprised 844 isolates from a trap cultivar sown in 10 geographical locations^15,18^ (Fig. 1). A similar sampling was performed in autumn 2022 in 11 sites (Table 1). Individual leaves with typical *L. maculans* leaf spots were collected (up to 300 leaves per site, all collected from distinct plants). Part of these leaves were used to isolate single-pycnidium isolates (between 20 and 80 per site, mean=65, total number 716 isolates) for pathotyping on a differential plant set^18^. In parallel, 1cm diameter leaf disks (one per leaf) centred on a single leaf spot were cut and stored at -20°C. Before DNA extraction, leaf spots from a given site and plot were pooled in one Falcon tube, mostly by 32 (three to six pools per site and per sampling campaign, Supporting Information Table S2) and freeze-dried for 24 hours. When typical *L. maculans* leaf spots were seen on cultivars with effective resistance genes *(Rlm6, Rlm11* and *RlmS-Lep2)*, isolates were recovered from these leaf spots in order to gather a collection of virulent isolates.

**Table 1.** Percentages of virulent *Leptosphaeria maculans* isolates toward nine *Brassica napus* resistance genes in sites sampled in 2019-2020 or 2022-2023, based on phenotyping (A) or using the MPSeqM tool (B)

| Year of isolation | French department | Location | Percentage of virulent isolates toward: |  |  |  |  |  |  |  |  |  |  |  |  |  |  |  |  |  |  |  |
|---|---|---|---|---|---|---|---|---|---|---|---|---|---|---|---|---|---|---|---|---|---|---|
|  |  |  | *Rlm1* |  | *Rlm2* |  | *Rlm3* |  | *Rlm4* |  | *Rlm5* |  | *Rlm6* |  | *Rlm7* |  | *Rlm9* |  | *LepR2/RlmS* |  | *Rlm11* |  |
|  |  |  | A | B | A | B | A | B | A | B | A | B | A | B | A | B | A | B | A | B | A | B |
| 2019 | Eure-et-Loir (28)<sup>†</sup> | Janville | 100 | 99.4 | 100 | 98.8 | 37.5 | 46.9 | 100 | 98 | nd‡ | 0 | 0 | 0 | 87.5 | 72.1 | 100 | 99.5 | 0 | 0 | 3.4 | 0.2 |
|  | Ille-et-Vilaine (35) | Le Rheu | 95.3 | 88.2 | 100 | 99.4 | 68.3 | 64.3 | 97.5 | 95.7 | nd | 0 | 0 | 0 | 32.9 | 35.3 | 100 | 99.5 | 0 | 0 | 6.9 | 0 |
|  | Indre (36) | Bouges-le-Chateau | 100 | 99.8 | 100 | 99.4 | 38.1 | 44.9 | 100 | 99 | nd | 0 | 0 | 0 | 79.8 | 66.5 | 100 | 99.5 | 0 | 0 | 3.3 | 0 |
|  | Indre (36) | Lange | 100 | nd | 100 | nd | 31.9 | nd | 100 | nd | nd | nd | 0 | nd | 84.7 | nd | 100 | nd | 1.4 | nd | 3.3 | nd |
|  | Loiret (45) | Coullons | 100 | 99.9 | 100 | 98.7 | 53 | 48.5 | 100 | 98.5 | nd | 0 | 0 | 0 | 74.7 | 60.6 | 100 | 99.5 | 0 | 0 | 3.3 | 1.9 |
|  | Loir-et-Cher (41) | Boisseau | 100 | 99.1 | 100 | 99.4 | 51.2 | 52.7 | 100 | 98.1 | nd | 0 | 0 | 0 | 91.2 | 72.7 | 100 | 99.6 | 0 | 0.7 | 3.3 | 1 |
|  | Maine-et-Loire (49) | Erdre-en-Anjou | 96.8 | 99 | 100 | 99.9 | 53.7 | 51.5 | 100 | 96.7 | nd | 0 | 0 | 0 | 51.2 | 49.7 | 100 | 98.9 | 0 | 0 | 3.3 | 2.9 |
|  | Marne (51) | Champigneul-Champagne | 96.7 | nd | 100 | nd | 62.2 | nd | 100 | nd | nd | nd | 0 | nd | 65.8 | nd | 100 | nd | 0 | nd | 6.9 | nd |
|  | Nord (59) | Frelinghien | 96.8 | 99.9 | 100 | 100 | 48.7 | 58.9 | 97.4 | 89.8 | nd | 0 | 0 | 0 | 60 | 53.7 | 100 | 99.4 | 0 | 0 | 0 | 0 |
|  | Yvelines (78) | Thiverval-Grignon | 100 | 99.3 | 100 | 99.5 | 38.4 | 51.1 | 100 | 94.9 | nd | 0 | 0 | 0 | 78.5 | 60.3 | 100 | 99.5 | 0 | 0.2 | 0 | 0.4 |
| 2022 | Côte-d'Or (21) | Bretenière | nd | 48.3 | nd | 99.6 | 97.4 | 80 | 53.2 | 61.5 | nd | 35.8 | 64.6 | 41.1 | 19 | 22.3 | nd | 100 | 1.3 | 0 | 21.5 | 0 |
|  | Eure-et-Loir (28) | Terminiers | nd | 100 | nd | 100 | nd | 69.6 | 100 | 91.8 | nd | 0 | 0 | 0 | nd | 59.5 | nd | 100 | 1.3 | 0.3 | 2.5 | 0.3 |
|  | Eure-et-Loir (28) | Toury | nd | 98.6 | nd | 99.3 | nd | 65.7 | 98.7 | 95.7 | nd | 0 | 0 | 0 | nd | 65.6 | nd | 100 | 0 | 0 | 3.9 | 0.3 |
|  | Haute Garonne (31) | Aucamville | nd | nd | nd | nd | nd | nd | 100 | nd | nd | nd | 0 | nd | nd | nd | nd | nd | 2.2 | nd | 2.2 | nd |
|  | Ille-et-Vilaine (35) | Le Rheu | nd | 84.6 | nd | 99.1 | nd | 75.1 | 96.1 | 95.5 | nd | 0 | 0 | 0 | nd | 37.6 | nd | 99.5 | 1.3 | 0.4 | 6.5 | 0.8 |
|  | Loir-et-Cher (41) | Villeneuve-Frouville | nd | 98.2 | nd | 100 | nd | 64.6 | 100 | 93.5 | nd | 0 | 0 | 0 | nd | 65.7 | nd | 100 | 0 | 0 | 7.7 | 0 |
|  | Nord (59) | Allènes-les-marais | nd | nd | nd | nd | nd | nd | 95 | nd | nd | nd | 0 | nd | nd | nd | nd | nd | 0 | nd | 5 | nd |
|  | Nord (59) | Ennevelin | nd | 99.8 | nd | 97.7 | nd | 73.3 | 95.9 | 92.6 | nd | 0 | 0 | 0 | nd | 48.9 | nd | 97 | 0 | 0.2 | 2.7 | 5.1 |
|  | Pas de Calais (62) | Saint-Pol-sur-Ternoise | nd | 100 | nd | 98.7 | nd | 71.7 | 90.9 | 90 | nd | 0 | 0 | 0.9 | nd | 41.2 | nd | 100 | 0 | 0 | 3.9 | 3.1 |
|  | Seine-et-Marne (77) | Verneuil-l'étang | nd | nd | nd | nd | nd | nd | 96.7 | nd | nd | nd | 0 | nd | nd | nd | nd | nd | 0 | nd | 0 | nd |
|  | Yvelines (78) | Versailles | nd | 99.8 | nd | 99.6 | nd | 77 | 97.5 | 94.2 | nd | 0 | 0 | 0 | nd | 49.6 | nd | 100 | 0 | 0.8 | 2.5 | 0.5 |

† refers to the french department number
‡ nd. not done

**Figure 1.**
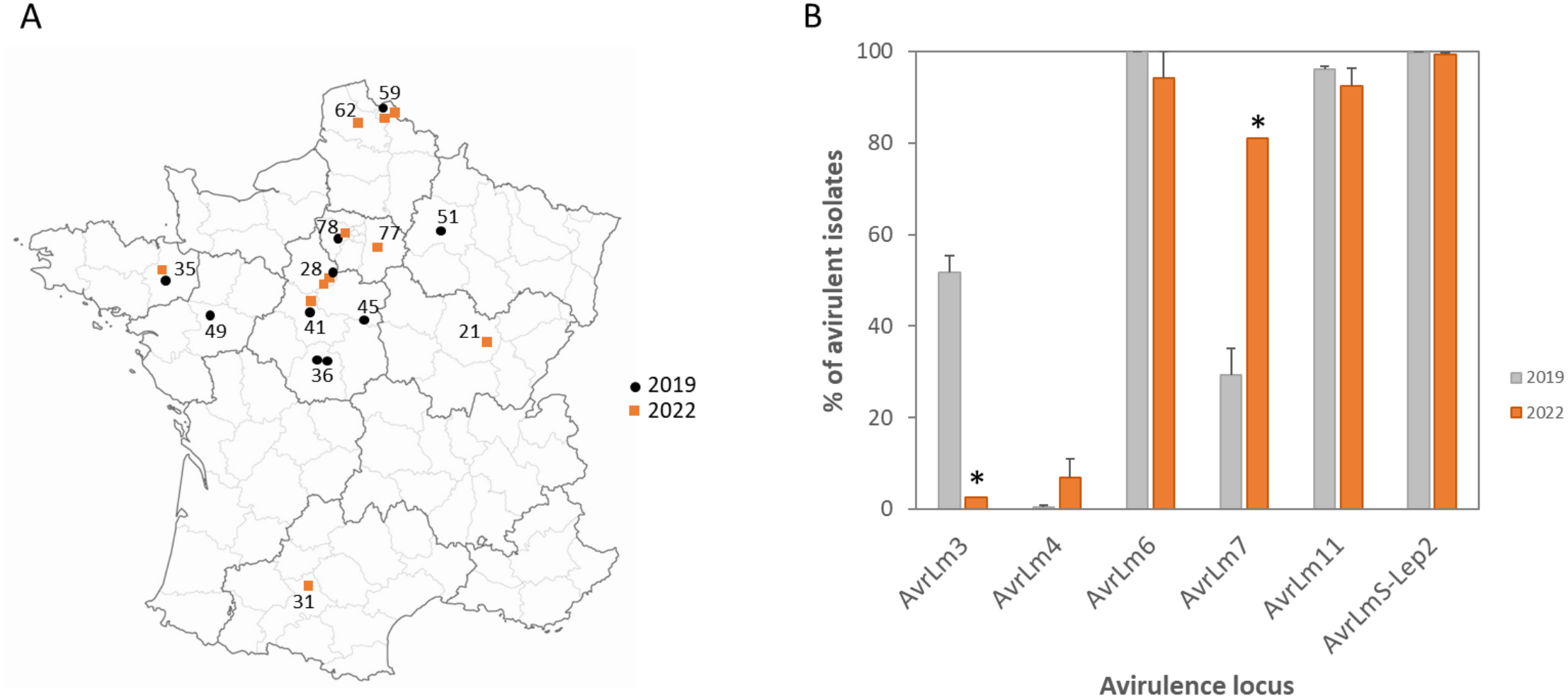
The geographical distribution of the sampled oilseed rape fields (sites) and the prevalence of avirulence alleles of Leptosphaeria maculans in France. (A) The numbers on the map indicate the French department number, with each dot representing a sampled *Brassica napus* field. (B) For each avirulence gene analyzed, the histogram displays the mean (and standard error) of frequencies of avirulent isolates. In 2019, six genes were analyzed on isolates from 10 sites (n=844 isolates). The asterisk indicates the specificity of AvrLm3 and AvrLm7 genes, in 2022, which were analyzed on isolates from one site (21), while the other genes were analyzed on isolates from 11 sites (n=716 isolates). Licence : CC BY-NC-SA, comersis.com, https://france.comersis.com/carte-france-gratuite.php?f=3 Carte:https://umap.openstreetmap.fr/fr/map/anonymous-edit/1280269:4Mc6TBbSCpL1srhvDqLAoLe4SCjeVxFPF_0_mtz_F8E

### 2.3 Plant inoculation for preparation of DNA control

A selection of 32 isolates with known *AvrLm* alleles was inoculated on cotyledons of *B. napus* cv. Westar (no *Rlm* gene) using standard protocols^17^ (Supporting information Table SI). Leaf spots developed on cotyledons were cut as described above. One spot per isolate was placed in a 50 mL Falcon tube, to constitute a pool of 32 leaf spots of known *AvrLm* allele composition (MIX_T).

### 2.4 Plant inoculation for isolate pathotyping

Isolates were inoculated on a *B. napus* differential set using standard protocols^14^. For the 2019-20 sampling, the set comprised cultivars Cooper *(Rlm1, Rlm4)*, Bristol *(Rlm2, Rlm9), 22*1-1 *(Rlm3)*, Darmor-MX *(Rlm6)*, 23-1-1 *(Rlm7)*, Darmor-R/m11 *(Rlm11)*, and Topas-lep/R2 *(LepR2/RlmS)*, allowing a complete phenotypic characterization of *AvrLm1, AvrLm2, AvrLm3, AvrLm4, AvrLm6, AvrLm7, AvrLm9, AvrLm11* and *AvrLmS-Lep2*^5,13,19,20^. The differential set for the 2022-23 survey was adapted to only characterize avirulence genes that were in the majority avirulent in 2019-20. It comprised Pixel *(Rlm4)* as a susceptible control, Darmor-MX, Darmor-R/m11, and Topas-lep/R2. Additional tests were performed to characterize *AvrLm3* and *AvrLm7* in the Bretenière site, as these populations were never characterized to date.

### 2.5 DNA extraction

Fungal DNA was extracted from 1 mL of spore suspensions (between 10^8^ and 10^9^ spores mL^-1^)^7^. Freeze-dried leaf spots were crushed with a tungsten carbide bead (3 mm, Qiagen) on a Retsch MM400 mixer mill for 60 seconds at 30 Hz. DNA was extracted using the DNeasy Plant Mini kit (Qiagen) and eluted in 100 µL of AE buffer (Qiagen). DNA quality and concentration were assessed with NanoDrop™ One (Thermo Scientific™) and Qubit™ 2.0 fluoRlmeter (Invitrogen™), respectively.

### 2.6 Control DNA samples

Four controls were included in the sequencing run: (i), JN3 DNA (positive control); (ii), the pool of 32 leaf spots from inoculations with known isolates (Hi), the pool of 71 *L. maculans* DNA; (iv), four DNA field samples from the 2019-20 survey^7^. Eleven points of dilution series for deletion quantification^7^ were prepared by mixing DNA of isolates JN3 and X83.51, or JN3 and IBCN18, in proportions varying from 0 % to 100% (with a 10% increment)^7^. X83.51 or IBCN18 alone in the dilution series served as negative controls for deleted genes (Supporting information Table SI).

### 2.7 Multiplex PCR and high-throughput sequencing

The first MPSeqM run (RunA, 2019-20 sampling) was performed using multiplex PCR and MiSeq sequencing as described previously^18^. For the 2022-23 sampling a second run (RunB) followed the same procedure. PRlmers, PCR mixes and conditions were identical for the two sampling years^18^, except for *AvrLm6* pRlmers (Supporting information Table S3). Each DNA sample was analysed in triplicate, i.e., following three independent PCRs and sequencing. PCR products were sent to the GenoToul platform (INRAE Toulouse) for sequencing. Samples were pooled in one library and sequenced on a single flow cell of the high-throughput Illumina MiSeq (Illumina, Inc., San Diego, CA, USA) cartridge according to manufacturer’s instructions. Sequencing was performed using a paired-end method (2×250 bp paired-end reads and single 6-bp index reads) with the MiSeq V3 Reagent Kit (Illumina).

### 2.8 Single avirulence genes amplification and sequencing

Full-length single avirulence genes were amplified in some individual isolates and sequenced as described previoulsy^7^, except for *AvrLm11*, for which AvrLm-11-long-BIS-F (TGTTCGTAGGATAAGTAGGTATTAC) and AvrLm-11-R (CCTCGTTTATTAAGTGTCCTCTGTGCC) pRlmers were used for PCR and sequencing. PCR fragments were sequenced with Sanger technology.

### 2.9 *AvrLm* allele databases

An in-house *L. maculans AvrLm* database (Database V1_2022 with 87 alleles for 63 isoforms, available at https://bioinfo.bioger.inrae.fr/portal/data-browser/public/leptosphaeria/avirulencegenes.) was constructed using verified *AvrLm* sequences^7^. Additional *AvrLm3* and *AvrLm4-7* alleles since described^14^ were added to the VI database, generating the V2_2022 database with 252 alleles for 220 isoforms. An implemented version (Database V3_2023) containing 305 alleles for 247 isoforms was constructed adding new sequences from a set of international *L. maculans* isolates^2^.

RunA was analysed with the V2_2022 database and RunB with the V3_2023 database enriched with the 206 new alleles MiSeq detected in RunA.

New alleles resulting from this study (RunB) were implemented in the V4 version of the database, available at https://bioinfo.bioger.inrae.fr/portal/data-browser/public/leptosphaeria/avirulencegenes.

### 2.10 Analysis of sequence dataset

For each sequencing sample, forward (R1) and reverse (R2) reads were paired using the tool Pear - Paired-End read merger (Galaxy Version 0.9.6.2) of Galaxy pipeline (https://vm-galaxy-prod.toulouse.inrae.fr/galaxy_main). Then, the data were processed gene by gene^18^. A stringent pipeline was used to filter and process paired sequences, utilising MOTHUR v.1.36.1 and parameters adapted to each gene^18^. Chimeras were removed, and the remaining sequences were clustered of into ASVs (Amplicon Sequence Variants), based on 100% identity across the full sequence length^18^. ASVs were assigned to the corresponding *AvrLm* alleles using the databases V2 (RunA) or V3 (RunB), to distinguish known alleles with 100% identity from sequences with less than 100% identity, considered as new alleles.

### 2.11 Selection of relevant ASVs

Three filters were sequentially applied to retain only reliable ASVs: ASVs had to be detected in at least three samples, have read numbers above a gene-specific threshold, and should not be present in negative controls (Table 2). Mean read counts across the three replicates were then calculated for each ASV and sequencing sample^18^. Finally, the resulting count table was simplified by considering an ASV absent if its proportion in the sample was lower than a gene-specific threshold based on the expected and observed ASV proportions in controls^18^ (Table 2).

**Table 2.** Sequencing statistics for the 2022-23 MPSeqM run including control DNA and number of ASVs retained at each filtering step.

| Gene | Mean read no. per sample (min;max)<sup>a</sup> | No. of ASVs in at least 3 samples | Max. Read no. threshold<sup>b</sup> | No. of ASVs above the threshold | No. of ASVs retained after analysis of control DNA<sup>c</sup> | Threshold for the detection of an ASV in a field sampled<sup>d</sup> | No. of ASVs retained above threshold | Final no. of new ASVs | Final no. of new RIPed ASVs<sup>e</sup> | Final no. of new isoforms (full lenght)<sup>f</sup> |
|---|---|---|---|---|---|---|---|---|---|---|
| *AvrLm1* | 936 (0 ; 8922) | 2320 | 14 | 152 | 6 | 1% | 6 | 2 | 1 | 0 |
| *AvrLm2* | 8199 (5825 ; 12405) | 18473 | 25 | 60 | 8 | 0.5% | 7 | 1 | 0 | 0 |
| *AvrLm3* | 4880 (3738 ; 8214) | 17693 | 11 | 382 | 52 | 0.4% | 52 | 20 | 0 | 7 |
| *AvrLm4-7* | 3267 (3 ; 4806) | 14034 | 11 | 292 | 138 | 0.55% | 132 | 59 | 22 | 22 |
| *AvrLm5-9* | 4831 (1583 ; 8905) | 9832 | 10 | 256 | 23 | 0.3% | 23 | 15 | 0 | 5 |
| *AvrLm6* | 3231 (0 ; 5713) | 8352 | 10 | 208 | 9 | 0.4% | 9 | 2 | 0 | 2 |
| *AvrLm11* | 3345 (4 ; 5090) | 6878 | 10 | 165 | 29 | 0.4% | 26 | 7 | 0 | 3 |
| *AvrLmS-Lep2* | 4446 (4 ; 6056) | 18984 | 11 | 188 | 58 | 0.55% | 58 | 18 | 0 | 2 |
| *Total* |  | 96566 |  | 1703 | 323 |  | 313 | 124 | 23 | 41 |

<sup>a</sup> minimum and maximum reads per sample including control and environmental samples
<sup>b</sup> All ASVs for which read counts are below these values in all samples are removed. These values are gene- and run- dependent
<sup>c</sup> based on unexpected ASVs observed in controls and with similar read counts in field samples
<sup>d</sup> This “percentage threshold” represents the percentage of read counts for an AvrLm ASV over all read counts for this gene in a given sample. All values with ASV read counts below this threshold are replaced by zero in the final dataset
<sup>e</sup> A sequence is considered as RIPped if it displays a few mutations from C to T or G to A, introducing stop codons in the sequence
<sup>f</sup> The number of new ASVs does not include new ASVs detected in Gautier et al., 2025

### 2.12 Detection and quantification of *AvrLm* gene deletion

Proportions of deleted alleles of *AvrLm1, AvrLm4-7, AvrLm11*, or *AvrLmS-Lep2* in field samples were estimated using dilution curves^18^. However, for some avirulence genes, deletion curves were not available and the deletion was calculated as follows. For each sample and *AvrLm* gene, the ratio of total *AvrLm* reads to *actin* reads was calculated. The MIX_T control was used as a reference. The *AvrLm/actin* ratio of MIX_T was set to 1. For each sample, the relative proportion of *AvrLm* was calculated with respect to this reference value. Finally, the estimated deletion level for each sample was obtained by subtracting this proportion from 1.

### 2.13 Statistical analysis

The non-parametric Mann-Whitney or Spearman tests were used to assess differences or correlations between expeRlmental groups, respectively. Hierarchical heatmaps were generated with the R package pheatmap (version 10.0.12) to visualize patterns and clustering among the samples based on their similarity. MDS (MultiDimentional Scaling) ordination based on Bray-Curtis similarity matrix was performed with the phyloseq R package (version 1.52.0). All analyses were conducted following standard statistical procedures and with default package parameters^21^. Analyses and graphical representations were conducted within the R statistical environment. Exact confidence intervals of *AvrLm* proportions were calculated at https://www2.dijon.inrae.fr/didactepic/exactci/exactci.php.

## 3 RESULTS

### 3.1 Virulence profiling of *L. maculans* populations on a plant differential set

In 2019-20, the frequency of virulent isolates toward *Rlm1* and *Rlm4* was higher than 95%, and reached 100% for virulence on *Rlm2* and Rlm9^15,18^ (Table 1). Therefore, *AvrLm1, AvrLm2* and *AvrLm9* were not further analysed in the 2022-23 survey, and Pixel *(Rlm4)* was used as a susceptible check in inoculation tests. Only virulence on the three effective resistance sources *Rm6, Rlm11* and *RlmS-LepR2*, (with respectively 100%, 96.1% and 99.9% of avirulent isolates in 2019-20^18^ (Fig. 1) were assessed in 2022-23.

As regards *RlmS-LepR2, we* observed a limited but not significant progression of virulent isolates in the 2022-23 survey. While only one virulent isolate was detected in 2019 (Lange, Table 1), virulent isolates were detected in 2022-23 in four out of 11 sites, with frequencies ranging from 1.3 to 2.2 (Table 1). At the national scale, virulent isolates toward *RlmS-LepR2* represented 0.5% of 716 isolates analysed (Fig. 1, Table 1).

For *Rlm6*, 100% of the isolates were still avirulent in 2022-23, except at Bretenière, displaying 40% of avirulent isolates only (Table 1, Fig. 1). Similarly, proportions of virulent isolates on *Rlm11* were low for the two surveys for all sites (mean 3.4% in 2019-20, 3.8% in 2022-23), except again at Bretenière displaying 21.5% of virulent isolates (Table 1, Fig. 1). In addition, a heterogeneous phenotype was observed for *AvrLm11* at Bretenière. This heterogeneous phenotype (termed “H”) was characterized by an important variability of symptoms, either among plants within the same expeRlment or between expeRlments. The symptoms varied from an intermediate avirulence (grey symptoms of medium size, with a black margin) to a slight virulence (grey symptoms of medium size without margin or pycnidia) (Supporting information Fig. S4). Isolates displaying this heterogeneous *AvrLm11* phenotype represented 26% of the Bretenière population.

The specificity of the Bretenière site was also evident for virulence on *Rlm4*, with only 53.2% of virulent isolates compared to 95% to 100% for other sites (Table 1). Because of these particularities and the lack of historical data for this site, it was further analysed for virulence on *Rlm3* and *Rlm7*. Bretenière also diverged from other sites for these two interactions, with low proportions of avirulent isolates on *Rlm3* (2.6% compared to 51.7% at the national scale in 2019-20) and high proportions of avirulent isolates on *Rlm7* (81% compared to 29.3% at the national scale in 2019-20; Fig. 1, Table 1).

Altogether, the 2022-23 sampling indicated a rather stable proportion of virulent isolates, compared to 2019-20, for all genes investigated, but identified Bretenière as a very divergent site compared to others.

### 3.2 Molecular characterization of *AvrLm* alleles with the MPSeqM tool: technical considerations

The MPSeqM tool had been previously designed and validated using the 2019-20 survey^18^ and used here to characterize the 2022-23 populations. Whatever the avirulence gene analysed, the reproducibility of the method was evident as shown by hierarchical clustering of ASV proportions of control plant samples in RunA and RunB (Fig. 2).

**Figure 2.**
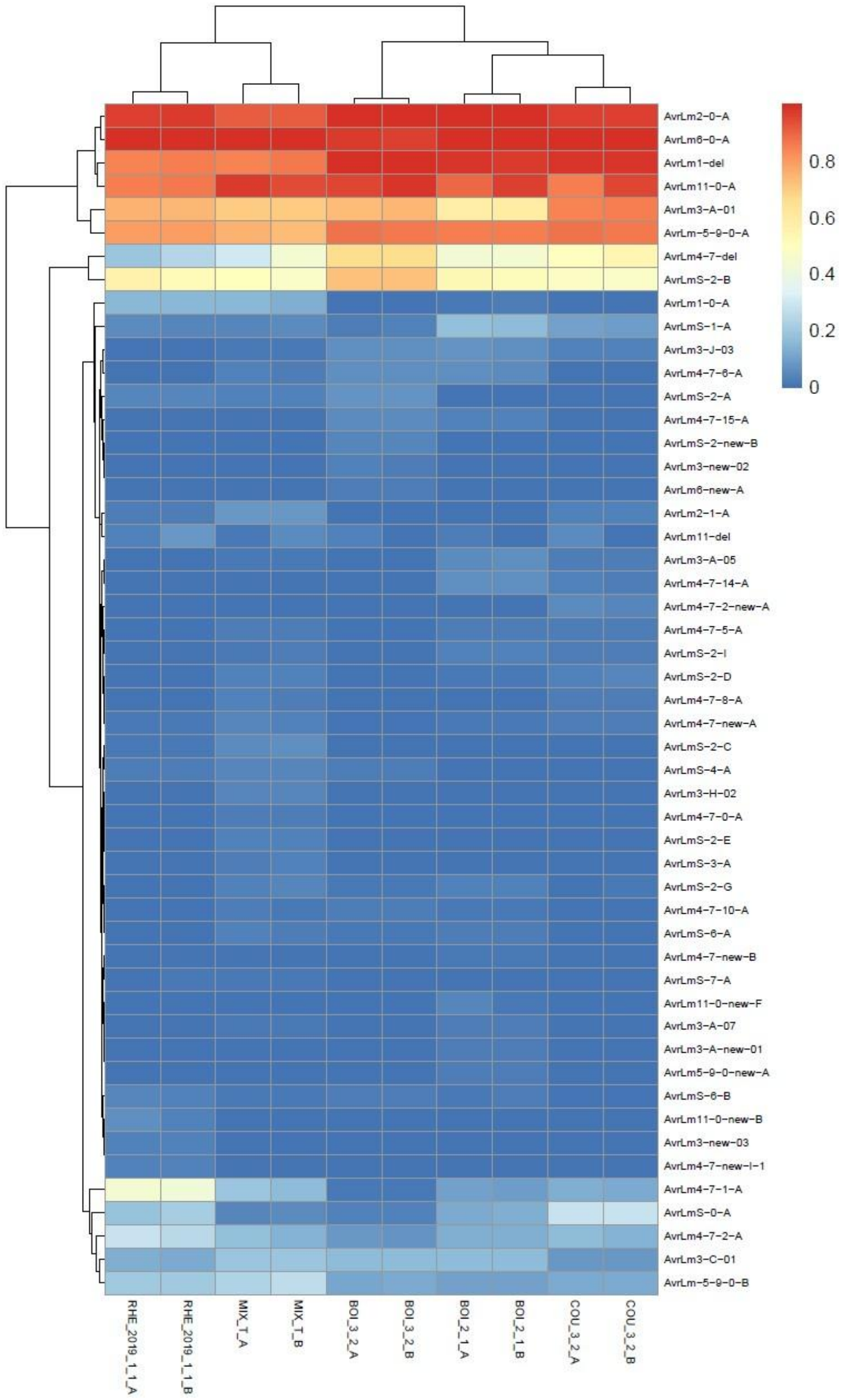
Hierarchical clustering of ASV proportions determined in RunA or in RunB for plant control samples. Four environmental samples from RunA (2019-20 sampling; “RHE”, Le Rheu; “BOI”, Boisseau; “COU”, Coullons) displaying distinct ASV proportions were selected and used as controls in RunB along with the plant control inoculated with known isolates (“MIX_T”). The heatmap shows proportions of 50 most detected ASVs after hierarchical clustering based on similarity of sample profiles. Rows correspond to avirulence alleles as indicated on the left, and columns to the control samples from RunA and RunB. Colors indicate the ASV proportion from low (blue) to high (red).

In environmental samples, we detected 313 reliable ASVs for eight avirulence genes (Table 2), including 124 unknown ASVs, corresponding to 41 new isoforms of the avirulence proteins (Table 2, Fig. 3). New ASVs were detected for all genes. The number of ASVs per gene was highly correlated between runs (R_Spearman_ = 0.934, p<0.001). In accordance with published data^2,7,18^, *AvrLm4-7, AvrLmS-Lep2*, and *AvrLm3* were the most polymorphic genes (Fig. 4, Supporting information Fig. S5), supporting the efficacy of MPSeqM to describe *AvrLm* gene diversity. Interestingly, 52 new ASVs (27 new isoforms) were detected in both sampling years. This suggests that they correspond to alleles steadily present in the *L. maculans* populations.

**Figure 3.**
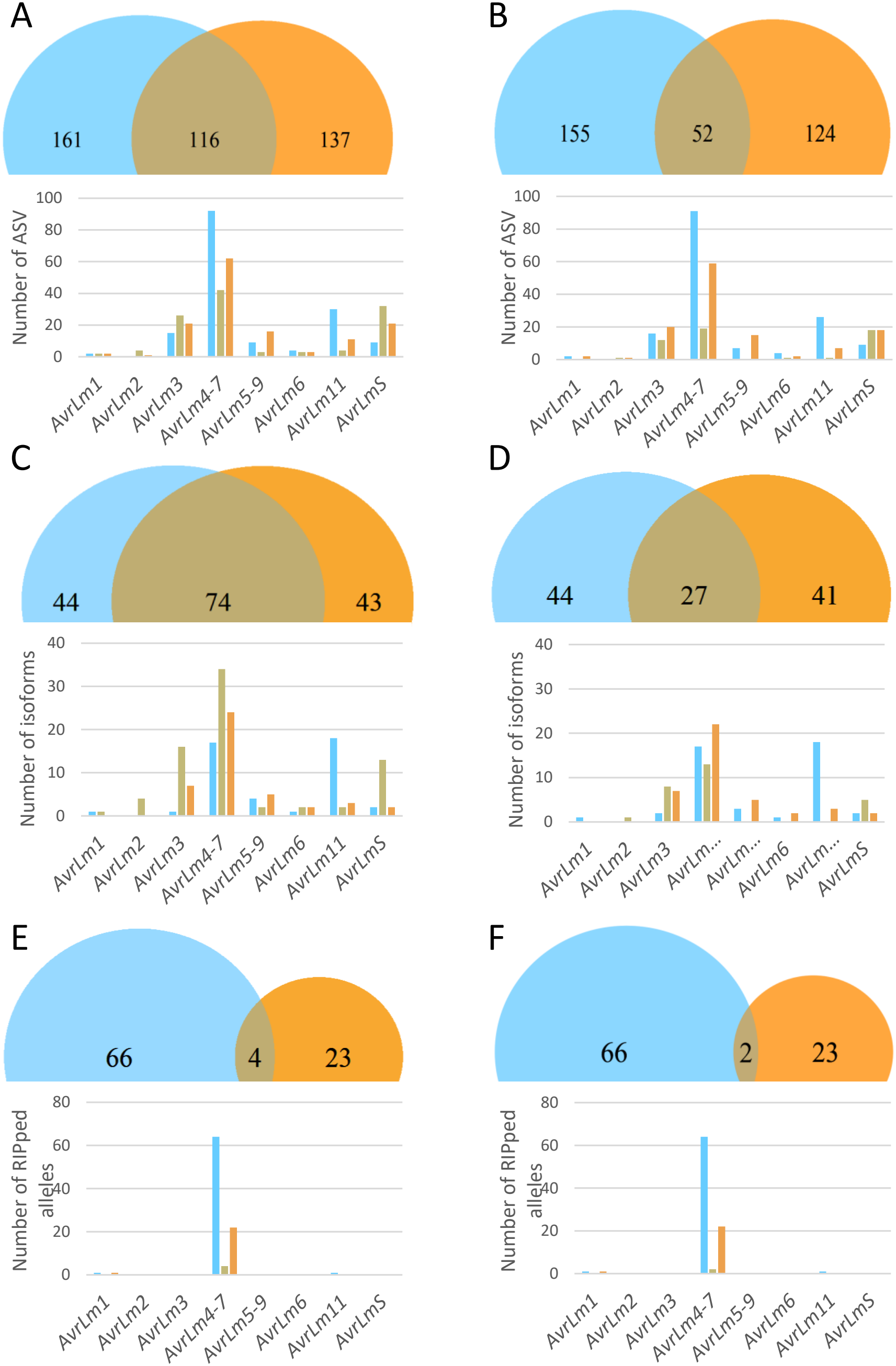
Number of *Leptosphaeria maculans AvrLm* alleles, isoforms, and RIPped alleles detected by MPSeqM. The Venn diagrams indicate the total number of detected ASVs above the thresholds (A, B); the number of corresponding isoforms (C, D); and the number of RIPped alleles (E, F). The histograms represent their repartition between the eight amplified *AvrLm* genes. In blue, ASVs or isoforms specifically detected in RunA (2019-20 samples); in orange, for RunB (2022-23 samples); and in brown, common ASVs or isoforms between the two runs. A, C and D, total allele, isoform and RIPped allele values, respectively; B, D, F, new alleles, isoforms and RIPped alleles, respectively, compared to previous databases.

**Figure 4.**
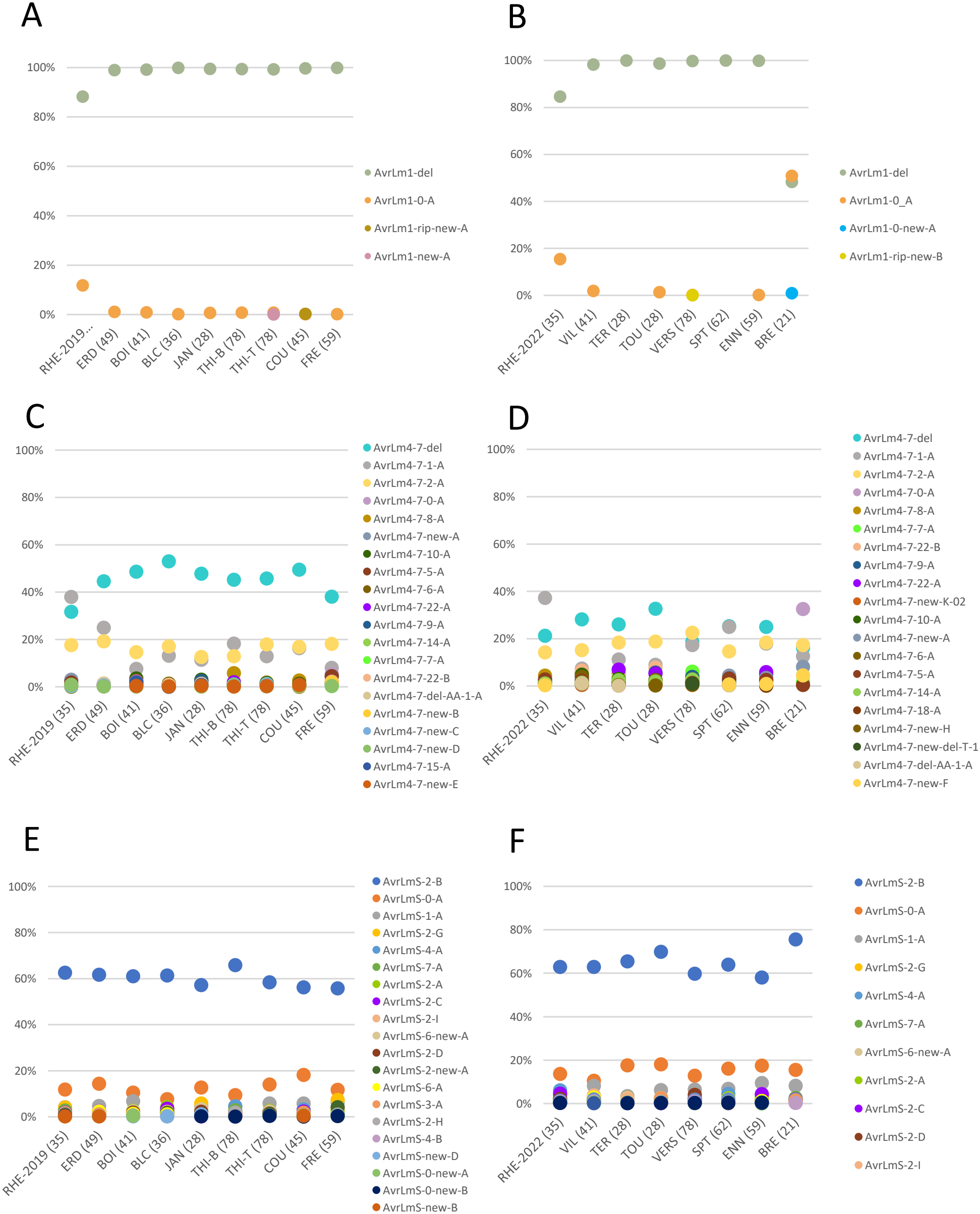
Regional distribution of *Leptosphaeria maculans* alleles of three avirulence genes for two growing seasons. A and B, *AvrLm1*; C and D, *AvrLm4-7*; E and F, *AvrLmS-Lep2*. A, C and E, 2019-20 survey; B, D, F, 2022-23 survey. Each point represents the frequency (%) of a specific allele within the regional population. The **color code identifies each allelic variant**.

Among new alleles, 23 corresponded to RIPped alleles, i.e., with numerous mutations from C-to-T or G-to-A introducing stop codons and inactivated versions of the corresponding proteins^22^. Here, such mutations were mainly detected for *AvrLm4-7*. Most of them (87%) were not detected in the previous sampling (Table 2, Fig. 3, Supporting information file S6). This suggests that *AvrLm4-7* is particularly subject to RIP mutations compared to other *AvrLm* genes and that this mechanism is still active at high frequencies.

The following MPSeqM step is the conversion of ASV proportions into virulent isolate proportions. For that, it is necessary to deduce the impact of each new mutation on the corresponding isoform and then on the phenotype of the isolate (Supporting information Table S7). Some mutations are easily characterized as leading to avirulent or virulent phenotypes^18^. However, others cannot be converted into a phenotype. Their proportion varies between genes, sites, and years (Supporting information Table S8). The highest “nonassigned ASV” proportion was observed for *AvrLm4-7* (2.22% and 3.39% in mean, reaching at some specific sites 4.5% and 6.9%, in 2019-20 and 2022-23, respectively). However, this proportion did not exceed 1% on average over all genes, sites, and years (Supporting information Table S8).

### 3.3 MPSeqM detects geographical structuring of the population

Following MDS analysis, the three samples from Bretenière clearly differed from the others (Fig. 5A). When the samplings from Bretenière were excluded, a clear divergence for nine of the 12 samplings from Le Rheu appeared (Fig. 5B). Besides that, no additional population structuration was detected. Hierarchical clustering of all data also revealed the specificity of the Bretenière population (Supporting information Fig. S9). Additionally, it illustrates a genotypic difference between the three Bretenière samples, with samples Bre_l and Bre_2 clustered apart from Bre_3.

**Figure 5.**
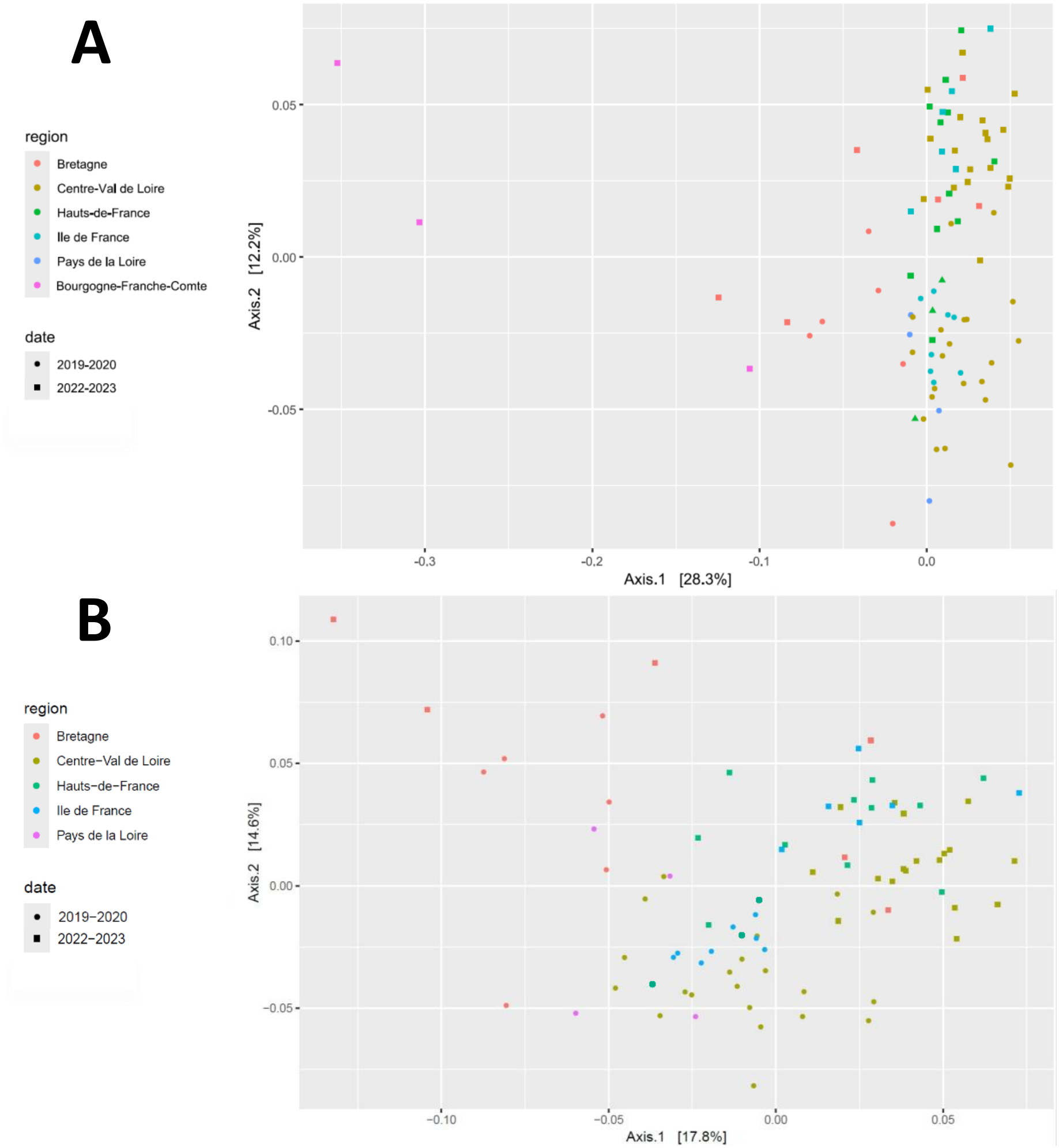
Multidimensional scaling (MDS) analysis of the samples showing the distribution of allelic diversity based on Bray-Curtis dissimilarities. Each dot corresponds to one DNA sample (32 pooled leaf spots). Shapes represent the sampling year, and colours represent the geographic region. (A) All regions are included. (B) Bretenière samples were excluded.

### 3.4 MPSeqM estimation of virulent isolate proportions per site and *AvrLm* gene in 2022-23

The MPSeqM data from RunB (2022-23 sampling) were converted into proportions of virulent isolates on 10 *Rlm* genes (eight *AvrLm*, see below; Table 1) and were compared to 2019-20 data (RunA) and to phenotypic data, when available. The 2022-23 sampling was not phenotyped for virulence toward *Rlm1, Rlm2, Rlm3, Rlm5, Rlm7*, and *Rlm9*, while MPSeqM generated data for these interactions.

MPSeqM estimated virulence toward *Rlm1* in 2022-23 as higher than 98% in all sites but two, Le Rheu and Bretenière, displaying 88.2 and 48.3 % of virulent isolates, respectively (Table 1). These proportions were close to what was observed previously in France, including the presence of avirulent isolates in Le Rheu. Virulence is mainly due to deletion, and *AvrLm1_0_A* is the main avirulent allele (Fig. 4). In accordance with previous samplings^13,15^, more than 98.5% of the isolates were found virulent toward *Rlm2* in 2022-23 (Table 1). The Bretinière site displayed a higher proportion (13.7%) of the allele *AvrLm2_l_A*, compared to less than 4% in other sites (Supporting information Fig. S5).

Ten *AvrLm3* ASVs were detected. Two of them, corresponding to the avirulent isoforms AvrLm3-A and AvrLm3-C, were the most common, representing 72% and 16% on average, respectively (Supporting information Fig. S5). Noticeably, ASVs encoding the AvrLm3-C isoform were nearly absent from Bretenière compared to 11% to 21% at the other sites. Most ASVs encoded avirulent isoforms of AvrLm3. However, the final phenotype toward *Rlm3* relies on the *AvrLm4-7* allele, due to the masking effect of AvrLm4-7 on the AvrLm3-Rlm3 interaction^14,16,17^. Taking into account this masking effect, the estimated proportion of virulent isolates on *Rlm3* in 2022-23 ranged between 64.5% and 80% with a mean proportion at 72%, significantly higher than what was measured in 2019-20 (52% in mean) (Table 1).

*AvrLm4-7* induces resistance on *Rlm4* and/or *Rlm7*, depending on the allele^14^. MPSeqM estimates of virulence on *Rlm4* ranged between 92% and 98%, except at Bretenière (61.5%), values close to what was observed by phenotyping (Table 1). As found previously, virulence toward *Rlm7* varied between sites, ranging from 22.3% (Bretenière) to 65.7% (Table 1). Although the mean proportion of virulence on *Rlm7* was lower in 2022-23 (48.9%) than in 2019-20 (58.9%), there was no statistical difference between the two surveys.

*AvrLm5-9* is also recognized by two genes, *Rlm5* and *Rlm9*^1,23^. MPSeqM data detected two main ASVs, *AvrLm5-9_0_A* and *AvrLm5-9_0_B*, encoding the AvrLm5-9_0 isoform and representing altogether more than 98% of *AvrLm5-9* ASVs (Supporting information Fig. S5). However, the proportion of *AvrLm5-9* reads compared to actin reads was clearly lower for Bretenière compared to other sites (data not shown), suggesting this gene could be deleted in 35.75% of the isolates at Bretenière. Altogether, MPSeqM suggests that nearly 100% of the isolates are virulent on *Rlm9* and avirulent on *Rlm5*, the phenotype induced by the AvrLm5-9_0 isoform, except at Bretenière where all isolates are virulent on *Rlm9* and 35.75% are virulent on *Rlm5* due to *AvrLm5-9* deletion (Table 1, Supporting information Fig. S5).

As found in 2019-20, *AvrLm6* showed very low polymorphism, with more than 99% of the reads corresponding to the avirulent allele *AvrLm6_0_A*. However, deletion of the gene was suggested to occur in 38.7% of the Bretenière population, with variation between replicate pools (from 22 to 48%). In addition, a new allele *(AvrLm6_new_B*, 2.4% of the Bretenière population) was detected with a point mutation introducing a stop codon in the sequence (Supporting Information Fig. S5). Altogether, the MPSeqM tool detected 100% of avirulent isolates, except for Bretenière, where 41.1% of isolates were virulent (Table 1).

MPSeqM data for *AvrLmS-Lep2* were comparable for the two growing seasons, with a high level of polymorphism and similar distributions of the main alleles (Fig. 4). Despite this high polymorphism, most alleles encoded avirulent isoforms, with 0 to 0.8% of virulent isolates per site in 2022-23, as observed by pathotyping (Table 1).

As in 2019-20, the avirulent *AvrLm11_0_A* allele represented more than 90% of the ASVs at all sites (Supporting information Fig. S5). A low proportion of virulent isolates was detected at all sites, including Bretenière (Table 1). Thus, an important discrepancy between phenotyping and MPSeqM analysis was detected for *AvrLm11* at this site. This led us to investigate in greater detail the alleles present in the Bretenière population used for pathotyping.

### 3.5 Characterization of virulent alleles in individual isolates from the 2022-23 sampling

The phenotyping of isolates collected in 2022-23 on trap cultivars or cultivars with effective *Rlm* genes enabled us to detect virulent isolates toward *RlmS-LepR2, Rlm6*, and *Rlm11*. A selection of 183 isolates was used for *AvrLm* gene sequencing to enrich our database with new and characterized alleles and to understand discrepancies between pathotyping and MPSeqM data *for AvrLm11*.

*AvrLm6* was sequenced in 120 isolates (105 isolates from Bretinière, including 77 virulent isolates, and 15 avirulent isolates from seven other sites; Supporting information Table S10). All avirulent isolates displayed avirulent alleles. Among the virulent isolates, 44 were deleted for *AvrLm6*, 29 had RIP-degenerated *AvrLm6* sequences, two displayed new virulent alleles, and two displayed avirulent alleles. For these last two isolates, the genotype alone cannot explain the phenotype, potentially due either to a low expression level as already described for *AvrLrrrf-T*^24^ or to a more complex interaction as observed between *AvrLm3* and *AvrLm4-7*^16^.

*AvrLmS_Lep2* was sequenced in five virulent and three avirulent isolates. One virulent isolate was deleted for *AvrLmS-Lep2*, two displayed the virulent allele *AvrLmS-Lep2_9*, and one displayed a new virulent allele. However, one virulent isolate displayed an avirulent allele (Supporting information Table S10), suggesting a more complex virulence determinism again. Among the avirulent isolates sequenced, one displayed a new allele also detected by MPSeqM, allowing us to infer an avirulent phenotype for this allele.

*AvrLm11* phenotyping has revealed an unstable or irreproducible phenotype for some isolates from Bretenière (see above). Thus, isolates were classified as avirulent (A), virulent (V), or H (variable/heterogeneous). *AvrLm11* was amplified and sequenced in 183 isolates from 10 sampling sites (Supporting information Table S10). All 71 isolates from the « A » category displayed the avirulent allele *AvrLm11_0_A*, except for two isolates displaying new alleles. Noticeably, the length of the amplified region varied, ranging from 830 to 910 bp due to a variable number of a 17 bp minisatellite motif in the 5’ region of the gene (Supporting Information Table S10), without correlation with the final phenotype. Among the 88 « V » isolates, 72 were deleted, three possessed a RIPped allele, one a new virulent allele, but 12 possessed the avirulent allele *AvrLm11_0_A*, pointing to a discrepancy between genotype and phenotype. All of them were from Bretenière. The “H” category comprised 24 isolates from Bretenière, and they all possessed the avirulent allele *AvrLm11_0_A*. However, none of them were isolated on the *Rlm11* plant genotype (Darmor-*Rlm11*). Focusing on the 44 virulent isolates recovered from *Rlm11*, including 21 isolates from Bretenière, all but one were deleted for *AvrLm11*.

*AvrLm5-9* was analysed in the 73 isolates from Bretenière sampled on the trap cultivars (Supporting information Table S10). *AvrLm5-9* was deleted in 31 isolates (42.5%, exact confidence interval [0.31-0.55], consistent with the deletion hypothesis estimated at 35.75% by MPSeqM). All isolates deleted for *AvrLm5-9* also possessed a virulent *AvrLm6* allele (Supporting information Table S10). Apart from deletion, six isolates displayed virulent *AvrLm5-9* alleles with RIP or point mutations introducing stop codons, leading to a total of 51 % of virulent isolates on *Rlm5* at Bretenière.

## DISCUSSION

This study describes a new large-scale survey of *L. maculans* populations in France. 716 isolates, sampled in 2022 at 11 sites across the main regions of oilseed rape production, were inoculated onto a plant differential set to assess their virulence against three effective resistance genes, *Rlm6, RlmS-Lep2*, and *Rlm11*. In parallel, the populations of the same sites were analysed for virulence toward 10 *Rlm* genes using the molecular tool MPSeqM.

This study revealed three major findings. Firstly, the proportions of virulent isolates on effective *Rlm* genes remained stable and very low compared to the previous survey done three years ago^7,15,18^. This was expected for *Rlm11* and *Rlm6*, as these two genes are not currently deployed in oilseed rape varieties in France and thus, *L. maculans* populations are not under pressure from these two genes. In contrast, *RlmS-Lep2* has been used since 2015 with cv. Napoli, followed by two varieties registered in 2018 (Adelmo and Feliciano). After that, *RlmS-Lep2* was detected in one to three new varieties each year (https://www.geves.fr/catalogue/). One of these varieties, at least, was commercially successful, as it became a control cultivar for yield in expeRlmental assays for rapeseed registration in France in 2022 (colza_synthèse_varietale_nationale2023_vl.O.pdf). Thus, *L. maculans* populations were under pressure from *RlmS-Lep2* between 2019 and 2022, but the proportion of virulent isolates did not vary significantly between the two surveys, remaining below 1% nationally in 2022 despite a high level of allelic diversity. This suggests that *RlmS-Lep2* remains effective, with potentially good durability. This raises the question of how frequently large-scale pathotyping surveys should be conducted after the introduction of a new resistance gene to provide informative and useful data.

Secondly, one site (Bretenière) differed from the others with high proportions of virulent isolates on *Rlm6* and *Rm11*. In addition, it showed an unusually high proportion of avirulent isolates on *Rlm4* compared with other sites and surveys^13,15^. The MPSeqM analysis confirmed the divergence of this site for the proportion of virulent isolates or the allelic variants identified. This result was unexpected, as previous analyses had concluded that European populations of *L. maculans* were relatively homogeneous^2,9,15^.

This divergent site had never been characterized before. It was located in a region of intense mustard *(Brassica Juncea)* cropping. In addition, the field assay was sown in an expeRlmental station dedicated both to the study of alternative agroecological practices (reduced tillage and no chemicals) and to the selection of *B. Juncea* varieties for mustard production^25,26^. The Bretenière population showed a high proportion of isolates virulent on *Rlm6*, but also on *Rlm5*, two resistance genes found in *B. Juncea*^27,28^. More importantly, virulence toward these two genes was associated, suggesting these isolates are adapted to *B. Juncea*, probably after local selection of virulent isolates favoured by these specific agronomic and expeRlmental contexts. For the first time, deletion of *AvrLm5* in French isolates was detected by MPSeqM and confirmed in individual isolates. These results question the geographical extent of such adapted populations in the proximity of *B. Juncea* fields. This will be the subject of further studies coupling biological and molecular analyses around this site.

Thirdly, this study revealed a major discrepancy between phenotyping of the *AvrLm11-Rlm11* interaction and molecular analyses at Bretenière. This site showed a high proportion of virulent isolates on *Rlm11* (21.5%) compared with other sites (mean 3.7%), as revealed by pathotyping. *Rlm11* is a resistance gene originating from *B. rapa*, and to our knowledge, no *B. rapa* selection pressure is exerted at the Bretenière site. However, numerous contradictory results were noticed for this interaction at this site: (i) for the Bretenière isolates, the pathotyping of this interaction under controlled conditions is often versatile with a lack of reproducibility between expeRlments; (ii) this versatile phenotype is only found for isolates collected on the trap cultivar, not for those collected on *Rlm11*, suggesting versatile isolates are not able to infect *Rlm11* under natural conditions; and (iii) all *AvrLm11* versatile isolates possessed an avirulent allele for *AvrLm11*. These data suggest that an unknown mechanism, specifically observed for isolates from Bretenière, is responsible for a versatile expression of the phenotype evaluated under controlled conditions, leading to virulence despite an avirulent allele, but that under natural infection conditions, such isolates are unable to infect resistant plants. Phenotyping variability under controlled conditions was described for other interactions. For instance, different teams using comparable inoculation techniques with the same isolates and plant material concluded to virulent or avirulent phenotypes for *AvrLmS-Lep2 I RlmS-LepR2* interaction^5^. It remains now to be determined whether inconsistencies between phenotyping and genotyping are linked to variable expression levels of *AvrLmS-Lep2* depending on environmental conditions, or to a more complex determinism of the interaction involving other avirulence or regulatory genes.

Altogether, our study reveals the complementarity between biological and molecular approaches to survey *L. maculans* populations. MPSeqM gives information for avirulence genes that are not easy to phenotype (such as *AvrLm5*, for which the plant differential is lacking). It allows a complete characterization of populations with limited workload (here, toward 10 *Rlm* genes compared to four by pathotyping), and it rapidly reveals sites with major differences compared to others. Focused on currently effective *Rlm* genes, the biological pathotyping allows precise quantification of *Rlm* breakdown and detects a few virulent isolates, whose individual characterization helps to extend the molecular database of virulent alleles. Following our epidemiological surveys experience, we could now propose the following scheme for furthers *L. maculans* surveys: (i), increase time intervals between large-scale surveys to every 4-5 years using biological pathotyping; (ii), focus such surveys on regions of main oilseed rape cultivation; (iii), combine biological surveys with more focused observation, isolation, pathotyping and molecular analyses of isolates recovered from resistant varieties; (iv), in parallel, apply MPSeqM every year or two years on much more sites to detect sites with specific populations; and (v), pathotype, if needed, populations from divergent sites as soon as they are detected by MPSeqM.

### Funding

This work was funded by the CASDAR project C2018-11 (Atipical), and the “Club Phoma” project. The “Effectors and Pathogenesis of *L. maculans”* (EPLM) group benefits from the support of Saclay Plant Sciences-SPS (ANR-17-EUR-0007).

### Authors’ contributions

All authors conceived the study. MHB, VL and AJ coordinated the projects, including funding. AG performed the molecular analyses, including pRlmer design, technical implementation and data analyses. AJ supervised the 2022 field sampling and performed its phenotyping. MHB and VL wrote the main manuscript text with major input from AG and AJ for analytical methods, datasets and figures. All authors reviewed the manuscript.

## Acknowledgements

We thank all contributors from the CTPS field expeRlments and from the Club Phoma network that provided infected oilseed rape leaves, Julie Noah, Marie-Paule Peler and Laurent Coudard for their contribution to leaf spot recovery and isolate management, and the whole EPLM team for help in inoculation tests. We thank the GeT-Biopuces and GeT-PlaGe platforms in Toulouse, Midi-Pyrénées for the MiSeq sequencing expeRlments. We are grateful to the GenoToul Bioinformatics platform in Toulouse, Midi-Pyrénées for providing help in computing and data storage. We thank IGEPP and the BrACySol CRB for Westar, Darmor-/?/m6 and Darmor-R/m11 genotypes and AAFC Saskatoon for the Topas-Lep/R2 genotype.

## Data availability

Raw sequencing data (Run B) are available at https://www.ncbi.nlm.nih.gov/sra/PRJNA1446249

## Notes

### Competing Interest Statement

The authors have declared no competing interest.

## REFERENCES

1. Borhan MH, Van de Wouw AP and Larkan NJ, Molecular interactions between Leptosphaeria maculans and Brassica species. Annu Rev Phytopathol 60:237–257 (2022). 10.1146/annurevphyto-021621-120602.

2. Van de Wouw AP, Scanlan JL, Al-Mamun HA, Balesdent MH, Bousset L, Burketová L et al., A new set of international Leptosphaeria maculans isolates as a resource for elucidation of the basis and evolution of blackleg disease on Brassica napus. Plant Pathol 73:170–185 (2024).

3. West JS, Fitt BDL, Leech PK, Biddulph JE, Huang YJ and Balesdent MH, Timing of Leptosphaeria maculans ascospore release, phoma leaf spot development and fungicide use in relation to stem canker development on winter oilseed rape (Brassica napus) in southern England. Plant Pathol 51:454–463 (2002).

4. Delourme R, Chèvre AM, Brun H, Rouxel T, Balesdent MH, Dias JS et al., Major gene and polygenic resistance to Leptosphaeria maculans in oilseed rape (Brassica napus). Eur J Plant Pathol 114:41–52 (2006).

5. Neik TX, Ghanbarnia K, Ollivier B, Scheben A, Severn-Ellis A, Larkan NJ, et al,. Two independent approaches converge to the cloning of a new Leptosphaeria maculans avirulence effector gene, AvrLmS-Lep2. Mol Plant Pathol 23:733–748 (2022).

6. Rouxel T and Balesdent MH. Life, death and rebirth of avirulence effectors in a fungal pathogen of Brassica crops, Leptosphaeria maculans. New Phytol 214:526–532 (2017).

7. Gautier A, Laval V, Faure S, Rouxel T and Balesdent MH, Polymorphism of avirulence genes and adaptation to Brassica resistance genes is gene-dependent in the phytopathogenic fungus Leptosphaeria maculans. Phytopathology 113:1222–1232 (2023).

8. McDonald BA and Linde C, Pathogen population genetics, evolutionary potential, and durable resistance. Annu Rev Phytopathol 40:349–379 (2002).

9. Alnajar D, von Tiedemann A and Koopmann B, Efficacy of blackleg major resistance genes in B. napus in Germany. Pathogens 11:461 (2022).

10. Liu F, Zou Z, Peng G and Fernando WGD, Leptosphaeria maculans isolates reveal their allele frequency in Western Canada. Plant Dis 105:1440–1447 (2021).

11. Stachowiak A, Olechnowicz J, Jedryczka M, Rouxel T, Balesdent MH, Happstadius I et al., Frequency of avirulence alleles in field populations of Leptosphaeria maculans in Europe. Eur J Plant Pathol 114:67–75 (2006).

12. Van de Wouw AP, Howlett BJ and Idnurm A, (2017). Changes in allele frequencies of avirulence genes in the blackleg fungus, Leptosphaeria maculans, over two decades in Australia. Crop Pasture Sci 69:20–29 (2017). 10.1071/CP16411

13. Balesdent MH, Louvard K, Pinochet X and Rouxel T, A large-scale survey of races of Leptosphaeria maculans occurring on oilseed rape in France. Eur J Plant Pathol 114:53–65 (2006).

14. Balesdent MH, Gautier A, Plissonneau C, Le Meur L, Loiseau A, Leflon M, et al., Twenty years of Leptosphaeria maculans population survey in France suggests pyramiding Rlm3 and Rlm7 in rapeseed is a risky resistance management strategy. Phytopathology 112:2126–2137 (2022).

15. Balesdent MH, Laval V, Bagot P and Rouxel T, Large-scale population survey of Leptosphaeria maculans in France highlights both on-going breakdowns and potentially effective resistance genes in oilseed rape. Pest Manag Sci 80:2426–2434 (2024). 10.1002/ps.7401

16. Plissonneau C, Daverdin G, Ollivier B, Blaise F, Degrave A, Fudal I et al., A game of hide and seek between avirulence genes AvrLm4-7 and AvrLm3 in Leptosphaeria maculans. New Phytol 209:1613–1624 (2016).

17. Plissonneau C, Blaise F, Ollivier B, Leflon M, Carpezat J, Rouxel T et al., 2017, Unusual evolutionary mechanisms to escape Effector-Triggered-Immunity in the fungal phytopathogen Leptosphaeria maculans. Mol Ecol 26:2183–2198 (2017).

18. Gautier A, Laval V and Balesdent MH, MPSeqM, a tool combining multiplex PCR and high-throughput sequencing to study the polymorphism of eight Leptosphaeria maculans avirulence genes and its application to field surveys in France. BMC Microbiol 25:159 (2025). 10.1186/s12866-025-03855-2

19. Balesdent MH, Fudal I, Ollivier B, Bally P, Grandaubert J, Eber F, et al., The dispensable chromosome of Leptosphaeria maculans shelters an effector gene conferring avirulence towards Brassica rapa. New Phytol 198:887–898 (2013).

20. Dilmaghani A, Balesdent MH, Didier JP, Wu C, Davey J, Barbetti MJ, et al., The Leptosphaeria maculans-Leptosphaeria biglobosa species complex in the American continent. Plant Pathol 58:1044–1058 (2009).

21. McMurdie PJ and Holmes S, phyloseq: An R package for reproducible interactive analysis and graphics of microbiome census data. PLoS ONE 8:e61217 (2013).

22. Fudal I, Ross S, Brun H, Besnard AL, Ermel M, Kuhn ML et al., Repeat-induced point mutation (RIP) as an alternative mechanism of evolution towards virulence in Leptosphaeria maculans. Mol Plant-Microbe Interact 22 :932–941 (2009).

23. Ghanbarnia K, Ma L, Larkan NJ, Haddadi P, Fernando WGD and Borhan MH, Leptosphaeria maculans AvrLm9: a new player in the game of hide and seek with AvrLm4-7. Mol Plant Pathol 19:1754–1764 (2018). doi: 10.1111/mpp.12658.

24. Daverdin G, Rouxel T, Gout L, Aubertot JN, Fudal I, Meyer M et al., Genome structure and reproductive behaviour influence the evolutionary potential of a fungal phytopathogen. PLoS Pathogens 8: e1003020 (2012).

25. Petit S, Deytieux V and Cordeau S, Landscape-scale approaches for enhancing biological pest control in agricultural systems. Environ Monit Assess 193:75 (2021). 10.1007/s10661-020-08812-2

26. Le Gall, S., Sole-Jamault, V., Nars-Chasseray, M., Le Goff, A., Le Bot, L., Guinet, T., Renaud, C., Gervais, J., Bansard, S., Ohleyer, L. and Jeandroz, S., Data on agronomic traits, biochemical composition of lipids, proteins and polysaccharides and rheological measurement in a brown mustard seed collection. Data in Brief, 38:107417 (2021).

27. Balesdent MH, Attard A, Kühn ML and Rouxel T, New avirulence genes in the phytopathogenic fungus Leptosphaeria maculans. Phytopathology 92:1122–1133 (2002).

28. Inturrisi FC, Barbetti MJ, Tirnaz S, Patel DA, Edwards D, and Batley J, Molecular characterization of disease resistance in Brassica juncea – The current status and the way forward. Plant Pathol. 70:13–34 (2021). 10.1111/ppa.13277

