## Supplementary data for "Unexpected population structure and plant-pathogen interaction as revealed by very large-scale molecular and biological analyses of *Leptosphaeria maculans* avirulence genes in France"

**Supplementary Table S1. Genotypes of reference *Leptosphaeria maculans* isolates at eight avirulence loci. Isolates in bold are those used in the mix of 32 *Brassica napus* leaf spots used as control.**

| Isolate[1] | *AvrLm1* | *AvrLm2* | *AvrLm3* | *AvrLm4-7* | *AvrLm5-9* | *AvrLm6* | *AvrLm11* | *AvrLmS-Lep2* |
|---|---|---|---|---|---|---|---|---|
| **EPH17.001** | *AvrLm1_del*[2] | *AvrLm2_0_A* | *AvrLm3_A_01* | *AvrLm4-7_del* | *AvrLm5-9_0_A* | *AvrLm6_0_A* | *AvrLm11_0_A* | *AvrLmS_2_B* |
| **EPH17.004** | *AvrLm1_del* | *AvrLm2_0_A* | *AvrLm3_A_01* | *AvrLm4-7_del* | *AvrLm5-9_0_A* | *AvrLm6_0_A* | *AvrLm11_0_A* | *AvrLmS_7_A* |
| **EPH17.010** | *AvrLm1_del* | *AvrLm2_0_A* | *AvrLm3_C_01* | *AvrLm4-7_rip_1* | *AvrLm5-9_0_B* | *AvrLm6_0_A* | *AvrLm11_0_A* | *AvrLmS_2_B* |
| **EPH17.016** | *AvrLm1_del* | *AvrLm2_0_A* | *AvrLm3_A_09* | *AvrLm4-7_9_A* | *AvrLm5-9_0_A* | *AvrLm6_0_A* | *AvrLm11_0_A* | *AvrLmS_2_B* |
| **EPH17.019** | *AvrLm1_del* | *AvrLm2_0_A* | *AvrLm3_A_09* | *AvrLm4-7_partial_del* | *AvrLm5-9_0_B* | *AvrLm6_0_A* | *AvrLm11_0_A* | *AvrLmS_2_F* |
| EPH17.029 | *AvrLm1_del* | *AvrLm2_0_A* | *AvrLm3_A_01* | *AvrLm4-7_del* | *AvrLm5-9_0_A* | *AvrLm6_0_A* | *AvrLm11_del* | *AvrLmS_2_B* |
| **EPH17.066** | *AvrLm1_del* | *AvrLm2_0_A* | *AvrLm3_H_02* | *AvrLm4-7_partial_del* | *AvrLm5-9_0_A* | *AvrLm6_0_A* | *AvrLm11_0_A* | *AvrLmS_4_A* |
| **EPH17.071** | *AvrLm1_rip3* | *AvrLm2_0_A* | *AvrLm3_C_01* | *AvrLm4-7_1_A* | *AvrLm5-9_0_A* | *AvrLm6_0_A* | *AvrLm11_0_A* | *AvrLmS_6_A* |
| **EPH17.084** | *AvrLm1_del* | *AvrLm2_0_A* | *AvrLm3_A_01* | *AvrLm4-7_5_A* | *AvrLm5-9_0_B* | *AvrLm6_0_A* | *AvrLm11_0_A* | *AvrLmS_2_B* |
| **EPH17.102** | *AvrLm1_del* | *AvrLm2_1_A* | *AvrLm3_H_02* | *AvrLm4-7_del* | *AvrLm5-9_0_A* | *AvrLm6_0_A* | *AvrLm11_0_A* | *AvrLmS_2_B* |
| **EPH17.105** | *AvrLm1_del* | *AvrLm2_0_A* | *AvrLm3_A_09* | *AvrLm4-7_6_A* | *AvrLm5-9_0_A* | *AvrLm6_0_A* | *AvrLm11_0_A* | *AvrLmS_2_A* |
| **EPH17.108** | *AvrLm1_del* | *AvrLm2_0_A* | *AvrLm3_A_09* | *AvrLm4-7_rip_2* | *AvrLm5-9_0_B* | *AvrLm6_0_A* | *AvrLm11_0_A* | *AvrLmS_2_B* |
| EPH17.111 | *AvrLm1_del* | *AvrLm2_0_A* | *AvrLm3_A_09* | *AvrLm4-7_5_A* | *AvrLm5-9_0_A* | *AvrLm6_0_A* | *AvrLm11_0_A* | *AvrLmS_0_A* |
| **EPH17.114** | *AvrLm1_del* | *AvrLm2_0_A* | *AvrLm3_A_09* | *AvrLm4-7_7_A* | *AvrLm5-9_0_A* | *AvrLm6_0_A* | *AvrLm11_0_A* | *AvrLmS_2_B* |
| **EPH17.117** | *AvrLm1_del* | *AvrLm2_0_A* | *AvrLm3_A_09* | *AvrLm4-7_2_A* | *AvrLm5-9_0_A* | *AvrLm6_0_A* | *AvrLm11_0_A* | *AvrLmS_2_B* |
| **EPH17.120** | *AvrLm1_del* | *AvrLm2_0_A* | *AvrLm3_A_09* | *AvrLm4-7_del_AA_A* | *AvrLm5-9_0_A* | *AvrLm6_0_A* | *AvrLm11_del* | *AvrLmS_2_B* |
| EPH17.124 | *AvrLm1_del* | *AvrLm2_0_A* | *AvrLm3_H_02* | *AvrLm4-7_partial_del* | *AvrLm5-9_0_A* | *AvrLm6_0_A* | *AvrLm11_0_A* | *AvrLmS_2_B* |
| **EPH17.127** | *AvrLm1_del* | *AvrLm2_0_A* | *AvrLm3_A_01* | *AvrLm4-7_1_A* | *AvrLm5-9_0_A* | *AvrLm6_0_A* | *AvrLm11_0_A* | *AvrLmS_2_B* |
| EPH17.130 | *AvrLm1_del* | *AvrLm2_0_A* | *AvrLm3_A_01* | *AvrLm4-7_ripdup_A* | *AvrLm5-9_0_A* | *AvrLm6_0_A* | *AvrLm11_0_A* | *AvrLmS_2_B* |
| **EPH17.133** | *AvrLm1_del* | *AvrLm2_0_A* | *AvrLm3_H_02* | *AvrLm4-7_del* | *AvrLm5-9_0_C* | *AvrLm6_0_A* | *AvrLm11_0_A* | *AvrLmS_2_D* |
| **EPH17.142** | *AvrLm1_del* | *AvrLm2_0_A* | *AvrLm3_A_01* | *AvrLm4-7_8_A* | *AvrLm5-9_0_A* | *AvrLm6_0_A* | *AvrLm11_0_A* | *AvrLmS_1_A* |
| EPH17.145 | *AvrLm1_del* | *AvrLm2_0_A* | *AvrLm3_A_01* | *AvrLm4-7_2_A* | *AvrLm5-9_0_A* | *AvrLm6_0_A* | *AvrLm11_0_A* | *AvrLmS_7_A* |
| EPH17.151 | *AvrLm1_del* | *AvrLm2_0_A* | *AvrLm3_A_09* | *AvrLm4-7_1_A* | *AvrLm5-9_0_A* | *AvrLm6_0_A* | *AvrLm11_0_A* | *AvrLmS_0_A* |
| **EPH17.157** | *AvrLm1_del* | *AvrLm2_1_A* | *AvrLm3_A_09* | *AvrLm4-7_2_A* | *AvrLm5-9_0_C* | *AvrLm6_0_A* | *AvrLm11_0_A* | *AvrLmS_2_B* |
| **EPH17.164** | *AvrLm1_del* | *AvrLm2_0_A* | *AvrLm3_C_01* | *AvrLm4-7_2_A* | *AvrLm5-9_0_B* | *AvrLm6_0_A* | *AvrLm11_0_A* | *AvrLmS_2_B* |
| EPH17.167 | *AvrLm1_del* | *AvrLm2_0_A* | *AvrLm3_A_01* | *AvrLm4-7_1_A* | *AvrLm5-9_0_A* | *AvrLm6_0_A* | *AvrLm11_0_A* | *AvrLmS_4_A* |
| EPH17.179 | *AvrLm1_del* | *AvrLm2_0_A* | *AvrLm3_A_01* | *AvrLm4-7_1_A* | *AvrLm5-9_0_A* | *AvrLm6_0_A* | *AvrLm11_0_A* | *AvrLmS_1_A* |

| | | | | | | | | |
|---|---|---|---|---|---|---|---|---|
| **EPH17.186** | *AvrLm1_del* | *AvrLm2_0_A* | *AvrLm3_A_09* | *AvrLm4-7_1_A* | *AvrLm5-9_0_A* | *AvrLm6_0_A* | *AvrLm11_0_A* | *AvrLmS_2_C* |
| **EPH17.194** | *AvrLm1_del* | *AvrLm2_0_A* | *AvrLm3_A_01* | *AvrLm4-7_10_A* | *AvrLm5-9_0_A* | *AvrLm6_0_A* | *AvrLm11_0_A* | *AvrLmS_2_B* |
| **EPH17.201** | *AvrLm1_del* | *AvrLm2_0_A* | *AvrLm3_A_01* | *AvrLm4-7_del* | *AvrLm5-9_0_B* | *AvrLm6_0_A* | *AvrLm11_0_A* | *AvrLmS_2_E* |
| EPH17.207 | *AvrLm1_del* | *AvrLm2_0_A* | *AvrLm3_A_01* | *AvrLm4-7_2_A* | *AvrLm5-9_0_A* | *AvrLm6_0_A* | *AvrLm11_0_A* | *AvrLmS_0_A* |
| EPH17.216 | *AvrLm1_del* | *AvrLm2_0_A* | *AvrLm3_C_01* | *AvrLm4-7_del* | *AvrLm5-9_0_B* | *AvrLm6_0_A* | *AvrLm11_0_A* | *AvrLmS_2_B* |
| **EPH17.225** | *AvrLm1_del* | *AvrLm2_0_A* | *AvrLm3_I_01* | *AvrLm4-7_partial_del* | *AvrLm5-9_0_A* | *AvrLm6_0_A* | *AvrLm11_0_A* | *AvrLmS_2_C* |
| EPH17.231 | *AvrLm1_del* | *AvrLm2_0_A* | *AvrLm3_A_09* | *AvrLm4-7_del* | *AvrLm5-9_0_A* | *AvrLm6_0_A* | *AvrLm11_0_A* | *AvrLmS_4_A* |
| **EPH17.234** | *AvrLm1_del* | *AvrLm2_0_A* | *AvrLm3_J_03* | *AvrLm4-7_del* | *AvrLm5-9_0_C* | *AvrLm6_0_A* | *AvrLm11_0_A* | *AvrLmS_2_B* |
| **EPH17.238** | *AvrLm1_0_A* | *AvrLm2_1_A* | *AvrLm3_A_09* | *AvrLm4-7_1_A* | *AvrLm5-9_0_A* | *AvrLm6_0_A* | *AvrLm11_0_A* | *AvrLmS_3_A* |
| **EPH17.250** | *AvrLm1_del* | *AvrLm2_0_A* | *AvrLm3_A_07* | *AvrLm4-7_2_A* | *AvrLm5-9_0_A* | *AvrLm6_0_A* | *AvrLm11_del* | *AvrLmS_2_B* |
| **EPH17.256** | *AvrLm1_0_A* | *AvrLm2_0_A* | *AvrLm3_C_01* | *AvrLm4-7_rip_3* | *AvrLm5-9_0_A* | *AvrLm6_0_A* | *AvrLm11_0_A* | *AvrLmS_5_A* |
| EPH17.272 | *AvrLm1_del* | *AvrLm2_0_A* | *AvrLm3_A_01* | *AvrLm4-7_5_A* | *AvrLm5-9_0_C* | *AvrLm6_0_A* | *AvrLm11_0_A* | *AvrLmS_2_D* |
| **EPH17.275** | *AvrLm1_del* | *AvrLm2_1_A* | *AvrLm3_C_01* | *AvrLm4-7_1_A* | *AvrLm5-9_0_A* | *AvrLm6_3_A* | *AvrLm11_0_A* | *AvrLmS_2_B* |
| EPH17.278 | *AvrLm1_0_A* | *AvrLm2_0_A* | *AvrLm3_A_09* | *AvrLm4-7_1_A* | *AvrLm5-9_0_A* | *AvrLm6_0_A* | *AvrLm11_0_A* | *AvrLmS_2_B* |
| **EPH17.287** | *AvrLm1_del* | *AvrLm2_0_A* | *AvrLm3_A_09* | *AvrLm4-7_del* | *AvrLm5-9_0_A* | *AvrLm6_0_A* | *AvrLm11_0_A* | *AvrLmS_2_G* |
| EPH17.296 | *AvrLm1_del* | *AvrLm2_0_A* | *AvrLm3_A_01* | *AvrLm4-7_del* | *AvrLm5-9_0_C* | *AvrLm6_0_A* | *AvrLm11_0_A* | *AvrLmS_1_A* |
| **EPH17.304** | *AvrLm1_del* | *AvrLm2_0_A* | *AvrLm3_J_04* | *AvrLm4-7_del* | *AvrLm5-9_0_A* | *AvrLm6_0_A* | *AvrLm11_0_A* | *AvrLmS_0_A* |
| EPH17.308 | *AvrLm1_del* | *AvrLm2_0_A* | *AvrLm3_A_09* | *AvrLm4-7_1_A* | *AvrLm5-9_0_B* | *AvrLm6_0_A* | *AvrLm11_del* | *AvrLmS_7_A* |
| **EPH17.324** | *AvrLm1_del* | *AvrLm2_0_A* | *AvrLm3_A_05* | *AvrLm4-7_2_A* | *AvrLm5-9_0_A* | *AvrLm6_0_A* | *AvrLm11_0_A* | *AvrLmS_1_A* |
| EPH17.336 | *AvrLm1_0_A* | *AvrLm2_0_A* | *AvrLm3_A_05* | *AvrLm4-7_del* | *AvrLm5-9_0_B* | *AvrLm6_0_A* | *AvrLm11_0_A* | *AvrLmS_0_A* |
| JN2(v23.1.2) | *AvrLm1_del* | *AvrLm2_0_A* | *AvrLm3_A_09* | *AvrLm4-7_1_A* | *AvrLm5-9_0_A* | *AvrLm6_0_A* | *AvrLm11_0_A* | *AvrLmS_1_A* |
| **JN3 (v23.1.3)** | *AvrLm1_0_A* | *AvrLm2_0_A* | *AvrLm3_A_01* | *AvrLm4-7_0_A* | *AvrLm5-9_0_A* | *AvrLm6_0_A* | *AvrLm11_0_A* | *AvrLmS_0_A* |
| BBA-62908 | *AvrLm1_0_A* | *AvrLm2_3_A* | *AvrLm3_G_1* | *AvrLm4-7_4_A* | *AvrLm5-9_1_A* | *AvrLm6_1_A* | *AvrLm11_1_A* | *AvrLmS_8_B* |
| CB-G3.5 | *AvrLm1-rip2* | *AvrLm2_0_A* | *AvrLm3_A_01* | *AvrLm4-7_1_A* | *AvrLm5-9_0_A* | *AvrLm6_0_A* | *AvrLm11_0_A* | *AvrLmS_0_A* |
| CB-i2-9 | *AvrLm1-rip1* | *AvrLm2_0_A* | *AvrLm3_A_09* | *AvrLm4-7_1_A* | *AvrLm5-9_0_A* | *AvrLm6_0_A* | *AvrLm11_0_A* | *AvrLmS_2_B* |
| IBCN14 | *AvrLm1_del* | *AvrLm2_0_A* | *AvrLm3_J_02* | *AvrLm4-7_del* | *AvrLm5-9_0_A* | *AvrLm6_1_A* | *AvrLm11_del* | *AvrLmS_2_B* |
| IBCN18 | *AvrLm1_1_A* | *AvrLm2_3_A* | *AvrLm3_P_01* | *AvrLm4-7_0_A* | *AvrLm5-9_4_A* | *AvrLm6_del* | *AvrLm11-0+2-dup* | *AvrLmS_7_A* |
| PHW1245 (IBCN74) | *AvrLm1_1_B* | *AvrLm2_4_A* | *AvrLm3_F_02* | *AvrLm4-7_3_A* | *AvrLm5-9_1_A* | *AvrLm6_2_A* | *AvrLm11_2_A* | *AvrLmS_8_B* |
| IBCN75 | *AvrLm1_0_A* | *AvrLm2_0_A* | *AvrLm3_J_03* | *AvrLm4-7_del* | *AvrLm5-9_1_A* | *AvrLm6_1_A* | *AvrLm11_0_A* | *AvrLmS_7_A* |

| | | | | | | | | |
|---|---|---|---|---|---|---|---|---|
| IBCN76 | *AvrLm1_0_A* | *AvrLm2_1_A* | *AvrLm3_A_01* | *AvrLm4-7_del* | *AvrLm5-9_0_A* | *AvrLm6_0_A* | *AvrLm11_0_A* | *AvrLmS_2_B* |
| IBCN79 | *AvrLm1_0_A* | *AvrLm2_1_A* | *AvrLm3_A_09* | *AvrLm4-7_del* | *AvrLm5-9_0_A* | *AvrLm6_0_A* | *AvrLm11_0_A* | *AvrLmS_7_A* |
| IBCN80 | *AvrLm1_del* | *AvrLm2_3_A* | *AvrLm3_D_01* | *AvrLm4-7_del* | *AvrLm5-9_1_A* | *AvrLm6_1_A* | *AvrLm11_1_A* | *AvrLmS_8_A* |
| IBCN85 | *AvrLm1_del* | *AvrLm2_2_A* | *AvrLm3_A_09* | *AvrLm4-7_2_A* | *AvrLm5-9_0_A* | *AvrLm6_1_A* | *AvrLm11_0_A* | *AvrLmS_8_B* |
| MX8-12 | *AvrLm1_0_A* | *AvrLm2_1_A* | *AvrLm3_A_01* | *AvrLm4-7_2_A* | *AvrLm5-9_0_B* | *AvrLm6_rip_1* | *AvrLm11_del* | *AvrLmS_4_B* |
| Nz-T4 | *AvrLm1_del* | *AvrLm2_0_A* | *AvrLm3_I_01* | *AvrLm4-7_rip* | *AvrLm5-9_0_A* | *AvrLm6_1_A* | *AvrLm11_0_A* | *AvrLmS_2_A* |
| OMR19 | *AvrLm1_0_A* | *AvrLm2_3_A* | *AvrLm3_F_02* | *AvrLm4-7_3_A* | *AvrLm5-9_1_A* | *AvrLm6_1_A* | *AvrLm11_1_A* | *AvrLmS_8_B* |
| UWAP11 | *AvrLm1_0_A* | *AvrLm2_1_A* | *AvrLm3_A_17* | *AvrLm4-7_1_A* | *AvrLm5-9_5_A* | *AvrLm6_0_A* | *AvrLm11_0_A* | *AvrLmS_2_B* |
| WAC4028 | *AvrLm1_del* | *AvrLm2_1_A* | *AvrLm3_A_09* | *AvrLm4-7_del* | *AvrLm5-9_0_A* | *AvrLm6_3_A* | *AvrLm11_0_A* | *AvrLmS_2_E* |
| WAC4057 | *AvrLm1_0_A* | *AvrLm2_0_A* | *AvrLm3_A_09* | *AvrLm4-7_del* | *AvrLm5-9_0_A* | *AvrLm6_1_A* | *AvrLm11_0_A* | *AvrLmS_2_B* |
| WAC4079 | *AvrLm1_0_A* | *AvrLm2_2_A* | *AvrLm3_K_01* | *AvrLm4-7_del* | *AvrLm5-9_0_A* | *AvrLm6_1_A* | *AvrLm11_0_A* | *AvrLmS_2_B* |
| WAC7750 | *AvrLm1_0_A* | *AvrLm2_2_A* | *AvrLm3_A_01* | *AvrLm4-7_0_A* | *AvrLm5-9_1_A* | *AvrLm6_1_A* | *AvrLm11_0_A* | *AvrLmS_2_B* |
| WAC7803 | *AvrLm1_del* | *AvrLm2_0_A* | *AvrLm3_K_01* | *AvrLm4-7_del* | *AvrLm5-9_3_A* | *AvrLm6_1_A* | *AvrLm11_0_A* | *AvrLmS_2_A* |
| X83-51 | *AvrLm1_del* | *AvrLm2_1_A* | *AvrLm3_R_01* | *AvrLm4-7_del* | *AvrLm5-9_0_A* | *AvrLm6_5_A* | *AvrLm11_del* | *AvrLmS_del* |
| HB10-19 | *AvrLm1_del* | *AvrLm2_X1* | *AvrLm3_del* | *AvrLm4-7_del* | *AvrLm5-9_del* | *AvrLm6_X1* | *AvrLm11_X1* | *AvrLmS_X1_A* |

[1] For more information on isolates origin, see Gautier et al, 2025

[2] Allele names as defined in Gautier et al., 2023. *del*, deleted allele.

**Supplementary Table S2. Characteristics of all plant DNA samples submitted to the MPSeqM tool**

| Sample name | Run | Sample type | Sampling season | French region | French department name and numer | Town and zip code | Rapeseed Variety‡ | *Rlm* gene in rapeseed variety | Number leaf spots :he pool |
|---|---|---|---|---|---|---|---|---|---|
| JAN_1 | RunA | environmental | 2019-20 | Centre-Val de Loire | Eure-et-Loir-28 | Janville-28310 | Westar | None | 32 |
| JAN_2 | RunA | environmental | 2019-20 | Centre-Val de Loire | Eure-et-Loir-28 | Janville-28310 | Westar | None | 32 |
| JAN_3_1 | RunA | environmental | 2019-20 | Centre-Val de Loire | Eure-et-Loir-28 | Janville-28310 | Westar | None | 32 |
| JAN_3_2 | RunA | environmental | 2019-20 | Centre-Val de Loire | Eure-et-Loir-28 | Janville-28310 | Westar | None | 32 |
| THI_B_1_1 | RunA | environmental | 2019-20 | Ile de France | Yvelines-78 | Thiverval-Grignon-78850 | Westar | None | 32 |
| THI_B_1_2 | RunA | environmental | 2019-20 | Ile de France | Yvelines-78 | Thiverval-Grignon-78850 | Westar | None | 32 |
| THI_B_2_1 | RunA | environmental | 2019-20 | Ile de France | Yvelines-78 | Thiverval-Grignon-78850 | Westar | None | 32 |
| THI_B_2_2 | RunA | environmental | 2019-20 | Ile de France | Yvelines-78 | Thiverval-Grignon-78850 | Westar | None | 32 |
| BOI_1_1 | RunA | environmental | 2019-20 | Centre-Val de Loire | Loir-et-Cher-41 | Boisseau-41290 | Westar | None | 32 |
| BOI_1_2 | RunA | environmental | 2019-20 | Centre-Val de Loire | Loir-et-Cher-41 | Boisseau-41290 | Westar | None | 32 |
| BOI_2_1 | RunA | environmental | 2019-20 | Centre-Val de Loire | Loir-et-Cher-41 | Boisseau-41290 | Westar | None | 32 |
| BOI_2_2 | RunA | environmental | 2019-20 | Centre-Val de Loire | Loir-et-Cher-41 | Boisseau-41290 | Westar | None | 32 |
| BOI_3_1 | RunA | environmental | 2019-20 | Centre-Val de Loire | Loir-et-Cher-41 | Boisseau-41290 | Westar | None | 32 |
| BOI_3_2 | RunA | environmental | 2019-20 | Centre-Val de Loire | Loir-et-Cher-41 | Boisseau-41290 | Westar | None | 32 |
| ERD_1 | RunA | environmental | 2019-20 | Pays de la Loire | Maine-et-Loire-49 | Erdre-en-Anjou-49220 | Westar | None | 32 |
| ERD_2 | RunA | environmental | 2019-20 | Pays de la Loire | Maine-et-Loire-49 | Erdre-en-Anjou-49220 | Westar | None | 32 |
| ERD_3 | RunA | environmental | 2019-20 | Pays de la Loire | Maine-et-Loire-49 | Erdre-en-Anjou-49220 | Westar | None | 32 |
| ERD_4 | RunA | environmental | 2019-20 | Pays de la Loire | Maine-et-Loire-49 | Erdre-en-Anjou-49220 | Westar | None | 32 |
| BLC_1 | RunA | environmental | 2019-20 | Centre-Val de Loire | Indre-36 | Bouges-le-Chateau-36110 | Westar | None | 33 |
| BLC_2 | RunA | environmental | 2019-20 | Centre-Val de Loire | Indre-36 | Bouges-le-Chateau-36110 | Westar | None | 32 |
| BLC_3 | RunA | environmental | 2019-20 | Centre-Val de Loire | Indre-36 | Bouges-le-Chateau-36110 | Westar | None | 32 |
| BLC_4 | RunA | environmental | 2019-20 | Centre-Val de Loire | Indre-36 | Bouges-le-Chateau-36110 | Westar | None | 32 |
| BLC_5 | RunA | environmental | 2019-20 | Centre-Val de Loire | Indre-36 | Bouges-le-Chateau-36110 | Westar | None | 32 |
| FRE_1 | RunA | environmental | 2019-20 | Hauts-de-France | Nord-59 | Frelinghien-59236 | Westar | None | 24 |

| | | | | | | | | | |
|---|---|---|---|---|---|---|---|---|---|
| FRE_2 | RunA | environmental | 2019-20 | Hauts-de-France | Nord-59 | Frelinghien-59236 | Westar | None | 32 |
| FRE_3 | RunA | environmental | 2019-20 | Hauts-de-France | Nord-59 | Frelinghien-59236 | Westar | None | 32 |
| RHE_2019_1_1 | RunA | environmental | 2019-20 | Bretagne | Ille-et-Vilaine-35 | Le-Rheu-35650 | Westar | None | 32 |
| RHE_2019_1_2 | RunA | environmental | 2019-20 | Bretagne | Ille-et-Vilaine-35 | Le-Rheu-35650 | Westar | None | 32 |
| RHE_2019_2_1 | RunA | environmental | 2019-20 | Bretagne | Ille-et-Vilaine-35 | Le-Rheu-35650 | Westar | None | 32 |
| RHE_2019_2_2 | RunA | environmental | 2019-20 | Bretagne | Ille-et-Vilaine-35 | Le-Rheu-35650 | Westar | None | 32 |
| RHE_2019_3_1 | RunA | environmental | 2019-20 | Bretagne | Ille-et-Vilaine-35 | Le-Rheu-35650 | Westar | None | 32 |
| RHE_2019_3_2 | RunA | environmental | 2019-20 | Bretagne | Ille-et-Vilaine-35 | Le-Rheu-35650 | Westar | None | 32 |
| THI_T_1_1 | RunA | environmental | 2019-20 | Ile de France | Yvelines-78 | Thiverval-Grignon-78850 | Westar | None | 32 |
| THI_T_1_2 | RunA | environmental | 2019-20 | Ile de France | Yvelines-78 | Thiverval-Grignon-78850 | Westar | None | 32 |
| THI_T_2_1 | RunA | environmental | 2019-20 | Ile de France | Yvelines-78 | Thiverval-Grignon-78850 | Westar | None | 32 |
| THI_T_2_2 | RunA | environmental | 2019-20 | Ile de France | Yvelines-78 | Thiverval-Grignon-78850 | Westar | None | 32 |
| COU_1_1 | RunA | environmental | 2019-20 | Centre-Val de Loire | Loiret-45 | Coullons-45720 | Westar | None | 32 |
| COU_1_2 | RunA | environmental | 2019-20 | Centre-Val de Loire | Loiret-45 | Coullons-45720 | Westar | None | 32 |
| COU_2_1 | RunA | environmental | 2019-20 | Centre-Val de Loire | Loiret-45 | Coullons-45720 | Westar | None | 32 |
| COU_2_2 | RunA | environmental | 2019-20 | Centre-Val de Loire | Loiret-45 | Coullons-45720 | Westar | None | 32 |
| COU_3_1 | RunA | environmental | 2019-20 | Centre-Val de Loire | Loiret-45 | Coullons-45720 | Westar | None | 32 |
| COU_3_2 | RunA | environmental | 2019-20 | Centre-Val de Loire | Loiret-45 | Coullons-45720 | Westar | None | 32 |
| SPT_3 | RunB | environmental | 2022-23 | Hauts-de-France | Pas-de-Calais-62 | Saint-Pol-sur-Ternoise-62130 | Marcopolos | Rlm9 | 32 |
| SPT_1_1 | RunB | environmental | 2022-23 | Hauts-de-France | Pas-de-Calais-62 | Saint-Pol-sur-Ternoise-62130 | Marcopolos | Rlm9 | 32 |
| SPT_1_2 | RunB | environmental | 2022-23 | Hauts-de-France | Pas-de-Calais-62 | Saint-Pol-sur-Ternoise-62130 | Marcopolos | Rlm9 | 32 |
| SPT_2_1 | RunB | environmental | 2022-23 | Hauts-de-France | Pas-de-Calais-62 | Saint-Pol-sur-Ternoise-62130 | Marcopolos | Rlm9 | 32 |
| SPT_2_2 | RunB | environmental | 2022-23 | Hauts-de-France | Pas-de-Calais-62 | Saint-Pol-sur-Ternoise-62130 | Marcopolos | Rlm9 | 32 |
| TOU_1_1 | RunB | environmental | 2022-23 | Centre-Val de Loire | Eure-et-Loir-28 | Toury-28310 | Drakkar | None | 32 |
| TOU_1_2 | RunB | environmental | 2022-23 | Centre-Val de Loire | Eure-et-Loir-28 | Toury-28310 | Drakkar | None | 31 |
| TOU_5 | RunB | environmental | 2022-23 | Centre-Val de Loire | Eure-et-Loir-28 | Toury-28310 | Drakkar | None | 33 |
| TOU_2 | RunB | environmental | 2022-23 | Centre-Val de Loire | Eure-et-Loir-28 | Toury-28310 | Drakkar | None | 32 |
| TOU_3 | RunB | environmental | 2022-23 | Centre-Val de Loire | Eure-et-Loir-28 | Toury-28310 | Drakkar | None | 32 |
| TOU_4 | RunB | environmental | 2022-23 | Centre-Val de Loire | Eure-et-Loir-28 | Toury-28310 | Drakkar | None | 32 |
| *BOI_2_1_B*[§] | RunB | environmental | 2019-20 | Centre-Val de Loire | Loir-et-Cher-41 | Boisseau-41290 | Westar | None | 32 |

| | | | | | | | | | |
|---|---|---|---|---|---|---|---|---|---|
| *BOI_3_2_B* | RunB | environmental | 2019-20 | Centre-Val de Loire | Loir-et-Cher-41 | Boisseau-41290 | Westar | None | 32 |
| VIL_1_1 | RunB | environmental | 2022-23 | Centre-Val de Loire | Loir-et-Cher-41 | Villeneuve-Frouville-41290 | Drakkar | None | 32 |
| VIL_1_2 | RunB | environmental | 2022-23 | Centre-Val de Loire | Loir-et-Cher-41 | Villeneuve-Frouville-41290 | Drakkar | None | 32 |
| VIL_4 | RunB | environmental | 2022-23 | Centre-Val de Loire | Loir-et-Cher-41 | Villeneuve-Frouville-41290 | Drakkar | None | 32 |
| VIL_2_1 | RunB | environmental | 2022-23 | Centre-Val de Loire | Loir-et-Cher-41 | Villeneuve-Frouville-41290 | Drakkar | None | 32 |
| VIL_2_2 | RunB | environmental | 2022-23 | Centre-Val de Loire | Loir-et-Cher-41 | Villeneuve-Frouville-41290 | Drakkar | None | 32 |
| VIL_3 | RunB | environmental | 2022-23 | Centre-Val de Loire | Loir-et-Cher-41 | Villeneuve-Frouville-41290 | Drakkar | None | 32 |
| RHE_2022_1 | RunB | environmental | 2022-23 | Bretagne | Ille-et-Vilaine-35 | Le-Rheu-35650 | Darmor | Rlm9 | 24 |
| RHE_2022_2 | RunB | environmental | 2022-23 | Bretagne | Ille-et-Vilaine-35 | Le-Rheu-35650 | Drakkar | None | 32 |
| RHE_2022_3 | RunB | environmental | 2022-23 | Bretagne | Ille-et-Vilaine-35 | Le-Rheu-35650 | Drakkar | None | 32 |
| RHE_2022_4 | RunB | environmental | 2022-23 | Bretagne | Ille-et-Vilaine-35 | Le-Rheu-35650 | Drakkar & Marcopolos ¶ | None & *Rlm9* | 32 |
| RHE_2022_5_1 | RunB | environmental | 2022-23 | Bretagne | Ille-et-Vilaine-35 | Le-Rheu-35650 | MarcoPolo | Rlm9 | 32 |
| RHE_2022_5_2 | RunB | environmental | 2022-23 | Bretagne | Ille-et-Vilaine-35 | Le-Rheu-35650 | MarcoPolo | Rlm9 | 32 |
| ENN_1 | RunB | environmental | 2022-23 | Hauts-de-France | Nord-59 | Ennevelin-59710 | Drakkar & Marcopolos | None & *Rlm9* | 32 |
| ENN_2_1 | RunB | environmental | 2022-23 | Hauts-de-France | Nord-59 | Ennevelin-59710 | Drakkar | None | 32 |
| ENN_2_2 | RunB | environmental | 2022-23 | Hauts-de-France | Nord-59 | Ennevelin-59710 | Drakkar | None | 32 |
| ENN_3_1 | RunB | environmental | 2022-23 | Hauts-de-France | Nord-59 | Ennevelin-59710 | Marcopolos | *Rlm9* | 32 |
| ENN_3_2 | RunB | environmental | 2022-23 | Hauts-de-France | Nord-59 | Ennevelin-59710 | Marcopolos | *Rlm9* | 32 |
| ENN_3_3 | RunB | environmental | 2022-23 | Hauts-de-France | Nord-59 | Ennevelin-59710 | Marcopolos | *Rlm9* | 32 |
| *RHE_2019_1_1_B* | RunB | environmental | 2019-20 | Bretagne | Ille-et-Vilaine-35 | Le-Rheu-35650 | Westar | None | 32 |
| TER_1 | RunB | environmental | 2022-23 | Centre-Val de Loire | Eure-et-Loir-28 | Terminiers-28140 | Drakkar | None | 31 |
| TER_2_1 | RunB | environmental | 2022-23 | Centre-Val de Loire | Eure-et-Loir-28 | Terminiers-28140 | Marcopolos | *Rlm9* | 32 |
| TER_2_2 | RunB | environmental | 2022-23 | Centre-Val de Loire | Eure-et-Loir-28 | Terminiers-28140 | Marcopolos | *Rlm9* | 32 |
| TER_2_3 | RunB | environmental | 2022-23 | Centre-Val de Loire | Eure-et-Loir-28 | Terminiers-28140 | Marcopolos | *Rlm9* | 32 |
| TER_3 | RunB | environmental | 2022-23 | Centre-Val de Loire | Eure-et-Loir-28 | Terminiers-28140 | Marcopolos | *Rlm9* | 32 |
| TER_4 | RunB | environmental | 2022-23 | Centre-Val de Loire | Eure-et-Loir-28 | Terminiers-28140 | Marcopolos | *Rlm9* | 32 |
| BRE_1 | RunB | environmental | 2022-23 | Bourgogne-Franche-Comté | Cote-d'Or-21 | Bretenière-21110 | Drakkar | None | 32 |
| BRE_2 | RunB | environmental | 2022-23 | Bourgogne-Franche-Comté | Cote-d'Or-21 | Bretenière-21110 | Drakkar & Marcopolos | None & *Rlm9* | 34 |

| | | | | | | | | | |
|---|---|---|---|---|---|---|---|---|---|
| BRE_3 | RunB | environmental | 2022-23 | Bourgogne-Franche-Comté | Cote-d'Or-21 | Bretenière-21110 | MarcoPolo | *Rlm9* | 32 |
| *COU_3_2_B* | RunB | environmental | 2019-20 | Centre-Val de Loire | Loiret-45 | Coullons-45720 | Westar | None | 32 |
| VERS_1 | RunB | environmental | 2022-23 | Ile de France | Yvelines-78 | Versailles-78000 | Drakkar | None | 34 |
| VERS_2 | RunB | environmental | 2022-23 | Ile de France | Yvelines-78 | Versailles-78000 | Drakkar & Marcopolos | None & *Rlm9* | 31 |
| VERS_3_1 | RunB | environmental | 2022-23 | Ile de France | Yvelines-78 | Versailles-78000 | Marcopolos | *Rlm9* | 32 |
| VERS_3_2 | RunB | environmental | 2022-23 | Ile de France | Yvelines-78 | Versailles-78000 | Marcopolos | *Rlm9* | 32 |
| VERS_3_3 | RunB | environmental | 2022-23 | Ile de France | Yvelines-78 | Versailles-78000 | Marcopolos | *Rlm9* | 32 |
| VERS_3_4 | RunB | environmental | 2022-23 | Ile de France | Yvelines-78 | Versailles-78000 | Marcopolos | *Rlm9* | 32 |

†, not applicable
‡, trap cultivar from which the leaf spots were collected

§, samples in italics were analized in RunA, but also used as controls for reproducibility in RunB
¶, leaf spots from different varieties were pooled when insufficient infected leaves were obtained with only one variety

Supplementary Table S3. Characteristics of the eight *AvrLm* and the *Actin* gene regions amplified with MiSeq primers

| Gene | Gene size (bp) for JN3 | No. of known polymorphic sites[a] | Forward Primer for MiSeq (5'-> 3') [b] | Reverse Primer for MiSeq (5'-> 3') [c] | Amplicon size (bp)[d] | Length of the analyzed part of the gene (bp) | No. (%) of known polymorphic sites in the amplicon[e] |
|---|---|---|---|---|---|---|---|
| *AvrLm1* | 667 | 3 | TCCGTCCAGATAAGCCCTGT | CACGAATGATGCACCAGAGGC | 455 | 358 | 3 (100%) |
| *AvrLm2* | 699 | 5 | CCCAGCAACAATTTTCGGAACTCC | TTCAACTGTGTATTGGCCCTGG | 433 | 331 | 5 (100%) |
| *AvrLm3* | 539 | 25 | CCAAATCTTGTTTCTATTATTTACCGCAT | GATGATCGCAGTGTACCTCGTCTTT | 511 | 401 | 21 (84%) |
| *AvrLm4-7* | 478 | 13 | ACTCGCTCTCTCTATCCCTACA | CAACCACGAGTCCTTGCTTTCA | 495 | 395 | 13 (100%) |
| *AvrLm5-9* | 426 | 6 | ACAACCACTCTTCTTCACAGTTT | TTAGAGATAACTAAGCTCCAATCCG | 486 | 382 | 5 (83.3%) |
| *AvrLm6*[f] | 581 | 3 | GCACGTTCCAAAGCCCTATTTA | ATCGTCATGGGCTTGGATGTTA | 506 | 406 | 2 (66.7%) |
| *AvrLm6*[g] | 581 | 3 | ACTACCACGATTAGTCAGCC | ACAGATAACCAAGCGCTGAC | 516 | 420 | 3 (100%) |
| *AvrLm11* | 365 | 3 | GCTTCATCATGCGTTTCTTGCT | CCACAGCACACTATTCACTTTAGC | 504 | 402 | 3 (100%) |
| *AvrLmS-Lep2* | 426 | 15 | GGCCAACTTTATCATGCTCTACCT | GGCAACTGATGGGACCGCTC | 487 | 387 | 15 (100%) |
| *Actin partial (512F/CA14R)* | 374 | - | ATGTGCAAGGCCGGTTTCG | GCCAGATCTTCTCCATGTCATCCCAGTT | 430 | 327 | - |

[a] From Start to Stop, data from Gautier et al. (2023)

[b] all forward primers were tagged in 5' with the MiSeq specific sequence CTTTCCCTACACGACGCTCTTCCGATCT

[c] all reverse primers were tagged in 5' with the MiSeq specific sequence GGAGTTCAGACGTGTGCTCTTCCGATCT

[d] including the 5' and 3' tags and for JN3 isolate

[e] except RIP mutations; known mutations as described by Gautier et al. (2023).

[f] for the 2019 sampling analysis

[g] for the 2022 sampling analysis

**Figure S4. Virulence, avirulence, and heterogeneous phenotypes of Leptosphaeria maculans isolates inoculated on Darmor-Rlm11 genotype.** Panels (A) and (B) illustrate the heterogeneous phenotype (H) of L. maculans isolates on the oilseed rape genotype, Darmor-Rlm11. In contrast, panel (C) illustrates typical symptoms of avirulence (A, left part of the cotyledons) and virulence (V, right part of the cotyledons) of L. maculans control isolates, on Darmor-Rlm11. Each panel represents 8 replicates of the same inoculation experiment, with two isolates per cotyledon, as follows; A, isolates CP.22.805 (H) on the left part of the cotyledons and CP.22.806 (H) on the right part of the same cotyledon; B, CP.22.878 (H) on the left part of the cotyledons and CP.22.879 (H) on the right part of the same cotyledon, and C, JN2 (A) on the left part of the cotyledons and X83.51 (V) on the right part of the same cotyledon. Pictures were taken 15 days after inoculation.

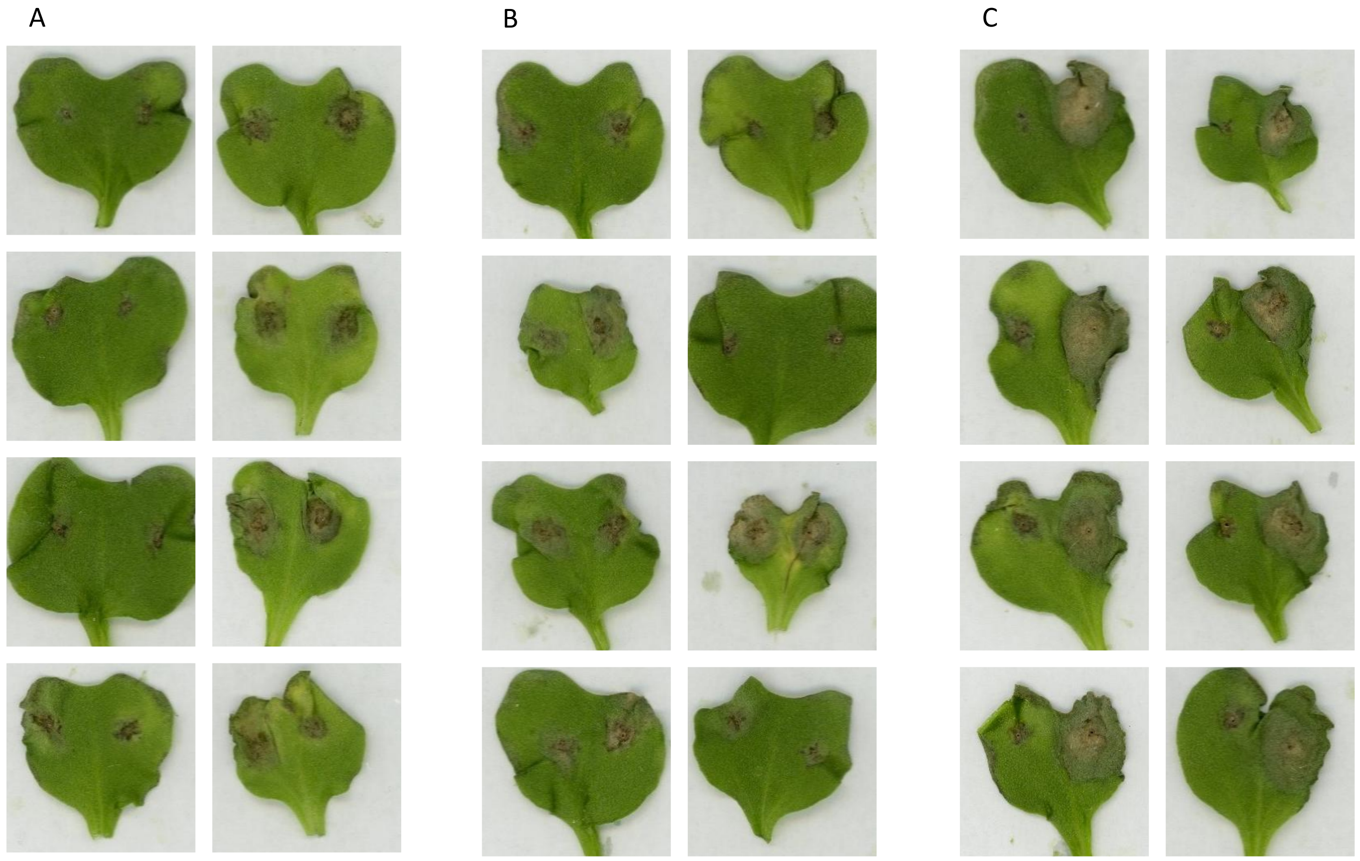

**Supplementary Figure S5. Regional distribution of *Leptosphaeria maculans* avirulence alleles for five genes during two growing seasons.** A and B, *AvrLm2*; C and D, *AvrLm3*; E and F, *AvrLm5-9*; G and H, *AvrLm6*; I and J, *AvrLm11*. A, C, E, G and I, 2019-20 survey; B, D, F, H and J, 2022-23 survey.

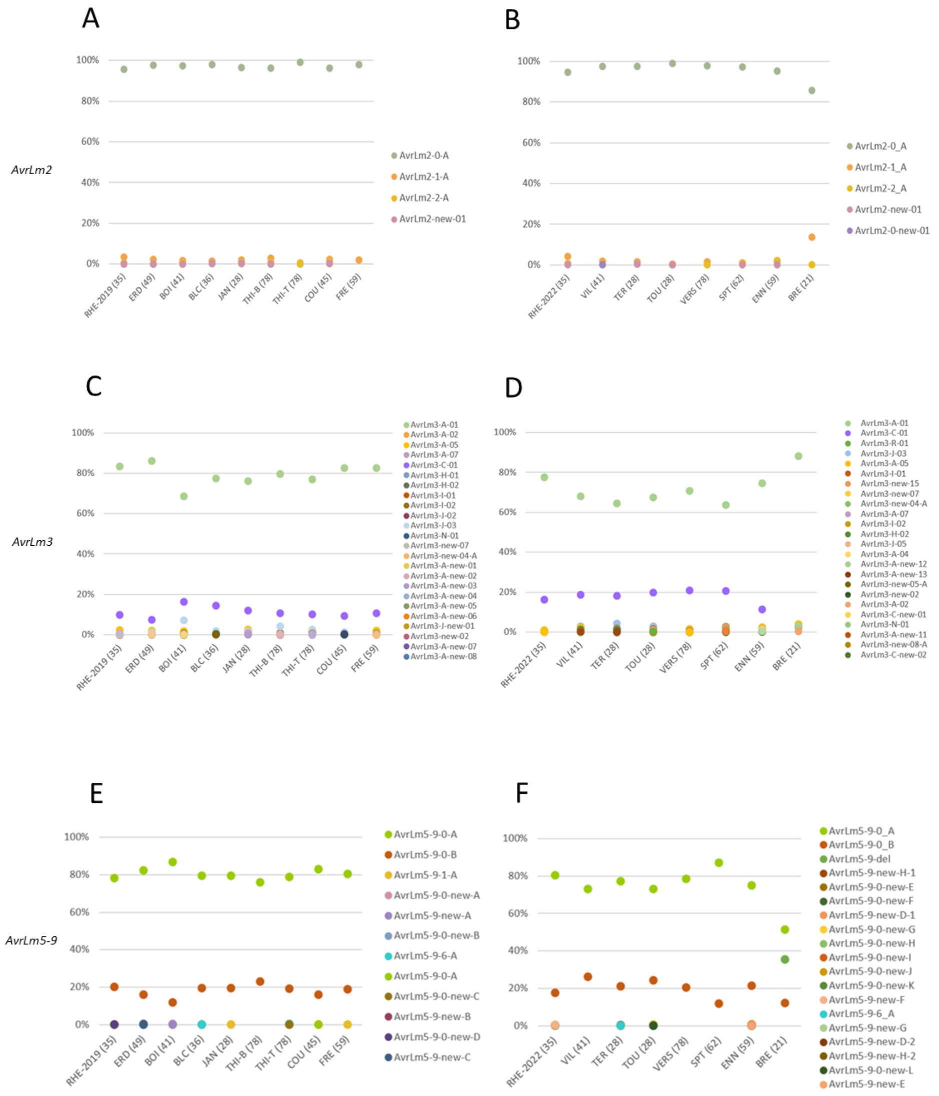

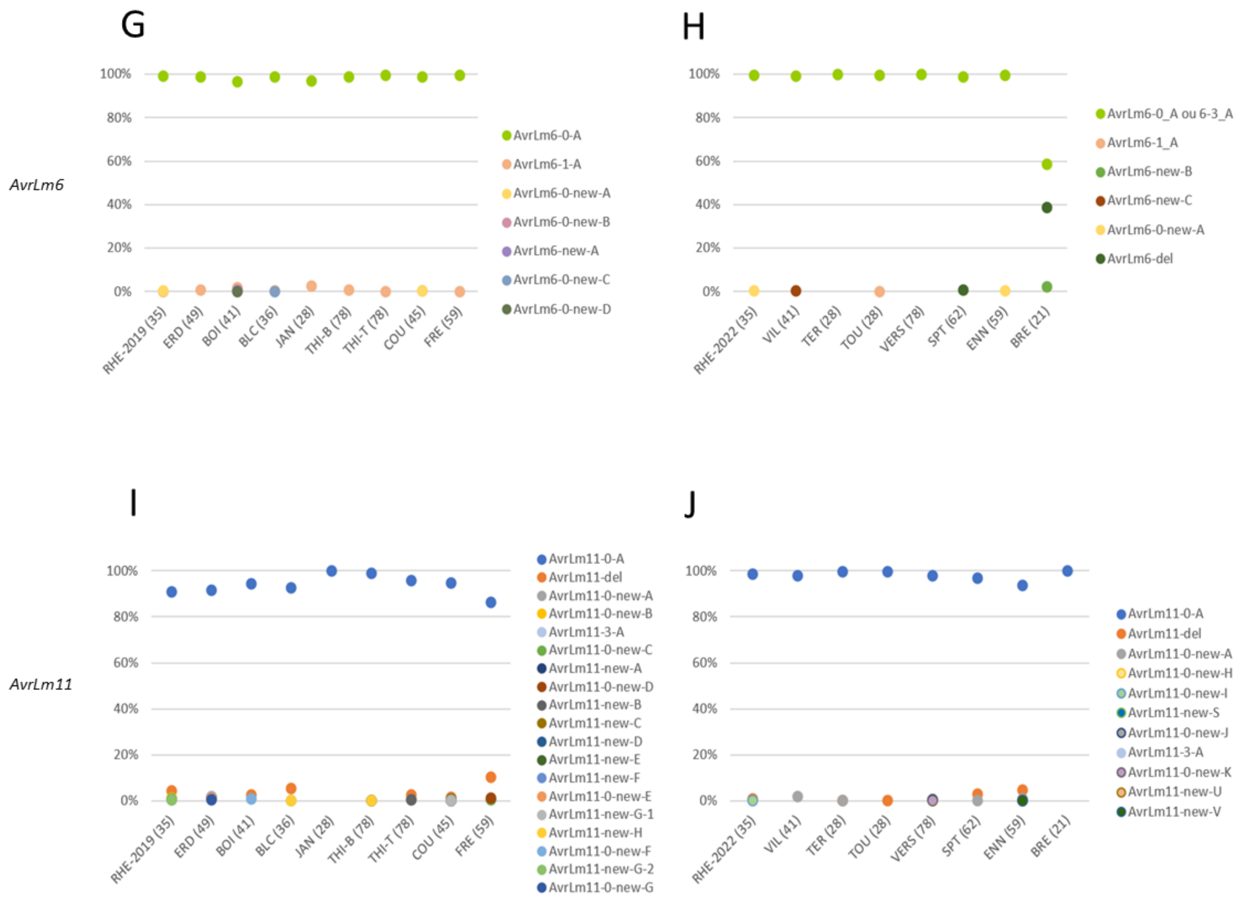
G
H
I
J
AvrLm6
AvrLm11
100%
80%
60%
40%
20%
0%
RHE-2019 (35)
ERD (49)
BOI (41)
BLC (36)
JAN (28)
THI-B (78)
THI-T (78)
COU (45)
FRE (59)
RHE-2022 (35)
VIL (41)
TER (28)
TOU (28)
VERS (78)
SPT (62)
ENN (59)
BRE (21)
AvrLm6-0-A
AvrLm6-1-A
AvrLm6-0-new-A
AvrLm6-0-new-B
AvrLm6-new-A
AvrLm6-0-new-C
AvrLm6-0-new-D
AvrLm6-0_A ou 6-3_A
AvrLm6-1_A
AvrLm6-new-B
AvrLm6-new-C
AvrLm6-0-new-A
AvrLm6-del
AvrLm11-0-A
AvrLm11-del
AvrLm11-0-new-A
AvrLm11-0-new-B
AvrLm11-3-A
AvrLm11-0-new-C
AvrLm11-new-A
AvrLm11-0-new-D
AvrLm11-new-B
AvrLm11-new-C
AvrLm11-new-D
AvrLm11-new-E
AvrLm11-new-F
AvrLm11-0-new-E
AvrLm11-new-G-1
AvrLm11-new-H
AvrLm11-0-new-F
AvrLm11-new-G-2
AvrLm11-0-new-G
AvrLm11-0-A
AvrLm11-del
AvrLm11-0-new-A
AvrLm11-0-new-H
AvrLm11-0-new-I
AvrLm11-new-S
AvrLm11-0-new-J
AvrLm11-3-A
AvrLm11-0-new-K
AvrLm11-new-U
AvrLm11-new-V

**Supplementary file S6. Percentages of RIPped MPSeqM ASV, leading to virulent alleles, for the two sampling seasons 2019-20 and 2022-23.** The tables indicate the percentages per avirulence gene, sampling site and season. Sampling sites ID are described in Supplementary Table S2. The graph represents the mean and standard error values over all sites per avirulence gene and growing season.

| Sampling year 2019-2020 | | | | | | | | | | | |
|---|---|---|---|---|---|---|---|---|---|---|---|
| *Avrlm gene* | Sampling site | | | | | | | | | Mean | Standard error |
| | RHE-2019 (35) | ERD (49) | BOI (41) | BLC (36) | JAN (28) | THI-B (78) | THI-T (78) | COU (45) | FRE (59) | | |
| *AvrLm1* | 0,00% | 0,00% | 0,00% | 0,00% | 0,00% | 0,00% | 0,00% | 0,25% | 0,00% | 0,03% | 0,08% |
| *AvrLm2* | 0,00% | 0,00% | 0,00% | 0,00% | 0,00% | 0,00% | 0,00% | 0,00% | 0,00% | 0,00% | 0,00% |
| *AvrLm3* | 0,00% | 0,00% | 0,00% | 0,00% | 0,00% | 0,00% | 0,00% | 0,00% | 0,00% | 0,00% | 0,00% |
| *AvrLm4-7* | 2,40% | 0,85% | 1,61% | 2,94% | 2,81% | 2,11% | 2,25% | 3,43% | 1,32% | 2,19% | 0,82% |
| *AvrLm5-9* | 0,00% | 0,00% | 0,00% | 0,00% | 0,00% | 0,00% | 0,00% | 0,00% | 0,00% | 0,00% | 0,00% |
| *AvrLm6* | 0,00% | 0,00% | 0,00% | 0,00% | 0,00% | 0,00% | 0,00% | 0,00% | 0,00% | 0,00% | 0,00% |
| *AvrLm11* | 0,00% | 0,00% | 0,00% | 0,00% | 0,00% | 0,00% | 0,00% | 0,20% | 0,00% | 0,02% | 0,07% |
| *AvrLmS* | 0,00% | 0,00% | 0,00% | 0,00% | 0,00% | 0,00% | 0,00% | 0,00% | 0,00% | 0,00% | 0,00% |
| Mean | 0,30% | 0,11% | 0,20% | 0,37% | 0,35% | 0,26% | 0,28% | 0,49% | 0,17% | 0,28% | 0,11% |

| Sampling year 2022-2023 | | | | | | | | | | |
|---|---|---|---|---|---|---|---|---|---|---|
| *AvrLm gene* | Sampling site | | | | | | | | Mean | Standard error |
| | RHE-2022 (35) | VIL (41) | TER (28) | TOU (28) | VERS (78) | SPT (62) | ENN (59) | BRE (21) | | |
| *AvrLm1* | 0,00% | 0,00% | 0,00% | 0,00% | 0,06% | 0,00% | 0,00% | 0,00% | 0,01% | 0,02% |
| *AvrLm2* | 0,00% | 0,00% | 0,00% | 0,00% | 0,00% | 0,00% | 0,00% | 0,00% | 0,00% | 0,00% |
| *AvrLm3* | 0,00% | 0,00% | 0,00% | 0,00% | 0,00% | 0,00% | 0,00% | 0,00% | 0,00% | 0,00% |
| *AvrLm4-7* | 1,07% | 0,50% | 0,80% | 1,59% | 0,00% | 0,00% | 0,40% | 3,44% | 0,98% | 1,13% |
| *AvrLm5-9* | 0,00% | 0,00% | 0,00% | 0,00% | 0,00% | 0,00% | 0,00% | 0,00% | 0,00% | 0,00% |
| *AvrLm6* | 0,00% | 0,00% | 0,00% | 0,00% | 0,00% | 0,00% | 0,00% | 0,00% | 0,00% | 0,00% |
| *AvrLm11* | 0,00% | 0,00% | 0,00% | 0,00% | 0,00% | 0,00% | 0,00% | 0,00% | 0,00% | 0,00% |
| *AvrLmS* | 0,00% | 0,00% | 0,00% | 0,00% | 0,00% | 0,00% | 0,00% | 0,00% | 0,00% | 0,00% |
| *Mean* | 0,13% | 0,06% | 0,10% | 0,20% | 0,01% | 0,00% | 0,05% | 0,43% | 0,12% | 0,14% |

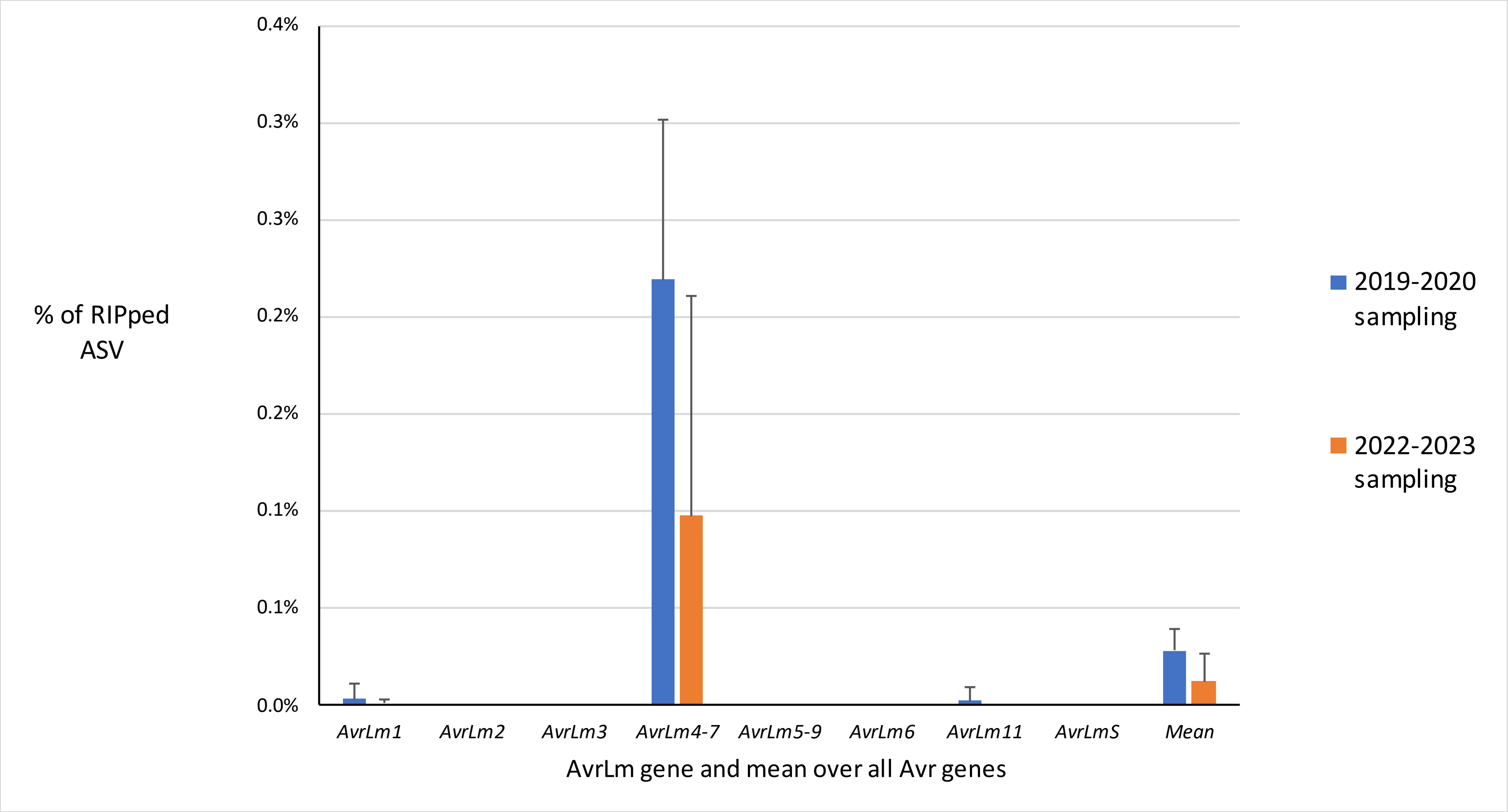

0.4%
0.3%
0.3%
0.2%
0.2%
0.1%
0.1%
0.0%
% of RIPped ASV
AvrLm1
AvrLm2
AvrLm3
AvrLm4-7
AvrLm5-9
AvrLm6
AvrLm11
AvrLmS
Mean
AvrLm gene and mean over all Avr genes
2019-2020 sampling
2022-2023 sampling

**Supplementary Table S7. Assignation of *L. maculans* avirulence alleles to a corresponding phenotype on the corresponding resistance gene**

| Gene | Genotype | Phenotype | Masking effect[a] |
|---|---|---|---|
| *AvrLm1* | *AvrLm1-0-A* | A[&]1 | |
| *AvrLm1* | *AvrLm1-rip-new-A* | a1 | |
| *AvrLm1* | *AvrLm1-new-A* | a1 | |
| *AvrLm1* | *AvrLm1-0-new-A* | A1 | |
| *AvrLm1* | *AvrLm1-rip-new-B* | a1 | |
| *AvrLm1* | *AvrLm1-del* | a1 | |
| *AvrLm2* | *AvrLm2-0-A* | a2 | |
| *AvrLm2* | *AvrLm2-1-A* | a2 | |
| *AvrLm2* | *AvrLm2-2-A* | A2 | |
| *AvrLm2* | *AvrLm2-5-A* | A2 | |
| *AvrLm2* | *AvrLm2-0-new-01* | a2 | |
| *AvrLm3* | *AvrLm3-A-01* | A3 | |
| *AvrLm3* | *AvrLm3-C-01* | A3 | |
| *AvrLm3* | *AvrLm3-J-03* | a3 | |
| *AvrLm3* | *AvrLm3-A-05* | A3 | |
| *AvrLm3* | *AvrLm3-I-01* | a3 | |
| *AvrLm3* | *AvrLm3-H-02* | a3 | |
| *AvrLm3* | *AvrLm3-I-02* | a3 | |
| *AvrLm3* | *AvrLm3-A-02* | A3 | |
| *AvrLm3* | *AvrLm3-A-07* | A3 | |
| *AvrLm3* | *AvrLm3-J-05-K-02-W-01* | a3 | |
| *AvrLm3* | *AvrLm3-A-16* | A3 | |
| *AvrLm3* | *AvrLm3-J-02* | a3 | |
| *AvrLm3* | *AvrLm3-N-01* | a3 | |
| *AvrLm3* | *AvrLm3-R-01* | a3 | |

| | | |
|---|---|---|
| *AvrLm3* | *AvrLm3-A-04* | A3 |
| *AvrLm3* | *AvrLm3-new-04-A* | A3 |
| *AvrLm3* | *AvrLm3-A-new-01* | A3 |
| *AvrLm3* | *AvrLm3-A-new-02* | A3 |
| *AvrLm3* | *AvrLm3-A-new-03* | A3 |
| *AvrLm3* | *AvrLm3-A-new-04* | A3 |
| *AvrLm3* | *AvrLm3-A-new-05* | A3 |
| *AvrLm3* | *AvrLm3-A-new-06* | A3 |
| *AvrLm3* | *AvrLm3-new-02* | A3 |
| *AvrLm3* | *AvrLm3-A-new-07* | A3 |
| *AvrLm3* | *AvrLm3-A-new-08* | A3 |
| *AvrLm3* | *AvrLm3-A-new-09* | A3 |
| *AvrLm3* | *AvrLm3-new-01* | A3 |
| *AvrLm3* | *AvrLm3-A-new-10* | A3 |
| *AvrLm3* | *AvrLm3-new-05-A* | A3 |
| *AvrLm3* | *AvrLm3-new-03* | A3 |
| *AvrLm3* | *AvrLm3-A-new-11* | A3 |
| *AvrLm3* | *AvrLm3-new-10* | a3 |
| *AvrLm3* | *AvrLm3-new-04-B* | A3 |
| *AvrLm3* | *AvrLm3-new-06-A* | A3 |
| *AvrLm3* | *AvrLm3-new-08-A* | NA |
| *AvrLm3* | *AvrLm3-N-new-01* | a3 |
| *AvrLm3* | *AvrLm3-new-09* | NA |
| *AvrLm3* | *AvrLm3-new-08-B* | NA |
| *AvrLm3* | *AvrLm3-N-new-02* | a3 |
| *AvrLm3* | *AvrLm3-new-05-B* | A3 |
| *AvrLm3* | *AvrLm3-new-06-B* | A3 |
| *AvrLm3* | *AvrLm3-new-15* | NA |
| *AvrLm3* | *AvrLm3-A-new-12* | A3 |
| *AvrLm3* | *AvrLm3-A-new-13* | A3 |

| | | | |
|---|---|---|---|
| *AvrLm3* | *AvrLm3-C-new-01* | A3 | |
| *AvrLm3* | *AvrLm3-C-new-02* | A3 | |
| *AvrLm3* | *AvrLm3-A-new-14* | A3 | |
| *AvrLm3* | *AvrLm3-A-new-15* | A3 | |
| *AvrLm3* | *AvrLm3-A-new-16* | A3 | |
| *AvrLm3* | *AvrLm3-new-01-B* | A3 | |
| *AvrLm3* | *AvrLm3-new-12* | A3 | |
| *AvrLm3* | *AvrLm3-C-new-03* | A3 | |
| *AvrLm3* | *AvrLm3-J-new-02* | a3 | |
| *AvrLm3* | *AvrLm3-new-13* | A3 | |
| *AvrLm3* | *AvrLm3-N-new-03* | a3 | |
| *AvrLm3* | *AvrLm3-new-16* | NA | |
| *AvrLm3* | *AvrLm3-A-new-17* | A3 | |
| *AvrLm3* | *AvrLm3-new-17* | a3 | |
| *AvrLm3* | *AvrLm3-new-11* | A3 | |
| *AvrLm3* | *AvrLm3-new-08-C* | NA | |
| *AvrLm3* | *AvrLm3-new-14* | A3 | |
| *AvrLm4-7* | *AvrLm4-7-0-A* | A4A7[b] | yes |
| *AvrLm4-7* | *AvrLm4-7-1-A* | a4A7 | yes |
| *AvrLm4-7* | *AvrLm4-7-2-A* | a4A7 | yes |
| *AvrLm4-7* | *AvrLm4-7-5-A* | a4A7 | yes |
| *AvrLm4-7* | *AvrLm4-7-6-A* | a4a7 | yes |
| *AvrLm4-7* | *AvrLm4-7-7-A* | a4a7 | yes |
| *AvrLm4-7* | *AvrLm4-7-8-A* | a4a7 | yes |
| *AvrLm4-7* | *AvrLm4-7-9-A* | a4a7 | yes |
| *AvrLm4-7* | *AvrLm4-7-10-A* | a4a7 | yes |
| *AvrLm4-7* | *AvrLm4-7-13-A* | a4a7 | yes |
| *AvrLm4-7* | *AvrLm4-7-14-A* | a4a7 | yes |
| *AvrLm4-7* | *AvrLm4-7-15-A* | a4a7 | yes |
| *AvrLm4-7* | *AvrLm4-7-16-A* | a4a7 | yes |

| | | | |
|---|---|---|---|
| *AvrLm4-7* | *AvrLm4-7-18-A* | a4a7 | yes |
| *AvrLm4-7* | *AvrLm4-7-21-A* | a4a7 | yes |
| *AvrLm4-7* | *AvrLm4-7-22-A* | a4a7 | yes |
| *AvrLm4-7* | *AvrLm4-7-22-B* | a4a7 | yes |
| *AvrLm4-7* | *AvrLm4-7-23-A* | a4a7 | yes |
| *AvrLm4-7* | *AvrLm4-7-24-A* | a4a7 | yes |
| *AvrLm4-7* | *AvrLm4-7-25-A* | a4a7 | yes |
| *AvrLm4-7* | *AvrLm4-7-26-A* | a4a7 | yes |
| *AvrLm4-7* | *AvrLm4-7-29-A* | a4A7 | yes |
| *AvrLm4-7* | *AvrLm4-7-34-A* | a4A7 | yes |
| *AvrLm4-7* | *AvrLm4-7-39-A* | a4A7 | yes |
| *AvrLm4-7* | *AvrLm4-7-40-A* | a4a7 | yes |
| *AvrLm4-7* | *AvrLm4-7-42-A* | a4a7 | yes |
| *AvrLm4-7* | *AvrLm4-7-del-AA-1-A* | a4a7 | no |
| *AvrLm4-7* | *AvrLm4-7-del-AA-2-A* | a4a7 | no |
| *AvrLm4-7* | *AvrLm4-7-rip-008* | a4a7 | no |
| *AvrLm4-7* | *AvrLm4-7-rip-009* | a4a7 | no |
| *AvrLm4-7* | *AvrLm4-7-del* | a4a7 | no |
| *AvrLm4-7* | *AvrLm4-7-new-A* | a4A7 | yes |
| *AvrLm4-7* | *AvrLm4-7-new-B* | NA | NK |
| *AvrLm4-7* | *AvrLm4-7-new-D* | NA | NK |
| *AvrLm4-7* | *AvrLm4-7-new-F* | A4A7 | yes |
| *AvrLm4-7* | *AvrLm4-7-new-G-1* | A4A7 | yes |
| *AvrLm4-7* | *AvrLm4-7-new-H* | NA | NK |
| *AvrLm4-7* | *AvrLm4-7-new-I-1* | NA | NK |
| *AvrLm4-7* | *AvrLm4-7-new-I-2* | NA | NK |
| *AvrLm4-7* | *AvrLm4-7-new-J* | NA | NK |
| *AvrLm4-7* | *AvrLm4-7-new-K-1* | NA | NK |
| *AvrLm4-7* | *AvrLm4-7-new-K-2* | NA | NK |
| *AvrLm4-7* | *AvrLm4-7-new-L* | NA | NK |

| | | | |
|---|---|---|---|
| *AvrLm4-7* | *AvrLm4-7-new-M* | NA | NK |
| *AvrLm4-7* | *AvrLm4-7-new-N* | NA | NK |
| *AvrLm4-7* | *AvrLm4-7-new-O* | a4A7 | yes |
| *AvrLm4-7* | *AvrLm4-7-new-P* | a4a7 | yes |
| *AvrLm4-7* | *AvrLm4-7-new-Q* | a4a7 | no |
| *AvrLm4-7* | *AvrLm4-7-new-R* | NA | NK |
| *AvrLm4-7* | *AvrLm4-7-new-S* | NA | NK |
| *AvrLm4-7* | *AvrLm4-7-new-U* | NA | NK |
| *AvrLm4-7* | *AvrLm4-7-new-V* | NA | NK |
| *AvrLm4-7* | *AvrLm4-7-new-W* | NA | NK |
| *AvrLm4-7* | *AvrLm4-7-new-X* | a4A7 | yes |
| *AvrLm4-7* | *AvrLm4-7-new-Y* | NA | NK |
| *AvrLm4-7* | *AvrLm4-7-new-Z* | NA | NK |
| *AvrLm4-7* | *AvrLm4-7-new-AA-1* | NA | NK |
| *AvrLm4-7* | *AvrLm4-7-new-AA-2* | NA | NK |
| *AvrLm4-7* | *AvrLm4-7-new-AB* | NA | NK |
| *AvrLm4-7* | *AvrLm4-7-new-AC* | NA | NK |
| *AvrLm4-7* | *AvrLm4-7-new-AD* | NA | NK |
| *AvrLm4-7* | *AvrLm4-7-new-AE* | NA | NK |
| *AvrLm4-7* | *AvrLm4-7-new-AG* | NA | NK |
| *AvrLm4-7* | *AvrLm4-7-new-AH* | a4a7 | no |
| *AvrLm4-7* | *AvrLm4-7-new-AI* | NA | NK |
| *AvrLm4-7* | *AvrLm4-7-new-AK* | a4a7 | no |
| *AvrLm4-7* | *AvrLm4-7-new-AL* | NA | NK |
| *AvrLm4-7* | *AvrLm4-7-new-AM* | NA | NK |
| *AvrLm4-7* | *AvrLm4-7-new-AN* | NA | NK |
| *AvrLm4-7* | *AvrLm4-7-new-AO* | NA | NK |
| *AvrLm4-7* | *AvrLm4-7-new-AP* | NA | NK |
| *AvrLm4-7* | *AvrLm4-7-new-AQ* | a4a7 | no |
| *AvrLm4-7* | *AvrLm4-7-new-AR* | a4a7 | no |

| | | | |
|---|---|---|---|
| *AvrLm4-7* | *AvrLm4-7-new-AS* | NA | NK |
| *AvrLm4-7* | *AvrLm4-7-new-AT* | a4a7 | no |
| *AvrLm4-7* | *AvrLm4-7-new-AU* | a4a7 | no |
| *AvrLm4-7* | *AvrLm4-7-new-del-A-1* | a4a7 | no |
| *AvrLm4-7* | *AvrLm4-7-new-del-A-2* | a4a7 | no |
| *AvrLm4-7* | *AvrLm4-7-new-del-AG* | a4a7 | no |
| *AvrLm4-7* | *AvrLm4-7-new-del-G* | a4a7 | no |
| *AvrLm4-7* | *AvrLm4-7-new-in-TC* | a4a7 | no |
| *AvrLm4-7* | *AvrLm4-7-new-del-T-1* | a4a7 | no |
| *AvrLm4-7* | *AvrLm4-7-1-new-A* | a4A7 | yes |
| *AvrLm4-7* | *AvrLm4-7-1-new-B* | a4A7 | yes |
| *AvrLm4-7* | *AvrLm4-7-1-new-C* | a4A7 | yes |
| *AvrLm4-7* | *AvrLm4-7-1-new-D* | a4A7 | yes |
| *AvrLm4-7* | *AvrLm4-7-1-new-E* | a4A7 | yes |
| *AvrLm4-7* | *AvrLm4-7-2-new-A* | a4A7 | yes |
| *AvrLm4-7* | *AvrLm4-7-2-new-B* | a4A7 | yes |
| *AvrLm4-7* | *AvrLm4-7-2-new-C* | a4A7 | yes |
| *AvrLm4-7* | *AvrLm4-7-2-new-D* | a4A7 | yes |
| *AvrLm4-7* | *AvrLm4-7-2-new-E* | a4A7 | yes |
| *AvrLm4-7* | *AvrLm4-7-2-new-F* | a4A7 | yes |
| *AvrLm4-7* | *AvrLm4-7-2-new-G* | a4A7 | yes |
| *AvrLm4-7* | *AvrLm4-7-2-new-H* | a4A7 | yes |
| *AvrLm4-7* | *AvrLm4-7-7-new-A* | a4a7 | yes |
| *AvrLm4-7* | *AvrLm4-7-7-new-B* | a4a7 | yes |
| *AvrLm4-7* | *AvrLm4-7-7-new-C* | a4a7 | yes |
| *AvrLm4-7* | *AvrLm4-7-8-new-A* | a4a7 | yes |
| *AvrLm4-7* | *AvrLm4-7-8-new-B* | a4a7 | yes |
| *AvrLm4-7* | *AvrLm4-7-9-new-A* | a4a7 | yes |
| *AvrLm4-7* | *AvrLm4-7-9-new-B* | a4a7 | yes |
| *AvrLm4-7* | *AvrLm4-7-10-new-A* | a4a7 | yes |

| | | | |
|---|---|---|---|
| *AvrLm4-7* | *AvrLm4-7-18-new-A* | a4a7 | yes |
| *AvrLm4-7* | *AvrLm4-7-18-new-B* | a4a7 | yes |
| *AvrLm4-7* | *AvrLm4-7-18-new-C* | a4a7 | yes |
| *AvrLm4-7* | *AvrLm4-7-18-new-D* | a4a7 | yes |
| *AvrLm4-7* | *AvrLm4-7-29-new* | a4A7 | yes |
| *AvrLm4-7* | *AvrLm4-7-del-AA-1-new* | a4a7 | no |
| *AvrLm4-7* | *AvrLm4-7-rip-new-A* | a4a7 | no |
| *AvrLm4-7* | *AvrLm4-7-rip-new-B* | a4a7 | no |
| *AvrLm4-7* | *AvrLm4-7-rip-new-C* | a4a7 | no |
| *AvrLm4-7* | *AvrLm4-7-rip-new-D* | a4a7 | no |
| *AvrLm4-7* | *AvrLm4-7-rip-new-E* | a4a7 | no |
| *AvrLm4-7* | *AvrLm4-7-rip-new-F* | a4a7 | no |
| *AvrLm4-7* | *AvrLm4-7-rip-new-G* | a4a7 | no |
| *AvrLm4-7* | *AvrLm4-7-rip-new-H* | a4a7 | no |
| *AvrLm4-7* | *AvrLm4-7-rip-new-I* | a4a7 | no |
| *AvrLm4-7* | *AvrLm4-7-rip-new-J* | a4a7 | no |
| *AvrLm4-7* | *AvrLm4-7-rip-new-K-1* | a4a7 | no |
| *AvrLm4-7* | *AvrLm4-7-rip-new-K-2* | a4a7 | no |
| *AvrLm4-7* | *AvrLm4-7-rip-new-L* | a4a7 | no |
| *AvrLm4-7* | *AvrLm4-7-rip-new-M* | a4a7 | no |
| *AvrLm4-7* | *AvrLm4-7-rip-new-N* | a4a7 | no |
| *AvrLm4-7* | *AvrLm4-7-rip-new-O* | a4a7 | no |
| *AvrLm4-7* | *AvrLm4-7-rip-new-P* | a4a7 | no |
| *AvrLm4-7* | *AvrLm4-7-rip-new-Q* | a4a7 | no |
| *AvrLm4-7* | *AvrLm4-7-rip-new-R* | a4a7 | no |
| *AvrLm4-7* | *AvrLm4-7-rip-new-S* | a4a7 | no |
| *AvrLm4-7* | *AvrLm4-7-rip-new-T* | a4a7 | no |
| *AvrLm4-7* | *AvrLm4-7-rip-new-U* | a4a7 | no |
| *AvrLm4-7* | *AvrLm4-7-rip-new-V* | a4a7 | no |
| *AvrLm4-7* | *AvrLm4-7-rip-new-W* | a4a7 | no |

| | | | |
|---|---|---|---|
| *AvrLm4-7* | *AvrLm4-7-rip-new-X* | a4a7 | no |
| *AvrLm4-7* | *AvrLm4-7-rip-new-Y* | a4a7 | no |
| *AvrLm4-7* | *AvrLm4-7-rip-new-Z* | a4a7 | no |
| *AvrLm4-7* | *AvrLm4-7-rip-new-AA* | a4a7 | no |
| *AvrLm4-7* | *AvrLm4-7-rip-new-AB* | a4a7 | no |
| *AvrLm4-7* | *AvrLm4-7-rip-new-AC* | a4a7 | no |
| *AvrLm4-7* | *AvrLm4-7-rip-new-AD* | a4a7 | no |
| *AvrLm4-7* | *AvrLm4-7-rip-new-AE* | a4a7 | no |
| *AvrLm4-7* | *AvrLm4-7-rip-new-AF* | a4a7 | no |
| *AvrLm4-7* | *AvrLm4-7-rip-new-AG* | a4a7 | no |
| *AvrLm4-7* | *AvrLm4-7-rip-new-AH* | a4a7 | no |
| *AvrLm4-7* | *AvrLm4-7-rip-new-AI* | a4a7 | no |
| *AvrLm4-7* | *AvrLm4-7-rip-new-AJ* | a4a7 | no |
| *AvrLm4-7* | *AvrLm4-7-rip-new-AK* | a4a7 | no |
| *AvrLm4-7* | *AvrLm4-7-rip-new-AL* | a4a7 | no |
| *AvrLm4-7* | *AvrLm4-7-rip-new-AM* | a4a7 | no |
| *AvrLm4-7* | *AvrLm4-7-rip-new-AN* | a4a7 | no |
| *AvrLm4-7* | *AvrLm4-7-rip-new-AO* | a4a7 | no |
| *AvrLm4-7* | *AvrLm4-7-rip-new-AP* | a4a7 | no |
| *AvrLm4-7* | *AvrLm4-7-rip-new-AQ* | a4a7 | no |
| *AvrLm4-7* | *AvrLm4-7-rip-new-AR* | a4a7 | no |
| *AvrLm4-7* | *AvrLm4-7-rip-new-AS* | a4a7 | no |
| *AvrLm4-7* | *AvrLm4-7-rip-new-AT* | a4a7 | no |
| *AvrLm4-7* | *AvrLm4-7-rip-new-AU* | a4a7 | no |
| *AvrLm4-7* | *AvrLm4-7-rip-new-AV* | a4a7 | no |
| *AvrLm4-7* | *AvrLm4-7-rip-new-AW* | a4a7 | no |
| *AvrLm4-7* | *AvrLm4-7-rip-new-AX* | a4a7 | no |
| *AvrLm4-7* | *AvrLm4-7-rip-new-AY* | a4a7 | no |
| *AvrLm4-7* | *AvrLm4-7-rip-new-AZ* | a4a7 | no |
| *AvrLm4-7* | *AvrLm4-7-rip-new-BA* | a4a7 | no |

| | | | |
|---|---|---|---|
| *AvrLm4-7* | *AvrLm4-7-rip-new-BB* | a4a7 | no |
| *AvrLm4-7* | *AvrLm4-7-rip-new-BC* | a4a7 | no |
| *AvrLm4-7* | *AvrLm4-7-rip-new-BD* | a4a7 | no |
| *AvrLm4-7* | *AvrLm4-7-rip-new-BE* | a4a7 | no |
| *AvrLm4-7* | *AvrLm4-7-rip-new-BF* | a4a7 | no |
| *AvrLm4-7* | *AvrLm4-7-rip-new-BG* | a4a7 | no |
| *AvrLm4-7* | *AvrLm4-7-rip-new-BH* | a4a7 | no |
| *AvrLm4-7* | *AvrLm4-7-rip-new-BI* | a4a7 | no |
| *AvrLm4-7* | *AvrLm4-7-rip-new-BJ* | a4a7 | no |
| *AvrLm4-7* | *AvrLm4-7-rip-new-BK* | a4a7 | no |
| *AvrLm4-7* | *AvrLm4-7-rip-new-BL* | a4a7 | no |
| *AvrLm4-7* | *AvrLm4-7-rip-new-BM* | a4a7 | no |
| *AvrLm4-7* | *AvrLm4-7-rip-new-BN* | a4a7 | no |
| *AvrLm4-7* | *AvrLm4-7-rip-new-BO* | a4a7 | no |
| *AvrLm4-7* | *AvrLm4-7-rip-new-BP* | a4a7 | no |
| *AvrLm4-7* | *AvrLm4-7-rip-new-BQ* | a4a7 | no |
| *AvrLm4-7* | *AvrLm4-7-rip-new-BR* | a4a7 | no |
| *AvrLm4-7* | *AvrLm4-7-rip-new-BS* | a4a7 | no |
| *AvrLm4-7* | *AvrLm4-7-rip-new-BT* | a4a7 | no |
| *AvrLm4-7* | *AvrLm4-7-rip-new-BU* | a4a7 | no |
| *AvrLm4-7* | *AvrLm4-7-rip-new-BV* | a4a7 | no |
| *AvrLm4-7* | *AvrLm4-7-rip-new-BW* | a4a7 | no |
| *AvrLm4-7* | *AvrLm4-7-rip-new-BX* | a4a7 | no |
| *AvrLm4-7* | *AvrLm4-7-rip-new-BY* | a4a7 | no |
| *AvrLm4-7* | *AvrLm4-7-rip-new-BZ* | a4a7 | no |
| *AvrLm4-7* | *AvrLm4-7-rip-new-CA* | a4a7 | no |
| *AvrLm4-7* | *AvrLm4-7-rip-new-CB* | a4a7 | no |
| *AvrLm4-7* | *AvrLm4-7-rip-new-CC* | a4a7 | no |
| *AvrLm4-7* | *AvrLm4-7-rip-new-CD* | a4a7 | no |
| *AvrLm4-7* | *AvrLm4-7-rip-new-CE* | a4a7 | no |

| | | | |
|---|---|---|---|
| *AvrLm4-7* | *AvrLm4-7-rip-new-CF* | a4a7 | no |
| *AvrLm4-7* | *AvrLm4-7-rip-new-CG* | a4a7 | no |
| *AvrLm4-7* | *AvrLm4-7-rip-new-CH* | a4a7 | no |
| *AvrLm4-7* | *AvrLm4-7-rip-new-CI* | a4a7 | no |
| *AvrLm5-9* | *AvrLm-5-9-0-A* | A5a9 | |
| *AvrLm5-9* | *AvrLm-5-9-0-B* | A5a9 | |
| *AvrLm5-9* | *AvrLm-5-9-1-A* | A5A9 | |
| *AvrLm5-9* | *AvrLm-5-9-6-A* | A5a9 | |
| *AvrLm5-9* | *AvrLm5-9-del* | a5a9 | |
| *AvrLm5-9* | *AvrLm-5-9-0-A-var-ext* | A5a9 | |
| *AvrLm5-9* | *AvrLm5-9-0-new-A* | A5a9 | |
| *AvrLm5-9* | *AvrLm-5-9-0-new-B* | A5a9 | |
| *AvrLm5-9* | *AvrLm-5-9-0-new-C* | A5a9 | |
| *AvrLm5-9* | *AvrLm-5-9-0-new-D* | A5a9 | |
| *AvrLm5-9* | *AvrLm5-9-0-new-E* | A5a9 | |
| *AvrLm5-9* | *AvrLm5-9-0-new-F* | A5a9 | |
| *AvrLm5-9* | *AvrLm5-9-0-new-G* | A5a9 | |
| *AvrLm5-9* | *AvrLm5-9-0-new-H* | A5a9 | |
| *AvrLm5-9* | *AvrLm5-9-0-new-I* | A5a9 | |
| *AvrLm5-9* | *AvrLm5-9-0-new-J* | A5a9 | |
| *AvrLm5-9* | *AvrLm5-9-0-new-K* | A5a9 | |
| *AvrLm5-9* | *AvrLm5-9-0-new-L* | A5a9 | |
| *AvrLm5-9* | *AvrLm-5-9-new-A* | A5a9 | |
| *AvrLm5-9* | *AvrLm-5-9-new-B* | NA | |
| *AvrLm5-9* | *AvrLm-5-9-new-C* | NA | |
| *AvrLm5-9* | *AvrLm5-9-new-D-1* | NA | |
| *AvrLm5-9* | *AvrLm5-9-new-D-2* | NA | |
| *AvrLm5-9* | *AvrLm5-9-new-E* | NA | |
| *AvrLm5-9* | *AvrLm5-9-new-F* | NA | |
| *AvrLm5-9* | *AvrLm5-9-new-G* | NA | |

| | | |
|---|---|---|
| *AvrLm5-9* | *AvrLm5-9-new-H-1* | NA |
| *AvrLm5-9* | *AvrLm5-9-new-H-2* | NA |
| *AvrLm6* | *AvrLm6-0-A-0-B-3-A* | A6 |
| *AvrLm6* | *AvrLm6-1-A* | A6 |
| *AvrLm6* | *AvrLm6-7-A* | a6 |
| *AvrLm6* | *AvrLm6-del* | a6 |
| *AvrLm6* | *AvrLm6-new-A* | NA |
| *AvrLm6* | *AvrLm6-new-C* | NA |
| *AvrLm6* | *AvrLm6-0-new-A* | A6 |
| *AvrLm6* | *AvrLm6-0-new-B* | A6 |
| *AvrLm6* | *AvrLm6-0-new-C* | A6 |
| *AvrLm6* | *AvrLm6-0-new-D* | A6 |
| *AvrLm11* | *AvrLm11-0-A* | A11 |
| *AvrLm11* | *AvrLm11-3-A* | a11 |
| *AvrLm11* | *AvrLm11-del* | a11 |
| *AvrLm11* | *AvrLm11-new-A* | NA |
| *AvrLm11* | *AvrLm11-new-B* | NA |
| *AvrLm11* | *AvrLm11-new-C* | a11 |
| *AvrLm11* | *AvrLm11-new-D* | a11 |
| *AvrLm11* | *AvrLm11-new-E* | NA |
| *AvrLm11* | *AvrLm11-new-F* | NA |
| *AvrLm11* | *AvrLm11-new-G-1* | a11 |
| *AvrLm11* | *AvrLm11-new-G-2* | a11 |
| *AvrLm11* | *AvrLm11-new-H* | NA |
| *AvrLm11* | *AvrLm11-new-I* | NA |
| *AvrLm11* | *AvrLm11-new-J* | NA |
| *AvrLm11* | *AvrLm11-new-K* | A11 |
| *AvrLm11* | *AvrLm11-new-L* | a11 |
| *AvrLm11* | *AvrLm11-new-M* | NA |
| *AvrLm11* | *AvrLm11-new-N* | a11 |

| | | |
|---|---|---|
| *AvrLm11* | *AvrLm11-new-O* | a11 |
| *AvrLm11* | *AvrLm11-new-P* | a11 |
| *AvrLm11* | *AvrLm11-new-Q* | NA |
| *AvrLm11* | *AvrLm11-new-R* | NA |
| *AvrLm11* | *AvrLm11-new-S* | NA |
| *AvrLm11* | *AvrLm11-new-U* | NA |
| *AvrLm11* | *AvrLm11-new-V* | a11 |
| *AvrLm11* | *AvrLm11-0-new-A* | A11 |
| *AvrLm11* | *AvrLm11-0-new-B* | A11 |
| *AvrLm11* | *AvrLm11-0-new-C* | A11 |
| *AvrLm11* | *AvrLm11-0-new-D* | A11 |
| *AvrLm11* | *AvrLm11-0-new-E* | A11 |
| *AvrLm11* | *AvrLm11-0-new-F* | A11 |
| *AvrLm11* | *AvrLm11-0-new-G* | A11 |
| *AvrLm11* | *AvrLm11-0-new-H* | A11 |
| *AvrLm11* | *AvrLm11-0-new-I* | A11 |
| *AvrLm11* | *AvrLm11-0-new-J* | A11 |
| *AvrLm11* | *AvrLm11-0-new-K* | A11 |
| *AvrLm11* | *AvrLm11-0-A-var-ext-01* | A11 |
| *AvrLm11* | *AvrLm11-0-A-var-ext-02* | A11 |
| *AvrLm11* | *AvrLm11-0-A-var-ext-03* | A11 |
| *AvrLm11* | *AvrLm11-0-A-var-ext-04* | A11 |
| *AvrLm11* | *AvrLm11-0-A-var-ext-05* | A11 |
| *AvrLm11* | *AvrLm11-0-A-var-ext-06* | A11 |
| *AvrLm11* | *AvrLm11-0-A-var-ext-07* | A11 |
| *AvrLm11* | *AvrLm11-0-A-var-ext-08* | A11 |
| *AvrLm11* | *AvrLm11-rip-new-A* | a11 |
| *AvrLmS* | *AvrLmS-0-A* | AS |
| *AvrLmS* | *AvrLmS-0-B* | AS |
| *AvrLmS* | *AvrLmS-0-C* | AS |

| | | |
|---|---|---|
| *AvrLmS* | *AvrLmS-1-A* | AS |
| *AvrLmS* | *AvrLmS-1-B* | AS |
| *AvrLmS* | *AvrLmS-2-A* | AS |
| *AvrLmS* | *AvrLmS-2-B* | AS |
| *AvrLmS* | *AvrLmS-2-C* | AS |
| *AvrLmS* | *AvrLmS-2-D* | AS |
| *AvrLmS* | *AvrLmS-2-E* | AS |
| *AvrLmS* | *AvrLmS-2-G* | AS |
| *AvrLmS* | *AvrLmS-2-H* | AS |
| *AvrLmS* | *AvrLmS-2-I* | AS |
| *AvrLmS* | *AvrLmS-2-J* | AS |
| *AvrLmS* | *AvrLmS-2-K* | AS |
| *AvrLmS* | *AvrLmS-3-A* | AS |
| *AvrLmS* | *AvrLmS-4-A* | AS |
| *AvrLmS* | *AvrLmS-4-B* | AS |
| *AvrLmS* | *AvrLmS-6-A* | AS |
| *AvrLmS* | *AvrLmS-6-B* | AS |
| *AvrLmS* | *AvrLmS-7-A* | AS |
| *AvrLmS* | *AvrLmS-7-B* | AS |
| *AvrLmS* | *AvrLmS-9-B* | aS |
| *AvrLmS* | *AvrLmS-21-A* | AS |
| *AvrLmS* | *AvrLmS-21-B* | AS |
| *AvrLmS* | *AvrLmS-new-A-01* | AS |
| *AvrLmS* | *AvrLmS-new-A-02* | AS |
| *AvrLmS* | *AvrLmS-new-B* | AS |
| *AvrLmS* | *AvrLmS-new-C* | AS |
| *AvrLmS* | *AvrLmS-new-E* | NA |
| *AvrLmS* | *AvrLmS-new-F* | AS |
| *AvrLmS* | *AvrLmS-new-G-01* | aS |
| *AvrLmS* | *AvrLmS-new-G-02* | aS |

| | | |
|---|---|---|
| *AvrLmS* | *AvrLmS-new-H* | NA |
| *AvrLmS* | *AvrLmS-new-I* | AS |
| *AvrLmS* | *AvrLmS-0-new-B* | AS |
| *AvrLmS* | *AvrLmS-0-new-D* | AS |
| *AvrLmS* | *AvrLmS-0-new-E* | AS |
| *AvrLmS* | *AvrLmS-0-new-F* | AS |
| *AvrLmS* | *AvrLmS-0-new-G* | AS |
| *AvrLmS* | *AvrLmS-1-new-A* | AS |
| *AvrLmS* | *AvrLmS-1-new-B* | AS |
| *AvrLmS* | *AvrLmS-2-new-B* | AS |
| *AvrLmS* | *AvrLmS-2-new-C* | AS |
| *AvrLmS* | *AvrLmS-2-new-E* | AS |
| *AvrLmS* | *AvrLmS-2-new-F* | AS |
| *AvrLmS* | *AvrLmS-2-new-G* | AS |
| *AvrLmS* | *AvrLmS-2-new-H* | AS |
| *AvrLmS* | *AvrLmS-2-new-I* | AS |
| *AvrLmS* | *AvrLmS-2-new-J* | AS |
| *AvrLmS* | *AvrLmS-2-new-L* | AS |
| *AvrLmS* | *AvrLmS-3-new-A* | AS |
| *AvrLmS* | *AvrLmS-4-new-A* | AS |
| *AvrLmS* | *AvrLmS-4-new-B* | AS |
| *AvrLmS* | *AvrLmS-6-new-B* | AS |
| *AvrLmS* | *AvrLmS-6-new-C* | AS |
| *AvrLmS* | *AvrLmS-6-new-D* | AS |
| *AvrLmS* | *AvrLmS-6-new-E* | AS |
| *AvrLmS* | *AvrLmS-7-new-A* | AS |
| *AvrLmS* | *AvrLmS-7-new-B* | AS |
| *AvrLmS* | *AvrLmS-7-new-C* | AS |
| *AvrLmS* | *AvrLmS-7-new-D* | AS |
| *AvrLmS* | *AvrLmS-del* | aS |

*& A, avirulent; a ; virulent; NA, not assigned*

[a] *Knowing that some AvrLm4-7 alleles are able to mask the AvrLm3/Rlm3 interaction leading to a virulent phenotype on Rlm3 whatever the AvrLm3 allele, the information in this column indicates the masking ability (Yes) or not (no) of each AvrLm4-7 Allele; NK, masking effect not known*

[b] For avirulence genes inducing resistance on potentially to *Rlm* genes (*AvrLm4-7*; *AvrLm5-9*), the two phenotypes are given, with for instance A4A7 for alleles avirulent on R-*Rlm4* and *Rlm7*, or A5a9 for alleles inducing avirulence on *Rlm5* but virulence on *Rlm9*

**Supplementary file S8. percentages of MPSeqM ASV that are not assigned to a corresponding avirulence phenotype for the two sampling seasons 2019-20 and 2022-23.** The tables indicate the percentages per avirulence gene, sampling site and season. Sampling sites ID are described in Supplementary Table S2. The graph represents the mean and standard error values over all sites per avirulence gene and growing season

| Sampling year 2019-2020 | | | | | | | | | | | |
|---|---|---|---|---|---|---|---|---|---|---|---|
| *Avrlm gene* | Sampling site | | | | | | | | | Mean | Standard error |
| | RHE-2019 (35) | ERD (49) | BOI (41) | BLC (36) | JAN (28) | THI-B (78) | THI-T (78) | COU (45) | FRE (59) | | |
| *AvrLm1* | 0,00% | 0,00% | 0,00% | 0,00% | 0,00% | 0,00% | 0,00% | 0,00% | 0,00% | 0,00% | 0,00% |
| *AvrLm2* | 0,00% | 0,00% | 0,00% | 0,00% | 0,00% | 0,00% | 0,00% | 0,00% | 0,00% | 0,00% | 0,00% |
| *AvrLm3* | 0,45% | 0,10% | 0,22% | 0,31% | 0,98% | 0,30% | 1,07% | 1,12% | 0,26% | 0,53% | 0,40% |
| *AvrLm4-7* | 2,04% | 2,58% | 1,58% | 0,56% | 1,61% | 3,59% | 4,56% | 1,11% | 2,32% | 2,22% | 1,24% |
| *AvrLm5-9* | 0,00% | 0,72% | 0,00% | 0,00% | 0,00% | 0,00% | 0,00% | 0,00% | 0,00% | 0,08% | 0,24% |
| *AvrLm6* | 0,00% | 0,00% | 0,45% | 0,00% | 0,00% | 0,00% | 0,00% | 0,00% | 0,00% | 0,05% | 0,15% |
| *AvrLm11* | 1,07% | 0,00% | 0,72% | 0,00% | 0,18% | 0,20% | 0,00% | 1,39% | 1,76% | 0,59% | 0,67% |
| *AvrLmS* | 0,00% | 0,00% | 0,00% | 0,00% | 0,00% | 0,43% | 0,00% | 0,00% | 0,00% | 0,05% | 0,14% |
| Mean | 0,45% | 0,43% | 0,37% | 0,11% | 0,35% | 0,56% | 0,70% | 0,45% | 0,54% | 0,44% | 0,17% |

| Sampling year 2022-2023 | | | | | | | | | | |
|---|---|---|---|---|---|---|---|---|---|---|
| *AvrLm gene* | Sampling site | | | | | | | | Mean | Standard error |
| | RHE-2022 (35) | VIL (41) | TER (28) | TOU (28) | VERS (78) | SPT (62) | ENN (59) | BRE (21) | | |
| *AvrLm1* | 0,00% | 0,00% | 0,00% | 0,00% | 0,00% | 0,00% | 0,00% | 0,00% | 0,00% | 0,00% |
| *AvrLm2* | 0,00% | 0,00% | 0,00% | 0,00% | 0,00% | 0,00% | 0,00% | 0,00% | 0,00% | 0,00% |
| *AvrLm3* | 1,00% | 1,29% | 3,05% | 1,79% | 1,43% | 4,11% | 1,44% | 0,00% | 1,76% | 1,27% |
| *AvrLm4-7* | 2,56% | 5,84% | 6,91% | 3,32% | 4,03% | 4,85% | 2,97% | 0,00% | 3,81% | 2,13% |
| *AvrLm5-9* | 0,39% | 0,00% | 0,00% | 0,00% | 0,00% | 0,00% | 2,35% | 0,00% | 0,34% | 0,82% |
| *AvrLm6* | 0,00% | 0,46% | 0,00% | 0,00% | 0,00% | 0,00% | 0,00% | 0,00% | 0,06% | 0,16% |
| *AvrLm11* | 0,00% | 0,00% | 0,00% | 0,00% | 0,00% | 0,00% | 0,26% | 0,00% | 0,03% | 0,09% |
| *AvrLmS* | 0,00% | 0,00% | 0,00% | 1,19% | 0,00% | 0,00% | 0,00% | 0,00% | 0,15% | 0,42% |
| *Mean* | 0,49% | 0,95% | 1,24% | 0,79% | 0,68% | 1,12% | 0,88% | 0,00% | 0,77% | 0,39% |

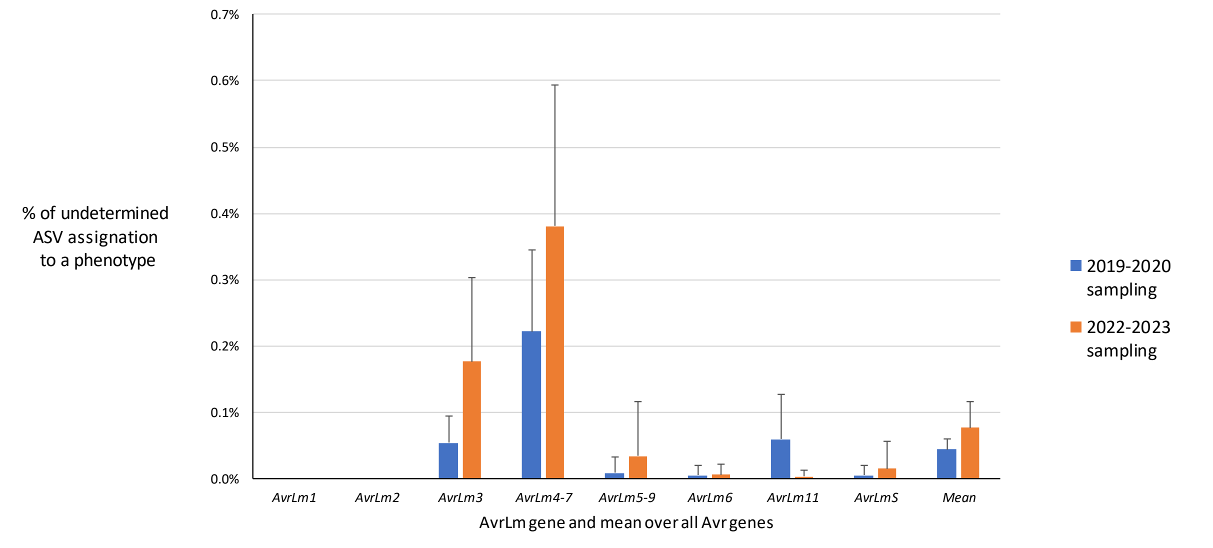

**Supplementary Figure S9. Hierarchical clustering of ASV proportions determined in the 86 environmental samples (pools of *Leptosphaeria maculans* spots on *Brassica napus* leaves) collected in France in 2019-20 and 2022-23 growing seasons.** Each column corresponds to one environmental sample as labelled on the bottom of the graph. Codes and details on all samples are described in Supplementary Table S2. The heatmap shows proportions of detected ASVs after hierarchical clustering based on similarity of sample profiles. Each row corresponds to one avirulence allele (ASV) as indicated on the right. The 50 most represented ASVs are used here. Colors indicate the ASV proportions from low (blue) to high (red).

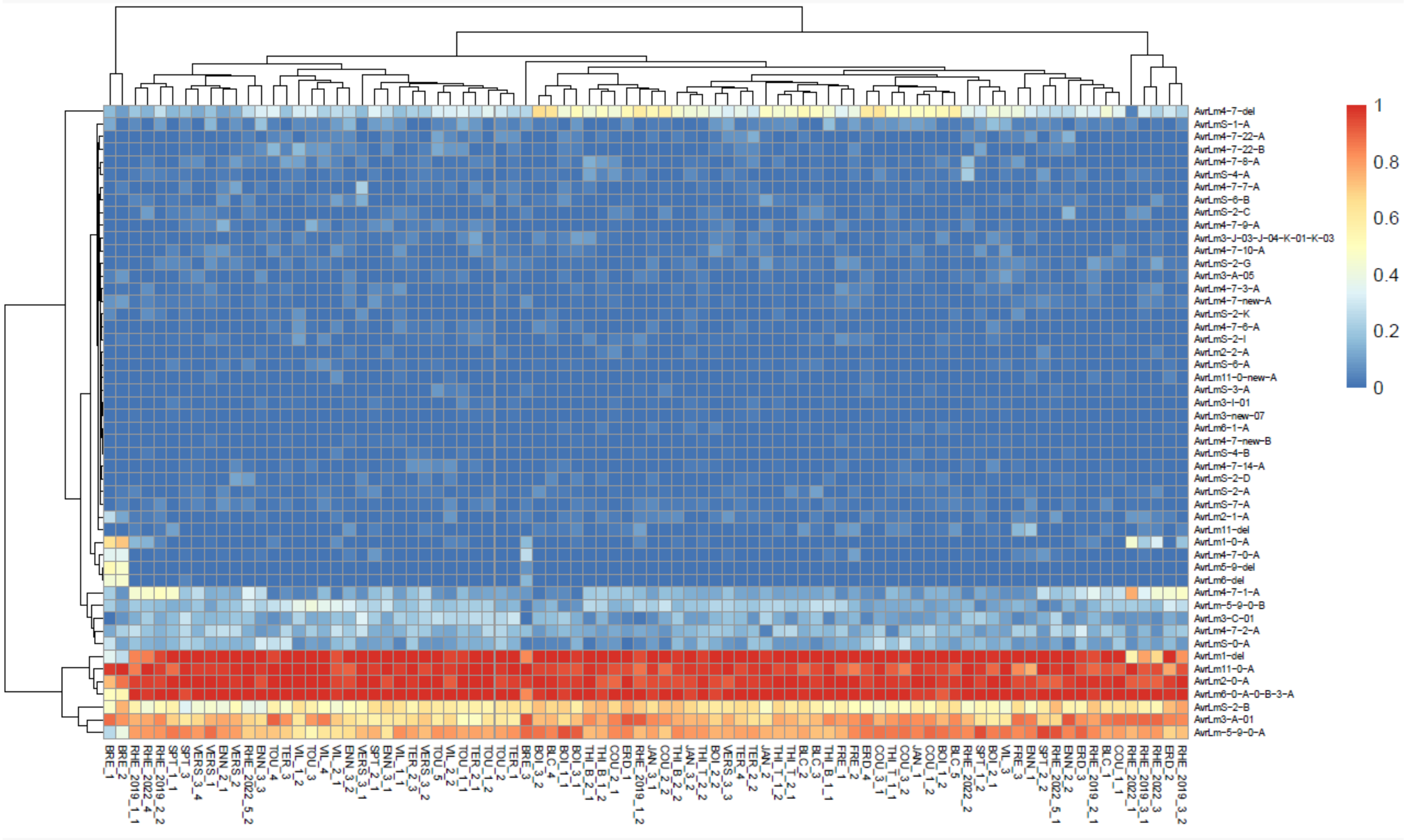

### Supplementary Table S10. Results of phenotyping, PCR amplification and *AvrLm* gene sequencing of selected individual field isolates from the 2022-23 growing season

| Isolate ID | isolated at | Isolated from (Rapeseed genotype) | Phenotype following inoculation | | | PCR amplication results | | | | Genotype identified by Sanger sequencing | | | |
|---|---|---|---|---|---|---|---|---|---|---|---|---|---|
| | | | *Avrlm6* | *AvrLm11* | *AvrLmS* | **AvrLm5-9** | **AvrLm6** | **AvrLm11** | **AvrLmS** | ***AvrLm6*** | ***AvrLm11*** | ***AvrLmS*** | ***AvrLm5-9*** |
| CP.22.800 | Bretenières | Drakkar | V[1] | A | A | -[2] | - | + | nd | *AvrLm6-del*[3] | *AvrLm11-0-A_(893 pb)*[4] | nd | *AvrLm5-9-del* |
| CP.22.801 | Bretenières | Drakkar | V | A | A | - | + | + | nd | *AvrLm6-ripdup-new-A* | *AvrLm11-0-A_(876 pb_1)* | nd | *AvrLm5-9-del* |
| CP.22.802 | Bretenières | Drakkar | A | A | A | + | + | + | nd | *AvrLm6-0-A* | *AvrLm11-0-A_(893 pb)* | nd | *A5-9-0-A* |
| CP.22.803 | Bretenières | Drakkar | V | V | A | + | - | + | nd | *AvrLm6-del* | *AvrLm11-0-A_(876 pb_1)* | nd | *A5-9-0-B* |
| CP.22.804 | Bretenières | Drakkar | V | V | A | - | - | + | nd | *AvrLm6-del* | *AvrLm11-0-A_(876 pb_1)* | nd | *AvrLm5-9-del* |
| CP.22.805 | Bretenières | Drakkar | V | H | A | + | - | + | nd | *AvrLm6-del* | *AvrLm11-0-A_(893 pb)* | nd | *A5-9-new-I* |
| CP.22.806 | Bretenières | Drakkar | V | H | A | - | +/- | + | nd | *AvrLm6-ripdup-new-B* | *AvrLm11-0-A_(842 pb)* | nd | *AvrLm5-9-del* |
| CP.22.808 | Bretenières | Drakkar | A | V | A | + | + | - | nd | *AvrLm6-0-A* | *AvrLm11-del* | nd | *A5-9-0-A* |
| CP.22.809 | Bretenières | Drakkar | V | V | A | - | + | + | nd | *AvrLm6-rip-new-A* | *AvrLm11-0-A_(893 pb)* | nd | *AvrLm5-9-del* |
| CP.22.810 | Bretenières | Drakkar | V | H | A | - | - | + | nd | *AvrLm6-del* | *AvrLm11-0-A_(893 pb)* | nd | *AvrLm5-9-del* |
| CP.22.811 | Bretenières | Drakkar | V | H | A | + | + | + | nd | *AvrLm6-rip-new-B* | *AvrLm11-0-A_(893 pb)* | nd | *AvrLm5-9-rip-01 (new)* |
| CP.22.812 | Bretenières | Drakkar | V | V | A | - | - | + | nd | *AvrLm6-del* | *AvrLm11-0-A_(893 pb)* | nd | *AvrLm5-9-del* |
| CP.22.813 | Bretenières | Drakkar | V | H | A | - | - | + | nd | *AvrLm6-del* | *AvrLm11-0-A_(876 pb_1)* | nd | *AvrLm5-9-del* |
| CP.22.814 | Bretenières | Drakkar | V | V | A | + | - | - | nd | *AvrLm6-del* | *AvrLm11-del* | nd | *A5-9-new-J* |
| CP.22.815 | Bretenières | Drakkar | V | H | A | + | + | + | nd | *AvrLm6-0-A* | *AvrLm11-0-A_(876 pb_1)* | nd | *A5-9-new-J* |
| CP.22.816 | Bretenières | Drakkar | V | V | A | - | - | + | nd | *AvrLm6-del* | *AvrLm11-0-A_(875 pb)* | nd | *AvrLm5-9-del* |
| CP.22.817 | Bretenières | Drakkar | V | V | A | - | - | + | nd | *AvrLm6-del* | *AvrLm11-0-A_(950 pb)* | nd | *AvrLm5-9-del* |
| CP.22.818 | Bretenières | Drakkar | V | A | A | - | + | + | nd | *AvrLm6-ripdup-L* | *AvrLm11-0-A_(876 pb_1)* | nd | *AvrLm5-9-del* |
| CP.22.819 | Bretenières | Drakkar | V | H | A | - | - | + | nd | *AvrLm6-del* | *AvrLm11-0-A_(893 pb)* | nd | *AvrLm5-9-del* |
| CP.22.820 | Bretenières | Drakkar | V | A | A | - | - | + | nd | *AvrLm6-del* | *AvrLm11-0-A_(893 pb)* | nd | *AvrLm5-9-del* |
| CP.22.821 | Bretenières | Drakkar | A | A | A | + | + | + | nd | *AvrLm6-0-A* | *AvrLm11-0-A_(876 pb_1)* | nd | *A5-9-0-A* |
| CP.22.822 | Bretenières | Drakkar | V | A | A | + | +/- | + | + | *AvrLm6-del* | *AvrLm11-0-A_(893 pb)* | *AvrLmS-0-A* | *A5-9-0-A* |

| | | | | | | | | | | | | | |
|---|---|---|---|---|---|---|---|---|---|---|---|---|---|
| CP.22.823 | Bretenières | Drakkar | V | A | A | - | - | + | nd | *AvrLm6-del* | *AvrLm11-0-A_(893 pb)* | nd | *AvrLm5-9-del* |
| CP.22.824 | Bretenières | Drakkar | V | V | A | - | - | - | nd | *AvrLm6-del* | *AvrLm11-del* | nd | *AvrLm5-9-del* |
| CP.22.826 | Bretenières | Drakkar | V | A | A | - | - | + | nd | *AvrLm6-del* | *AvrLm11-0-A_(893 pb)* | nd | *AvrLm5-9-del* |
| CP.22.827 | Bretenières | Drakkar | V | V | A | + | + | + | nd | *AvrLm6-ripdup-new-C* | *AvrLm11-0-A_(876 pb_1)* | nd | *A5-9-0-C* |
| CP.22.828 | Bretenières | Drakkar | V | H | A | - | - | + | nd | *AvrLm6-del* | *AvrLm11-0-A_(876 pb_1)* | nd | *AvrLm5-9-del* |
| CP.22.829 | Bretenières | Drakkar | A | A | A | + | + | + | nd | *AvrLm6-0-A* | *AvrLm11-0-A_(859 pb)* | nd | *A5-9-0-B* |
| CP.22.830 | Bretenières | Drakkar | V | H | A | + | - | + | nd | *AvrLm6-del* | *AvrLm11-0-A_(876 pb_1)* | nd | *AvrLm5-9-ripdup-A* |
| CP.22.831 | Bretenières | Drakkar | V | V | A | - | - | - | + | *AvrLm6-del* | *AvrLm11-del* | *AvrLmS-2-B* | *AvrLm5-9-del* |
| CP.22.832 | Bretenières | Drakkar | V | H | A | - | + | + | nd | *AvrLm-6-rip-new-C* | *AvrLm11-0-A_(876 pb_1)* | nd | *AvrLm5-9-del* |
| CP.22.833 | Bretenières | Drakkar | V | V | A | + | +/- | + | + | *AvrLm6-del* | *AvrLm11-0-A_(876 pb_2)* | *AvrLmS-2-B* | *A5-9-0-B* |
| CP.22.835 | Bretenières | Drakkar | A | A | A | + | + | + | nd | *AvrLm6-0-A* | *AvrLm11-0-A_(876 pb_1)* | nd | *A5-9-0-B* |
| CP.22.836 | Bretenières | Drakkar | V | H | A | + | + | + | nd | *AvrLm6-7-A* | *AvrLm11-0-A_(910 pb)* | nd | *A5-9-new-J* |
| CP.22.837 | Bretenières | Drakkar | V | V | A | - | +/- | + | nd | *AvrLm6-ripdup-new-D* | *AvrLm11-0-A_(876 pb_1)* | nd | *AvrLm5-9-del* |
| CP.22.838 | Bretenières | Drakkar | V | A | A | - | - | + | nd | *AvrLm6-del* | *AvrLm11-0-A_(876 pb_1)* | nd | *AvrLm5-9-del* |
| CP.22.839 | Bretenières | Drakkar | V | H | A | - | - | + | nd | *AvrLm6-del* | *AvrLm11-0-A_(893 pb)* | nd | *AvrLm5-9-del* |
| CP.22.840 | Bretenières | Drakkar | V | H | A | - | + | + | nd | *AvrLm-6-rip-new-D* | *AvrLm11-0-A_(859 pb)* | nd | *AvrLm5-9-del* |
| CP.22.842 | Bretenières | Drakkar | V | H | A | + | + | + | nd | *AvrLm6-ripdup-new-E* | *AvrLm11-0-A_(893 pb)* | nd | *A5-9-0-B* |
| CP.22.843 | Bretenières | Drakkar | V | V | A | + | + | + | nd | *AvrLm-6-rip-new-E* | *AvrLm11-0-A_(876 pb_1)* | nd | *A5-9-0-B* |
| CP.22.844 | Bretenières | Drakkar | V | A | A | - | - | + | nd | *AvrLm6-del* | *AvrLm11-0-A_(876 pb_1)* | nd | *AvrLm5-9-del* |
| CP.22.845 | Bretenières | Drakkar | V | A | A | - | - | + | nd | *AvrLm6-del* | *AvrLm11-0-A_(893 pb)* | nd | *AvrLm5-9-del* |
| CP.22.846 | Bretenières | Drakkar | V | H | A | - | - | + | nd | *AvrLm6-del* | *AvrLm11-0-A_(876 pb_1)* | nd | *AvrLm5-9-del* |
| CP.22.847 | Bretenières | Marcopolo | A | A | A | + | + | + | nd | *AvrLm6-0-A + AvrLm6-0-C* | *AvrLm11-0-A_(876 pb_1)* | nd | *A5-9-0-B* |
| CP.22.848 | Bretenières | Marcopolo | A | A | A | + | + | + | nd | *AvrLm6-0-A* | *AvrLm11-0-A_(876 pb_1)* | nd | *A5-9-0-A* |
| CP.22.850 | Bretenières | Marcopolo | A | A | A | + | + | + | nd | *AvrLm6-0-A* | *AvrLm11-0-A_(876 pb_1)* | nd | *A5-9-0-B* |
| CP.22.851 | Bretenières | Marcopolo | V | A | A | + | + | + | nd | *AvrLm-6-rip-new-F* | *AvrLm11-0-A_(893 pb)* | nd | *A5-9-0-B* |
| CP.22.852 | Bretenières | Marcopolo | A | A | A | + | + | + | nd | *AvrLm6-0-A* | *AvrLm11-0-A_(876 pb_1)* | nd | *A5-9-0-A* |
| CP.22.853 | Bretenières | Marcopolo | A | A | A | + | + | + | nd | *AvrLm6-0-A* | *AvrLm11-0-A_(893 pb)* | nd | *A5-9-0-A* |
| CP.22.854 | Bretenières | Marcopolo | A | A | V | + | + | + | nd | *AvrLm6-0-A* | *AvrLm11-0-A_(876 pb_1)* | nd | *A5-9-0-A* |
| CP.22.855 | Bretenières | Marcopolo | A | V | A | + | + | - | nd | *AvrLm6-0-A* | *AvrLm11-del* | nd | *A5-9-0-A* |
| CP.22.856 | Bretenières | Marcopolo | A | A | A | + | + | + | nd | *AvrLm6-0-A* | *AvrLm11-0-A_(893 pb)* | nd | *A5-9-0-A* |

| | | | | | | | | | | | | | |
|---|---|---|---|---|---|---|---|---|---|---|---|---|---|
| CP.22.857 | Bretenières | Marcopolo | A | A | A | + | + | + | nd | *AvrLm6-0-A* | *AvrLm11-0-A_(876 pb_1)* | nd | *A5-9-0-A* |
| CP.22.858 | Bretenières | Marcopolo | V | V | A | + | +/- | + | nd | Probable RIP[4] | *AvrLm11-0-A_(893 pb)* | nd | *AvrLm5-9-ripdup-B* |
| CP.22.859 | Bretenières | Marcopolo | A | A | A | + | + | + | nd | *AvrLm6-0-A* | *AvrLm11-0-A_(893 pb)* | nd | *A5-9-0-B* |
| CP.22.860 | Bretenières | Marcopolo | V | A | A | - | +/- | + | nd | *AvrLm6-ripdup-new-F* | *AvrLm11-0-A_(893 pb)* | nd | *AvrLm5-9-del* |
| CP.22.861 | Bretenières | Marcopolo | A | A | A | + | + | + | nd | *AvrLm6-0-A* | *AvrLm11-0-A_(893 pb)* | nd | *A5-9-0-C* |
| CP.22.862 | Bretenières | Marcopolo | V | H | A | - | - | + | nd | *AvrLm6-del* | *AvrLm11-0-A_(876 pb_1)* | nd | *AvrLm5-9-del* |
| CP.22.864 | Bretenières | Marcopolo | A | A | A | + | + | + | nd | *AvrLm6-0-A* | *AvrLm11-0-A_(876 pb_1)* | nd | *A5-9-0-B* |
| CP.22.865 | Bretenières | Marcopolo | A | A | A | + | + | + | nd | *AvrLm6-0-A + AvrLm6-0-D* | *AvrLm11-0-A_(893 pb)* | nd | *A5-9-0-A* |
| CP.22.866 | Bretenières | Marcopolo | A | A | A | + | + | + | nd | *AvrLm6-3-A* | *AvrLm11-0-A_(893 pb)* | nd | *A5-9-0-A* |
| CP.22.867 | Bretenières | Marcopolo | V | A | A | + | +/- | + | nd | *AvrLm6-rip-new-G* | *AvrLm11-0-A_(876 pb_1)* | nd | *A5-9-0-A* |
| CP.22.868 | Bretenières | Marcopolo | V | A | A | - | +/- | + | nd | *AvrLm6-ripdup-new-G* | *AvrLm11-0-A_(876 pb_1)* | nd | *AvrLm5-9-del* |
| CP.22.869 | Bretenières | Marcopolo | A | A | A | + | + | + | nd | *AvrLm6-0-A* | *AvrLm11-new-AA_(910 pb)* | nd | *A5-9-0-A* |
| CP.22.870 | Bretenières | Marcopolo | A | A | A | + | + | + | nd | *AvrLm6-0-A* | *AvrLm11-0-A_(876 pb_1)* | nd | *A5-9-0-B* |
| CP.22.871 | Bretenières | Marcopolo | A | A | A | + | + | + | nd | *AvrLm6-0-A* | *AvrLm11-0-A_(876 pb_1)* | nd | *A5-9-0-A* |
| CP.22.872 | Bretenières | Marcopolo | V | H | A | + | + | + | nd | *AvrLm6-rip-new-H* | *AvrLm11-0-A_(876 pb_1)* | nd | *A5-9-0-B* |
| CP.22.873 | Bretenières | Marcopolo | V | A | A | - | + | + | nd | *AvrLm6-rip-new-I* | *AvrLm11-0-A_(893 pb)* | nd | *AvrLm5-9-del* |
| CP.22.874 | Bretenières | Marcopolo | A | A | A | + | + | + | nd | *AvrLm6-0-A* | *AvrLm11-0-A_(842 pb)* | nd | *A5-9-0-A* |
| CP.22.875 | Bretenières | Marcopolo | A | A | A | + | + | + | nd | *AvrLm6-0-A+AvrLm6-8-A* | *AvrLm11-0-A_(876 pb_1)* | nd | *A5-9-0-B* |
| CP.22.876 | Bretenières | Marcopolo | V | H | A | - | +/- | + | nd | Probable RIP[4] | *AvrLm11-0-A_(893 pb)* | nd | *AvrLm5-9-del* |
| CP.22.877 | Bretenières | Marcopolo | A | A | A | + | + | + | nd | *AvrLm6-0-A* | *AvrLm11-0-A_(910 pb)* | nd | *A5-9-0-A* |
| CP.22.878 | Bretenières | Marcopolo | A | H | A | + | + | + | nd | *AvrLm6-0-A* | *AvrLm11-0-A_(876 pb_1)* | nd | *A5-9-0-C* |
| CP.22.886 | Bretenières | Feliciano *(RlmS)* | A | A | A | + | + | + | + | *AvrLm6-0-A* | *AvrLm11-0-A_(893 pb)* | *AvrLmS-0-A* | *A5-9-0-D (new)* |
| CP.22.887 | Bretenières | Darmor-*Rlm6* | V | A | - | nd | - | + | nd | *AvrLm6-del* | *AvrLm11-0-A_(893 pb)* | nd | nd |
| CP.22.888 | Bretenières | Darmor-*Rlm6* | V | A | - | nd | +/- | + | nd | Probable RIP | *AvrLm11-0-A_(876 pb_1)* | nd | nd |
| CP.22.889 | Bretenières | Darmor-*Rlm6* | V | A | - | nd | - | + | nd | *AvrLm6-del* | *AvrLm11-0-A_(893 pb)* | nd | nd |
| CP.22.890 | Bretenières | Darmor-*Rlm6* | V | A | - | nd | - | + | nd | *AvrLm6-del* | *AvrLm11-0-A_(876 pb_1)* | nd | nd |
| CP.22.891 | Bretenières | Darmor-*Rlm6* | V | A | - | nd | + | + | nd | *AvrLm6-ripdup-new-H* | *AvrLm11-0-A_(893 pb)* | nd | nd |
| CP.22.892 | Bretenières | Darmor-*Rlm6* | V | H | - | nd | - | + | nd | *AvrLm6-del* | *AvrLm11-0-A_(893 pb)* | nd | nd |
| CP.22.893 | Bretenières | Darmor-*Rlm6* | V | V | - | nd | +/- | - | nd | *AvrLm6-del* | *AvrLm11-del* | nd | nd |

| | | | | | | | | | | | | | |
|---|---|---|---|---|---|---|---|---|---|---|---|---|---|
| CP.22.894 | Bretenières | Darmor-*Rlm6* | V | A | - | nd | - | + | nd | *AvrLm6-del* | *AvrLm11-0-A_(876 pb_1)* | nd | nd |
| CP.22.895 | Bretenières | Darmor-*Rlm6* | V | V | - | nd | + | - | nd | *AvrLm6-ripdup-new-I* | *AvrLm11-del* | nd | nd |
| CP.22.896 | Bretenières | Darmor-*Rlm6* | V | A | - | nd | + | + | nd | *AvrLm6-9-A* | *AvrLm11-0-A_(876 pb_1)* | nd | nd |
| CP.22.897 | Bretenières | Darmor-*Rlm6* | V | A | - | nd | + | + | nd | *AvrLm6-0-A* | *AvrLm11-0-A_(876 pb_1)* | nd | nd |
| CP.22.898 | Bretenières | Darmor-*Rlm6* | V | A | - | nd | + | + | nd | *AvrLm6-rip-new-J* | *AvrLm11-0-A_(876 pb_1)* | nd | nd |
| CP.22.899 | Bretenières | Darmor-*Rlm6* | V | A | - | nd | - | + | nd | *AvrLm6-del* | *AvrLm11-0-A_(876 pb_1)* | nd | nd |
| CP.22.900 | Bretenières | Darmor-*Rlm6* | V | H | A | nd | +/- | + | + | *AvrLm6-del* | *AvrLm11-0-A_(893 pb)* | *AvrLmS-1-A (new)* | nd |
| CP.23.114 3 | Aucamville | Darmor-*Rlm11* | A | V | A | nd | + | - | nd | *AvrLm6-0-A* | *AvrLm11-del* | nd | nd |
| CP.22.122 | Versailles | Drakkar | A | V | A | nd | nd | - | nd | nd | *AvrLm11-del* | nd | nd |
| CP.22.177 | Versailles | Marcopolo | A | V | A | nd | nd | - | nd | nd | *AvrLm11-del* | nd | nd |
| CP.22.244 | Terminiers | Marcopolo | A | V | V | nd | + | - | + | *AvrLm6-0-A* | *AvrLm11-del* | *AvrLmS-1-C (new)* | nd |
| CP.22.258 | Terminiers | Marcopolo | A | V | A | nd | nd | - | nd | nd | *AvrLm11-del* | nd | nd |
| CP.22.304 | Villeneuve-Frouville | Drakkar | A | V | A | nd | nd | - | nd | nd | *AvrLm11-del* | nd | nd |
| CP.22.308 | Villeneuve-Frouville | Drakkar | A | V | A | nd | nd | - | nd | nd | *AvrLm11-del* | nd | nd |
| CP.22.325 | Villeneuve-Frouville | Drakkar | A | V | A | nd | nd | + | nd | nd | *AvrLm11-new-Z_(876pb)* | nd | nd |
| CP.22.345 | Villeneuve-Frouville | Drakkar | A | V | A | nd | nd | - | nd | nd | *AvrLm11-del* | nd | nd |
| CP.22.358 | Villeneuve-Frouville | Drakkar | A | A | A | nd | nd | + | nd | nd | *AvrLm11-0-A_(893 pb)* | nd | nd |
| CP.22.371 | Villeneuve-Frouville | Drakkar | A | V | A | nd | nd | - | nd | nd | *AvrLm11-del* | nd | nd |
| CP.22.379 | Villeneuve-Frouville | Drakkar | A | V | A | nd | nd | + | nd | nd | *AvrLm11-rip-new-B_(876pb)* | nd | nd |
| CP.22.461 | St-Pol sur Ternoise | Marcopolo | A | V | A | nd | nd | - | nd | nd | *AvrLm11-del* | nd | nd |
| CP.22.465 | St-Pol sur Ternoise | Marcopolo | A | V | A | nd | nd | - | nd | nd | *AvrLm11-del* | nd | nd |
| CP.22.473 | St-Pol sur Ternoise | Marcopolo | A | V | A | nd | nd | - | nd | nd | *AvrLm11-del* | nd | nd |
| CP.22.506 | Le Rheu | Drakkar | A | V | A | nd | nd | - | nd | nd | *AvrLm11-del* | nd | nd |
| CP.22.508 | Le Rheu | Drakkar | A | V | A | nd | nd | - | nd | nd | *AvrLm11-del* | nd | nd |
| CP.22.533 | Le Rheu | Drakkar | A | V | A | nd | nd | - | nd | nd | *AvrLm11-del* | nd | nd |
| CP.22.540 | Le Rheu | Drakkar | A | V | A | nd | nd | - | nd | nd | *AvrLm11-del* | nd | nd |
| CP.22.573 | Le Rheu | Marcopolo | A | V | A | nd | nd | - | nd | nd | *AvrLm11-del* | nd | nd |
| CP.22.602 | Toury | Drakkar | A | V | A | nd | nd | - | nd | nd | *AvrLm11-del* | nd | nd |

| | | | | | | | | | | | | | |
|---|---|---|---|---|---|---|---|---|---|---|---|---|---|
| CP.22.604 | Toury | Drakkar | A | V | A | nd | nd | - | nd | nd | *AvrLm11-del* | nd | nd |
| CP.22.634 | Toury | Drakkar | A | V | A | nd | nd | + | nd | nd | *AvrLm11-rip-new-C_(893 pb)* | nd | nd |
| CP.22.701 | Ennevin | Drakkar | A | V | A | nd | nd | + | nd | nd | *AvrLm11-rip-new-D_(876 pb)* | nd | nd |
| CP.22.707 | Ennevin | Drakkar | A | V | A | nd | nd | - | nd | nd | *AvrLm11-del* | nd | nd |
| CP.22.1001 | Allenes les Marais | Darmor | A | V | A | nd | + | - | nd | *AvrLm6-0-A* | *AvrLm11-del* | nd | nd |
| CP.23.1135 | Aucamville | Drakkar | A | V | V | nd | + | - | - | *AvrLm6-0-A* | *AvrLm11-del* | *AvrLmS-del* | nd |
| CP.22.568 | Le Rheu | Marcopolo | A | A | V | nd | + | + | + | *AvrLm6-0-A* | *AvrLm11-0-A_(893 pb)* | *AvrLmS-16-B (new)* | nd |
| CP.23.1205 | Verneuil L'Etang | Drakkar | A | A | A | nd | + | + | + | *AvrLm6-0-A* | *AvrLm11-0-A_(876 pb_1)* | *AvrLmS-21-A (new)* | nd |
| CP.23.1213 | Verneuil L'Etang | Drakkar | A | A | A | nd | + | + | + | *AvrLm6-0-A* | *AvrLm11-0-A_(893 pb)* | *AvrLmS-6-A* | nd |
| CP.22.901 | Bretenières | Darmor-*Rlm6* | V | A | - | nd | - | +/- | nd | *AvrLm6-del* | *AvrLm11-0-A_(830 pb)* | nd | nd |
| CP.22.902 | Bretenières | Darmor-*Rlm6* | V | A | - | nd | - | + | nd | *AvrLm6-del* | *AvrLm11-0-A_(876 pb_1)* | nd | nd |
| CP.22.903 | Bretenières | Darmor-*Rlm6* | V | A | - | nd | - | + | nd | *AvrLm6-del* | *AvrLm11-0-A_(876 pb_1)* | nd | nd |
| CP.22.906 | Bretenières | Darmor-*Rlm6* | V | V | - | nd | - | - | nd | *AvrLm6-del* | *AvrLm11-del* | nd | nd |
| CP.22.907 | Bretenières | Darmor-*Rlm6* | V | H | - | nd | - | + | nd | *AvrLm6-del* | *AvrLm11-0-A_(893 pb)* | nd | nd |
| CP.22.908 | Bretenières | Darmor-*Rlm6* | V | A | - | nd | +/- | + | nd | Probable RIP | *AvrLm11-0-A_(876 pb_1)* | nd | nd |
| CP.22.909 | Bretenières | Darmor-*Rlm6* | V | A | - | nd | +/- | + | nd | *AvrLm6-ripdup-new-J* | *AvrLm11-0-A_(876 pb_1)* | nd | nd |
| CP.22.910 | Bretenières | Darmor-*Rlm6* | V | A | - | nd | - | + | nd | *AvrLm6-del* | *AvrLm11-0-A_(876 pb_1)* | nd | nd |
| CP.22.911 | Bretenières | Darmor-*Rlm6* | V | H | - | nd | +/- | + | nd | Probable RIP | *AvrLm11-0-A_(893 pb)* | nd | nd |
| CP.22.912 | Bretenières | Darmor-*Rlm6* | V | H | - | nd | - | + | nd | *AvrLm6-del* | *AvrLm11-0-A_(876 pb_3)* | nd | nd |
| CP.22.913 | Bretenières | Darmor-*Rlm6* | V | A | - | nd | +/- | + | nd | Probable RIP | *AvrLm11-0-A_(893 pb)* | nd | nd |
| CP.22.914 | Bretenières | Darmor-*Rlm6* | V | A | - | nd | + | + | nd | *AvrLm6-ripdup-new-K* | *AvrLm11-0-A_(876 pb_1)* | nd | nd |
| CP.22.915 | Bretenières | Darmor-*Rlm6* | V | A | - | nd | + | + | nd | *AvrLm6-ripdup-new-L* | *AvrLm11-0-A_(876 pb_1)* | nd | nd |
| CP.22.916 | Bretenières | Darmor-*Rlm6* | V | A | - | nd | - | + | nd | *AvrLm6-del* | *AvrLm11-0-A_(876 pb_1)* | nd | nd |
| CP.22.917 | Bretenières | Darmor-*Rlm6* | V | A | - | nd | - | + | nd | *AvrLm6-del* | *AvrLm11-0-new-L_(893 pb)* | nd | nd |
| CP.22.918 | Bretenières | Darmor-*Rlm11* | V | V | - | nd | - | - | nd | *AvrLm6-del* | *AvrLm11-del* | nd | nd |
| CP.22.919 | Bretenières | Darmor-*Rlm11* | A | V | - | nd | + | - | nd | *AvrLm6-0-A* | *AvrLm11-del* | nd | nd |
| CP.22.920 | Bretenières | Darmor-*Rlm11* | V | V | - | nd | nd | - | nd | nd | *AvrLm11-del* | nd | nd |
| CP.22.921 | Bretenières | Darmor-*Rlm11* | V | V | - | nd | nd | - | nd | nd | *AvrLm11-del* | nd | nd |
| CP.22.922 | Bretenières | Darmor-*Rlm11* | V | V | - | nd | nd | - | nd | nd | *AvrLm11-del* | nd | nd |

| | | | | | | | | | | | | | |
|---|---|---|---|---|---|---|---|---|---|---|---|---|---|
| CP.22.923 | Bretenières | Darmor-*Rlm11* | V | V | - | nd | nd | - | nd | nd | *AvrLm11-del* | nd | nd |
| CP.22.924 | Bretenières | Darmor-*Rlm11* | V | V | - | nd | nd | - | nd | nd | *AvrLm11-del* | nd | nd |
| CP.22.925 | Bretenières | Darmor-*Rlm11* | V | V | - | nd | nd | - | nd | nd | *AvrLm11-del* | nd | nd |
| CP.22.926 | Bretenières | Darmor-*Rlm11* | V | V | - | nd | nd | - | nd | nd | *AvrLm11-del* | nd | nd |
| CP.22.927 | Bretenières | Darmor-*Rlm11* | V | V | - | nd | nd | - | nd | nd | *AvrLm11-del* | nd | nd |
| CP.22.928 | Bretenières | Darmor-*Rlm11* | V | V | - | nd | nd | - | nd | nd | *AvrLm11-del* | nd | nd |
| CP.22.929 | Bretenières | Darmor-*Rlm11* | V | A | - | nd | nd | + | nd | nd | *AvrLm11-0-A_(893 pb)* | nd | nd |
| CP.22.930 | Bretenières | Darmor-*Rlm11* | V | V | - | nd | nd | - | nd | nd | *AvrLm11-del* | nd | nd |
| CP.22.931 | Bretenières | Darmor-*Rlm11* | V | V | - | nd | nd | - | nd | nd | *AvrLm11-del* | nd | nd |
| CP.22.932 | Bretenières | Darmor-*Rlm11* | V | V | - | nd | nd | - | nd | nd | *AvrLm11-del* | nd | nd |
| CP.22.933 | Bretenières | Darmor-*Rlm11* | V | V | - | nd | nd | - | nd | nd | *AvrLm11-del* | nd | nd |
| CP.22.934 | Bretenières | Darmor-*Rlm11* | V | V | - | nd | nd | - | nd | nd | *AvrLm11-del* | nd | nd |
| CP.22.935 | Bretenières | Darmor-*Rlm11* | V | V | - | nd | nd | - | nd | nd | *AvrLm11-del* | nd | nd |
| CP.22.936 | Bretenières | Darmor-*Rlm11* | V | V | - | nd | nd | - | nd | nd | *AvrLm11-del* | nd | nd |
| CP.22.937 | Bretenières | Darmor-*Rlm11* | V | V | - | nd | nd | - | nd | nd | *AvrLm11-del* | nd | nd |
| CP.22.938 | Bretenières | Darmor-*Rlm11* | V | V | - | nd | nd | - | nd | nd | *AvrLm11-del* | nd | nd |
| CP.22.939 | Bretenières | Darmor-*Rlm11* | V | V | - | nd | nd | + | nd | nd | *duplication AvrLm11-0-A_(893 pb) & AvrLm11-new-Y_(961 pb)=AvrLm11-7-A* | nd | nd |
| CP.22.382 | Villeneuve-Frouville | Darmor-*Rlm11* | A | V | A | nd | nd | - | nd | nd | *AvrLm11-del* | nd | nd |
| CP.22.383 | Villeneuve-Frouville | Darmor-*Rlm11* | A | V | A | nd | nd | - | nd | nd | *AvrLm11-del* | nd | nd |
| CP.22.384 | Villeneuve-Frouville | Darmor-*Rlm11* | A | V | A | nd | nd | - | nd | nd | *AvrLm11-del* | nd | nd |
| CP.22.385 | Villeneuve-Frouville | Darmor-*Rlm11* | A | V | A | nd | nd | - | nd | nd | *AvrLm11-del* | nd | nd |
| CP.22.386 | Villeneuve-Frouville | Darmor-*Rlm11* | A | V | A | nd | nd | - | nd | nd | *AvrLm11-del* | nd | nd |
| CP.22.387 | Villeneuve-Frouville | Darmor-*Rlm11* | A | V | A | nd | nd | - | nd | nd | *AvrLm11-del* | nd | nd |
| CP.22.388 | Villeneuve-Frouville | Darmor-*Rlm11* | A | V | A | nd | nd | - | nd | nd | *AvrLm11-del* | nd | nd |
| CP.22.389 | Villeneuve-Frouville | Darmor-*Rlm11* | A | V | A | nd | nd | - | nd | nd | *AvrLm11-del* | nd | nd |
| CP.22.390 | Villeneuve-Frouville | Darmor-*Rlm11* | A | V | A | nd | nd | - | nd | nd | *AvrLm11-del* | nd | nd |
| CP.22.391 | Villeneuve-Frouville | Darmor-*Rlm11* | A | V | A | nd | nd | - | nd | nd | *AvrLm11-del* | nd | nd |

| | | | | | | | | | | | | | |
|---|---|---|---|---|---|---|---|---|---|---|---|---|---|
| CP.22.392 | Villeneuve-Frouville | Darmor-*Rlm11* | A | V | V | nd | nd | - | nd | nd | *AvrLm11-del* | nd | nd |
| CP.22.393 | Villeneuve-Frouville | Napoli *(RlmS)* | A | A | V | nd | nd | **+** | **+** | nd | *AvrLm11-0-A_(876 pb_1)* | *AvrLmS-9-A* | nd |
| CP.22.394 | Villeneuve-Frouville | Napoli *(RlmS)* | A | A | V | nd | nd | **+** | **+** | nd | *AvrLm11-0-A_(876 pb_1)* | *AvrLmS-9-A* | nd |
| CP.22.485 | St-Pol sur Ternoise | Darmor-*Rlm11* | A | V | A | nd | nd | - | nd | nd | *AvrLm11-del* | nd | nd |
| CP.22.486 | St-Pol sur Ternoise | Darmor-*Rlm11* | A | V | A | nd | nd | - | nd | nd | *AvrLm11-del* | nd | nd |
| CP.22.487 | St-Pol sur Ternoise | Darmor-*Rlm11* | A | A | A | nd | nd | **+** | nd | nd | *AvrLm11-0-A_(876 pb_1)* | nd | nd |
| CP.22.488 | St-Pol sur Ternoise | Darmor-*Rlm11* | A | V | A | nd | nd | - | nd | nd | *AvrLm11-del* | nd | nd |
| CP.22.489 | St-Pol sur Ternoise | Darmor-*Rlm11* | A | A | A | nd | nd | **+** | nd | nd | *AvrLm11-0-A_(876 pb_1)* | nd | nd |
| CP.22.683 | Toury | Darmor-*Rlm11* | A | A | - | nd | nd | **+** | nd | nd | *AvrLm11-0-A_(893 pb)* | nd | nd |
| CP.22.684 | Toury | Darmor-*Rlm11* | A | V | - | nd | nd | - | nd | nd | *AvrLm11-del* | nd | nd |
| CP.22.685 | Toury | Darmor-*Rlm11* | A | V | - | nd | **+** | - | nd | *AvrLm6-0-A* | *AvrLm11-del* | nd | nd |
| CP.22.686 | Toury | Darmor-*Rlm11* | A | V | - | nd | **+** | - | nd | *AvrLm6-0-A* | *AvrLm11-del* | nd | nd |
| CP.22.687 | Toury | Darmor-*Rlm11* | A | V | - | nd | **+** | - | nd | *AvrLm6-0-A* | *AvrLm11-del* | nd | nd |
| CP.22.688 | Toury | Darmor-*Rlm11* | A | V | - | nd | **+** | - | nd | *AvrLm6-0-A* | *AvrLm11-del* | nd | nd |
| CP.22.689 | Toury | Darmor-*Rlm11* | A | V | - | nd | **+** | - | nd | *AvrLm6-0-A* | *AvrLm11-del* | nd | nd |
| CP.22.1022 | Allenes les Marais | Darmor-*Rlm11* | A | V | A | nd | **+** | - | nd | *AvrLm6-0-A* | *AvrLm11-del* | nd | nd |
| CP.22.1023 | Allenes les Marais | Darmor-*Rlm11* | A | V | A | nd | **+** | - | nd | *AvrLm6-0-A* | *AvrLm11-del* | nd | nd |
| CP.22.1025 | Allenes les Marais | Darmor-*Rlm6* | A | A | - | nd | **+** | **+** | nd | *AvrLm6-0-A* | *AvrLm11-0-A_(876 pb_1)* | nd | nd |

[1], V, Virulent, A, avirulent, H, versatile phenotype depending on replicates

[2] -, lack of amplification; + positive amplification,; +/- weak or variable amplification; nd, (often due to RIP mutations within the primer sequence); not done

[3] del, deleted allele; rip, RIPped allele; dup, duplicated sequence in the geneome; new, dew allele compared to published data. Correspondance between geneotype and assigned phenotype is in Supplementary Table 6

[4] Probable RIP, allele supposedely RIPed due to low amplification and low quality sequence

[5] the numbers between brakets indicate the size of the amplified region, the difference is due to variable numbers of a 17 bp minisatellite sequence before the start codon
